# Spatiotemporal dynamics of lexical and morphosyntactic planning during language production revealed by magnetoencephalography

**DOI:** 10.64898/2026.09.17.752155

**Authors:** Yi Wei, L. Robert Slevc, Christian Brodbeck, Yasmeen Faroqi-Shah

**Affiliations:** Department of Hearing and Speech Sciences, University of Maryland, College Park, MD, United States; Department of Psychology, University of Maryland, College Park, MD, United States; Program in Neuroscience and Cognitive Science, University of Maryland, College Park, MD, United States; Department of Computing and Software, McMaster University, Hamilton, ON, Canada

## Abstract

Fluent sentence production relies on the precise and timely coordination of lexical retrieval, morphosyntactic planning, and motor execution. While a left-dominant fronto-parieto-temporal network has been identified for these subcomponents of sentence production, less is known about the time course of engagement of these regions. One key issue is the extent to which the engagement of neural regions is sequential. The aim of the present study was to delineate spatiotemporal patterns of subcomponents of sentence production, namely lexical retrieval, inflection, and constituent assembly, and identify any lexical category differences (nouns versus verbs) in these processes. Magnetoencephalographic (MEG) responses were measured using a novel picture-icon paradigm in which verbal responses were elicited using object or action pictures followed by an icon that indicated the subcomponent sentence processes: lexical access, inflection (plurals for nouns, tense for verbs), and constituent assembly (adjective + object or pronoun + action). MEG data from twenty native English-speaking right-handed neurotypical adults was analysed for both stimulus-locked and response-locked timelines. We focused on responses in left hemisphere speech-language regions of interest (ROI): precentral gyrus (M1), inferior frontal gyrus (IFG), anterior temporal lobe (ATL), posterior temporal lobe (PTL), and inferior parietal lobe (IPL). The results revealed the following: 1) early and largely simultaneous engagement of all ROIs for lexical access; 2) compared to noun lexical access, verb lexical access is associated with a larger neural response across multiple language ROIs; 3) morphosyntactic planning engaged all ROIs, showing commonalities as well as unique patterns for each morphosyntactic condition; 4) noun morphosyntax vs noun lexical access showed a larger response across multiple ROIs compared to verb morphosyntax vs verb lexical access; 5) M1 was coactivated with other ROIs early in the planning stage, suggesting simultaneity of language and motor planning. The findings support a network model of language in which the syntactic, semantic, and phonological properties of a word are bound to a single anatomically distributed representation.

## 1. Introduction

Sentence production involves conveying thoughts (which are often nonsequential) using words that are linearly sequenced according to grammatical rules, and expressed via articulatory movements. In fact, sentence production is considered one of the most sophisticated and temporally rapid cognitive operations performed by the human brain. Neuroimaging and neuropsychological studies have identified a left-hemisphere dominant fronto-parieto-temporal network of regions involved in sentence production (Faroqi-Shah, et al., 2014; Gleichgerrcht, et al., 2021; Hauptman et al., 2022; Matchin & Hickok, 2020; Morgan et al., 2025; see review by Yeaton, 2025). Research has attempted to identify the functional specialization (if any) of these language network regions for component processes of sentence production but a comprehensive model is yet to emerge. What is particularly unclear is the time course of activation of these neural regions during language production. A better understanding of when neural regions are activated for which component of sentence planning will inform fundamental questions about the neural architecture of language production. In particular, it will help inform how lexical and morphosyntactic planning is implemented and adjudicate between sequential and parallel views of language production.

### 1.1. Lexical and morphosyntactic planning

It is generally agreed that sentence production entails the planning of content (lexical), structural (morphosyntactic), and articulatory (motoric) information (Bock & Levelt, 1994; Dell, 1986; Garrett, 1975; Levelt, 1989; Slevc, 2023; but see Krauska & Lau, 2023). Lexical items are assembled into phrases, and eventually into sentences which convey the entities involved and the relationships between those entities (i.e., grammatical functions indicating who did what to whom) while adhering to the structural properties of the language (e.g., word order, inflectional morphemes, compounding, etc.).

#### 1.1.1. Lexical retrieval

The posterior superior temporal region (posterior superior temporal gyrus, pSTG, and posterior superior temporal sulcus, pSTS) has been associated with phonological representations of words (Indefrey, 2011, Maess et al., 2002; Miozzo et al., 2014). The left anterior temporal lobe (ATL) and the posterior middle temporal gyrus (pMTG) have been associated with word semantics (e.g., Abel et al., 2009; Hocking et al., 2009). A majority of neural studies of lexical retrieval have focused on noun retrieval (e.g., Indefrey, 2011) and generally less is known about verb retrieval. Some studies have reported neural differences between noun and verb retrieval (Bedny & Caramazza, 2011; Bedny et al., 2012; Berlingeri et al., 2008; see meta-analysis by Faroqi-Shah et al., 2018). Unique activation for verbs but not nouns has been found in the left pMTG, inferior frontal gyrus (IFG), and inferior parietal lobe (IPL) (Faroqi-Shah et al., 2018; Shapiro et al., 2006; Thompson et al., 2007). These unique findings have been associated with action concepts and visual motion in the pMTG, morphological encoding in the IFG, and verb argument structure and thematic relations in the IPL (Bedny et al., 2012; Faroqi-Shah et al., 2018; Shapiro, et al., 2006; Thompson et al., 2007). Verb retrieval is considered to be more central to sentence planning because the type of verb selected influences the morphosyntactic structure of a sentence (Bock & Levelt, 1994;Thompson et al., 2007). Thus, verb retrieval and morphosyntactic planning should share more neural similarities than noun retrieval and morphosyntactic planning. However, this interaction between lexical category and morphosyntactic planning has not been systematically examined.

As for the time course of lexical processes, Indefrey (2011) compiled a spatio-temporal map by consolidating findings from different neuroimaging methods. This map suggests that semantic/grammatical information and phonological representations are sequentially activated at 200-275 ms and 275-355 ms in the posterior and middle temporal lobe respectively. However, Mundig, Dubarry, and Alario’s (2016) meta-analysis of MEG data questioned the sequential nature of semantic and phonological (and articulatory) activation, because they found multiple temporal, parietal, and frontal areas to be simultaneously active as early as 100 ms (also see Miozzo, Pulvermuller, and Hauk, 2014).

#### 1.1.2. Morphosyntactic Planning

Morphosyntactic processes have been described using several terms (e.g., unification, MERGE, hierarchical syntax, linearization, etc.). In this study, we will use the terms *inflection* and *constituent assembl*y to refer to two distinct rule-based processes. Inflection is defined as a word-internal process involving the addition of an inflectional suffix to a word stem to create a bimorphemic word. Examples of inflectional processes in English (and the present study) are pluralization of nouns (e.g., tree+s = trees) and simple past tense on verbs (e.g. cook+ed = cooked). Constituent assembly is defined as combining two or more words in a grammatically acceptable manner to result in a phrase or sentence (e.g., the + tree = the tree).

Neuroimaging studies of inflection production (covert and overt) have identified a wide network of bilateral cortical regions, including the mid-inferior frontal lobe, premotor and primary motor regions, posterior temporal lobe (PTL), ATL, and IPL (Beretta et al., 2003; Desai et al., 2006; Hauptman et al., 2022; Kielar et al., 2011; Lee et al., 2018; Marangolo et al., 2006; Sahin et al., 2006; Slioussar et al., 2014; reviewed in Leminen et al., 2019). While far fewer studies have examined noun inflection compared to verb inflection, both lexical categories have shown comparable findings (Beretta et al., 2003; Hauptman et al., 2022; Sahin et al., 2006; Slioussar et al., 2014). Limited information on the time course of inflection production comes from either anatomically restricted intracranial recordings (Lee et al., 2018, Sahin et al., 2009) or MEG (Hauptman et al., 2022). Intracranial stimulation has identified IFG involvement at 320 ms after stimulus presentation (Sahin et al., 2009) and temporoparietal junction involvement around 1000 - 1500 ms prior to articulation (Lee et al., 2018). An MEG study identified a ventral prefrontal and anterior temporal response at 200-300 ms specifically for the addition of -s suffix and a response at 335-400 ms for all inflectional conditions (Hauptman et al., 2022).

Studies of constituent assembly have examined the production of phrases and sentences and have identified a wide cortical network, including the left ventromedial prefrontal cortex, IFG, PTL, ATL, and IPL (Burki & Laganaro, 2014; Del Prato & Pylkkanen, 2014; Giglio et al., 2022; Giglio et al., 2024; Goldman et al., 2023; Habets et al., 2008; Indefrey et al., 2004; Schell et al., 2022; Pylkkanen et al., 2014). The limited neurophysiological evidence suggests that constituent assembly is initiated concurrently with lexical access, starting as early as 180 ms after the stimulus or speaking prompt is presented (Burki & Laganaro, 2014; Habets et al., 2008; Pylkkanen et al., 2014; Stephan et al., 2020).

Among the cortical regions associated with morphosyntax, it has been proposed that the left IFG and PTL are particularly critical as syntactic hubs (e.g., Matchin & Hickok, 2020; Maran et al., 2022; Murphy, 2024). One view is that IFG and PTL have distinct functions, with production tasks showing that IFG is associated with inflection and production more generally, while the PTL is associated with constituent assembly and comprehension more generally (Brennan & Hale, 2019; Burki & Laganaro, 2014; Giglio et al., 2022; Law & Pylkkanen, 2021; Matchin & Wood, 2020; Sahin et al., 2009; Tyler et al., 2005). Another view is that the PTL and IFG interact in a dynamically reverberating network (Giglio et al., 2024). Finally, some authors have questioned the notion of specialized morphosyntactic production regions (Fedorenko et al., 2020). Overall, relatively little is known about the time course of morphosyntactic planning for language production, particularly with anatomical specificity (but see Del Prato & Pylkkanen, 2014; Hauptman et al., 2022; Lee et al., 2018; Sahin et al., 2009). To our knowledge, inflection and constituent assembly have not previously been examined within the same experimental paradigm.

### 1.2. Sequential versus parallel language planning

Hemodynamically-based neuroimaging methods have advanced current understanding of the neuroanatomy of lexical and morphosyntactic planning. However, there is less clarity on the relative time course of these processes and the extent to which identified neural regions are engaged in a sequential or parallel manner. While conventional models assume that meaning, grammar, and different aspects of sound are activated in a roughly sequential manner (either in a cascading or interactive manner) (e.g., Levelt et al., 1999; Dell 1986), more recent views favor parallelism (e.g., Pickering & Strijkers, 2024).

Neurophysiological methods which provide millisecond level temporal resolution, such as electrocorticography (ECoG), electroencephalography (EEG), and magnetoencephalography (MEG) provide valuable insights into the time course of linguistic operations. For example, Sahin et al. (2009) tracked activity in the left IFG using ECoG when participants produced uninflected and inflected nouns and verbs. They found a distinct time course of neuronal activity for lexical (∼200 ms), inflectional (∼320 ms), and phonological (∼450 ms) processing in the IFG. This time course was similar for nouns and verbs. The findings from Sahin et al. (2009) thus support a sequential pattern of activation in the IFG (see also Carota et al., 2022; Indefrey & Levelt, 2004). In contrast, a MEG study by Miozzo et al. (2014) found evidence of early and parallel activation (∼150 ms) of semantic and phonological information in fronto-temporal and middle temporal regions respectively. Participants engaged in a picture naming task in which the semantic and phonological features of the pictures were orthogonally manipulated. Miozzo et al. (2014) referred to this early and parallel activation of semantic and phonological features as the *simultaneous ignition hypothesis*.

Early or simultaneous ignition has also been recently proposed for sentence planning based on a review of neurophysiological findings (Pickering & Strijkers, 2024). Pickering and Strijkers proposed that the semantic, syntactic, and phonological properties of a word are bound into a single representation (see also Matchin & Hickok, 2020). During language production, this representation is activated as a whole and all linguistic properties are available simultaneously. This simultaneous ignition is proposed to be followed by more controlled, sequential processing based on intentions and task demands. This latter phase is referred to as *reverberation* (Fairs et al., 2021; Pickering & Strijkers, 2024; Strijkers & Costa, 2016).

Note, however, that most current evidence for the time course of language operations is based on single word naming and sentence comprehension studies (Fairs et al., 2021; Miozzo et al., 2014). There is relatively little direct research on the time course of sentence planning for production, with the exception of a handful of studies involving the production of inflectional morphology (tree >>trees), two-word phrases (red tree), and narrative language (Burki & Laganaro, 2014; Del Prato & Pylkkännen, 2015; Hauptman, et al., 2022; Morgan et al., 2025). Although these studies have examined some element(s) of sentence planning, a common confound in these studies has been isolating motor planning from lexical and morphosyntactic planning. This is an important confound because there is evidence that motor planning occurs very early and may interact with sentence planning (Carota et al., 2023; Pickering & Garrod, 2013).

Given these unknowns, the goal of present study is to examine the time course of neural activity in the left fronto-parieto-temporal language network for lexical and morphosyntactic planning for production using MEG. The broader goal of the present study is to fine-tune the neural architecture of language production and inform the debate between sequential and parallel encoding of language and motor processes. Parallel encoding (i.e., simultaneous ignition) of sentence components will be supported if we find simultaneous and early activation of fronto-parieto-temporal regions for the different experimental conditions. In contrast, temporally separated activations across experimental conditions would support a sequential view of sentence planning.

### 1.3. The present study

The appraisal of the literature on the spatiotemporal architecture of sentence production provides a tentative time course for some, but not all, components of sentence production. Few studies have examined multiple components of sentence production within the same experimental paradigm and participant group, and fewer studies have examined how these operations differ for each lexical category (nouns vs verbs). To address these gaps, the goal of the present study is to examine the time course of neural activity in the left fronto-parieto-temporal language network for lexical and morphosyntactic processes underpinning sentence production using MEG. The ultimate goal is to fine-tune the neural architecture of sentence production and inform the debate between sequential and parallel encoding of language and motor planning processes.

In the present study, we investigated the time course of sentence production components using an overt, rather than covert, speech task for increased ecological validity. We specifically examined lexical access (single word production), inflection, and constituent assembly across nouns and verbs. Inflection was investigated in the context of plurals for nouns (*tree+s*) and past tense for verbs (*cook+ed*). Constituent assembly was engaged in nouns by eliciting modifier noun phrases (*a + blue + tree*) and in verbs using short sentences with past and future tense (*She + cooked, She + will + cook*).

To elicit these utterances, we designed a novel picture-icon paradigm in which the picture guided lexical access and the icon indicated the type of utterance that had to be produced (Figure 1A). MEG studies involving overt verbal production of sentences pose methodological challenges which are enumerated here, along with how our study was designed to address these challenges. The first issue is the complexity of the stimuli used to elicit words from different lexical categories (nouns, verbs) because action images used to elicit verbs can elicit a variety of labels (Szekely et al., 2005). We addressed this challenge by selecting noun and verb pictures that had comparable and high name agreement (Szekely et al., 2005). Differences in stimulus complexity across experimental conditions have confounded some prior studies, particularly when single pictures are used to elicit words and contrasted with a series of pictures or complex scenes to elicit phrases/sentences (e.g., Burki & Laganaro, 2014; Goldman et al., 2023; Morgan et al., 2025). To address this confound, we elicited responses of various lengths by pairing the single pictures with a unique *condition* icon that indicated the type of response. Examples of verb and noun pictures and the various *condition icons* are illustrated in Figure 1B. The experimental conditions are not parallel between nouns and verbs and are dictated by the morphosyntactic structure of the English language. Having unique condition icons also allowed us to randomize the presentation sequence rather than using a blocked design (e.g., Burki & Laganaro, 2014; Goldman et al., 2023). In blocked designs, participants can create a syntactic frame (e.g. color+noun) and apply it to each trial within the block, and may thus not fully engage in syntactic planning for each trial. Similarly, to engage a deeper level of lexical planning, we used a larger set of pictures (N=100) compared to prior studies that used a small set of 3-6 pictures for the entire experiment (e.g., Goldman et al., 2023; Hauptman et al., 2022).

**Figure 1.**
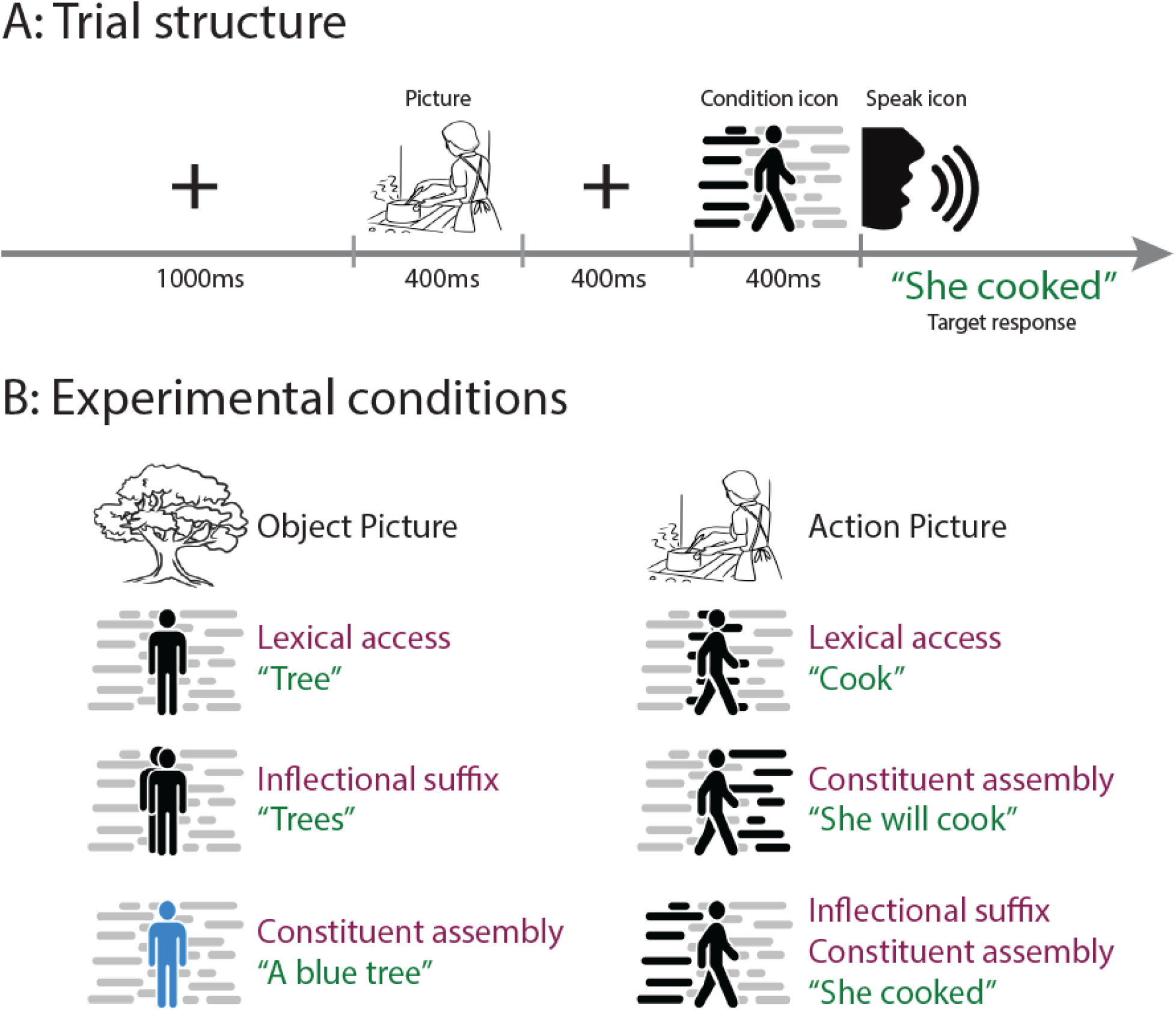
(A) The sequence and timing of each experimental trial, illustrating the stimuli for the target response “She cooked”; (B) The six experimental conditions, three each for object and action pictures, and the icons used to elicit each expected verbal response.

As shown in Figure 1A, the *condition* icon was presented 800 ms after the picture. The advantage of this temporal spacing was that it allowed us to separately examine neural responses to lexical access (0-800 ms) and morphosyntactic planning (> 800 ms). Separating the picture and condition icons also limited oculo-muscular artifacts from eye saccades which would occur if the picture and condition icon were presented simultaneously on different parts of the visual display. In order to minimize concurrent motor planning and the introduction of oro-muscular artifacts, participants were asked to remain silent until a *speak icon* was presented 400 ms after the condition icon (Figure 1A). The 400 ms spacing between the *condition* and *speak* icons allowed sufficient time for visual recognition of the icon meaning, while limiting the time available for pre-articulatory motor planning. Additionally, we used electromyography (EMG) of facial muscles to control for speech-related artifacts from the MEG signal. Further details of the experimental design are provided in the Materials and Methods section 2.2. Given the variability in production latencies across word categories, morphosyntactic conditions, and participants, neural responses that are time-locked to stimulus onset may washout activity for articulatory planning. Hence we conducted both stimulus-locked and speech onset-locked analyses (Salmelin et al., 1994).

## 2. Materials and Methods

The Organization for Human Brain Mapping’s (OHBM) best practice guidelines for reproducibility of MEG research were followed in reporting study methods (Pernet et al., 2020).

### 2.1. Participants

Participants were recruited from the local university community through emails, flyers, and word-of-mouth. All participants provided written informed consent in compliance with the ethical guidelines of the University of Maryland, and were paid for participation. Twenty-nine participants were initially recruited. Data from nine participants was excluded for the following reasons: unusable data due to corrupted speech recordings (N=1), non-native speaker of English (N=5), and not right-handed (N=2 left-handed, N=1 ambidextrous).

MEG data of 20 right-handed neurotypical native English-speaking adults were analyzed and are reported here (12 females, age: M (SD) = 22.05 (3.56)). The participants had no neurological or psychiatric diagnoses and had corrected or normal vision and hearing, as per self-report. There is no straightforward way to estimate statistical power a priori for the source space free orientation vector analyses used in the present study, however the sample size of the present study is comparable with other MEG studies of language production (Del Prato & Pylkkanen, 2014; Miozzo et al., 2015).

### 2.2. Stimuli and experimental conditions

Fifty object and fifty action pictures were selected from the International Picture Naming Project (IPNP, Szekely et al., 2005) to elicit nouns and verbs respectively. The IPNP database provides psycholinguistic norms, including name agreement, and speech onset latencies for a large set of object and action pictures. The main metrics by which object and action pictures were matched was IPNP name agreement (proportion name agreement: Nouns M (SD) = .91 (.08), Verbs M (SD) = .93 (.07), t=1.45, p=.15) and number of syllables. All word stems were monosyllabic. Word frequency and age of acquisition did not differ across nouns and verbs (part of speech log frequency/million from Brysbaert & New, 2009: Nouns M (SD) = 2.94 (.64), Verbs M (SD) = 3.05 (.78), t=0.73, p=.47; age of acquisition from Fenson et al., 2007: Nouns M (SD) = 2.08 (.92), Verbs M (SD) = 2.28 (.90), t=1.09, p=.28). Naming latencies in the IPNP database were slower for action pictures than object pictures (Nouns M (SD) = 949.67 ms (170.62), Verbs M (SD) = 1065 ms (169.32), t=3.41, p < .001). The IPNP uses image file size as a proxy for visual complexity, and action pictures were visually more complex than object pictures (Nouns M (SD) = 14486.16 KB (5913.72), Verbs M (SD) = 22482 KB (7,800), t=5.78, p < .001).

The *condition* icons were specifically created for this study, illustrating a standing stick figure for noun trials and a walking stick figure for verb trials, as shown in Figure 1B. There were three noun conditions and three verb conditions, resulting in a total of 300 trials (2 word categories x 3 conditions x 50 pictures). The location of solid black horizontal lines in the verb icons indicated past versus future tense.

### 2.3. Study procedure and data acquisition

The data reported here were part of a longer experimental session that included: resting state, the present study’s trial sequence, a different trial sequence (condition icon preceding the picture)^1^, a non-speech condition (saying “blah blah”), and an “empty room” recording. The full session lasted about 1.5 hours.

After providing informed consent, participants were introduced to the *condition* icons, and the corresponding expected verbal response. Participants received 12 practice trials (6 conditions x 2 trials) using a different set of pictures not used in the main experiment. They were allowed to review these 12 practice trials until they felt comfortable performing the experimental task. Head shape digitization was performed next using a Polhemus 3SPACE FASTRAK system (Polhemus, 2012) with 3 fiducial points and 5 marker positions. Five marker coils were placed on the participant’s head at these marker locations to measure head position relative to the MEG sensors at the beginning and the end of each recording session. On average, the distance between pre- and post-experiment marker positions was 4.59 mm, although 3 participants showed movements greater than 10 mm (12.36, 10.04, 14.5 mm). In order to detect and remove speech-related orofacial movement artifacts resulting from the MEG signal, three EMG sensors were used (iWorx Systems Inc). Based on the findings of Abbasi et al. (2021), one pair of EMG electrodes were placed at the location of the right orbicularis oris muscle, and a ground electrode was placed on the right shoulder (see Supplementary Figure S1). Participants were then familiarized with the picture stimuli and their target names. Participants were allowed to repeat the familiarization process until they felt comfortable naming all the pictures.

Following this, MEG data were acquired on a 157 axial gradiometer whole head MEG system (KIT, Kanazawa, Japan), with an online 200 Hz low pass filter and a 60 Hz notch filter at a sampling rate of 1 kHz. Participants lay in supine position and stimuli were presented on a screen located about 31 inches above the participant’s face. Participants’ verbal responses were recorded via a microphone (Optoacoustics Optimic MEG Fiberoptic microphone) inside the magnetically shielded room. A trigger code was sent to MEG, EMG, and audio recording simultaneously at the beginning of each trial. The hardware set up is illustrated in Supplementary Figure S2.

The experiment was programmed in Presentation version 16.3 (Neurobehavioral Systems, 2022). As illustrated in Figure 1A, each trial began with a fixation cross for 1000 ms, followed by an object or action picture for 400 ms, then another fixation cross for 400 ms, followed by a *condition* icon for 400 ms, then the *speak* icon^2^. Participants were instructed to speak only after seeing the *speak* icon. The total stimulus presentation time, which is time between the appearance of the first fixation cross and the speak icon, was 2.2 seconds. The next trial was initiated manually by the experimenter, who clicked the mouse on the stimulus presentation computer. The experimental trials were presented in a pseudorandomized sequence across eight blocks of 50 trials each^3^, such that the same picture-condition combination was not repeated on successive trials.

### 2.4. Behavioral Word onset time detection

Participants’ verbal responses were manually transcribed. Trials with incorrect responses (including self corrections) were excluded from behavioral and MEG data analysis. The speech onset time for each trial was defined as the time between the presentation of the speak icon and the initiation of the verbal response. Onset times were estimated using Montreal forced aligner (v3.1.2, McAuliffe et al., 2017), using the acoustic model *english_mfa v3.1.0*. To correct for errors generated by the Montreal forced aligner due to noise in the audio recordings, trials with a speech onset time faster than 300 ms were flagged (M (SD) = 13.95 (11.74)) and manually timed using Praat (Boersma & Weenik, 2025).

### 2.5. Data preprocessing

MEG data was pre-processed with mne-python version 1.8.0 (Gramfort et al., 2014) and Eelbrain version 0.40b1 (Brodbeck et al., 2019). The code for processing can be accessed at INSERT OSF OR GITHUB LINK. Flat channels were automatically detected and excluded, after which temporal signal space separation was applied to remove artifacts (Taulu & Simola, 2006). The data were then band-pass filtered between 1 and 40 Hz using a zero-phase FIR filter with default MNE-Python settings. Independent component analysis (ICA) was then applied. Components corresponding to blinks, heartbeats, and motion artifacts were removed with visual inspection. A trial was defined as the total stimuli presentation time + two seconds after the onset of the *speak* icon. Data was epoched 1) from 100 ms before the trial onset time to the end of the trial for the stimulus-locked analysis, and 2) from 1 second before the speech onset time to 1 second after the speech onset time for the response-locked analysis.

To remove articulation related muscle artifacts, we first epoched the EMG data the same way as the MEG data. Then we calculated the correlation between the EMG averaged epoch data and each of the first twenty ICA components’ averaged epoch data. Any ICA components that had a correlation coefficient higher than 0.4 were excluded as arising from speech movement artifacts. After all ICA components were removed (M (SD) = 6.75 (1.68)), we visually inspected the time series and topography of the averaged, epoched MEG data for each participant and did not identify any remaining artifacts. After pre-processing, data were epoched as described above and averaged for source localization without baseline correction.

### 2.6. Source localization

The ‘fsaverage’ brain model (Bruce et al., 1999) was co-registered to each participant’s head, using the previously obtained digitized head shapes, via 3-parameter MRI scaling. First, the three fiducial points were used for an approximate alignment (with nasion weighted at 5 and other points at 1), followed by a fit minimizing the distance from all head shape points to the MRI scalp surface using the Iterative Closest Point (ICP) algorithm, and then the three fiducials were used for a last alignment again. A surface based source space was first created on the ‘fsaverage’ brain white matter surface with a recursively subdivided icosahedron at 4 steps. This source space was then scaled to fit each participant’s head shape using the parameters derived above. These source spaces were then used to compute free orientation lead-field matrices by placing 3 orthogonal virtual current dipoles on each of the grid points. The computed lead field matrices contained contributions from 2562 current dipole triplets per hemisphere. MEG signal in the source space was estimated using minimum-norm estimate (MNE) inverse solutions, with regularization corresponding to an assumed signal-to-noise ratio (SNR) of 3 and no depth weighting, returning estimates of the three orthogonal components of the current vector at each source location. A free vector orientation was used for the inverse solution instead of a fixed orientation (i.e., perpendicular to the cortical surface) to accommodate differences in current direction across participants that arise from differences in neuroanatomy not captured by the template brain.

### 2.7. Statistical analyses

#### 2.7.1. Speech Onset Times

Separate regression models were built for noun and verb conditions using *lmer* (Bates et al., 2015) from the *lme4* package (version 1.1-37, Kuznetsova et al., 2017) in Rstudio (version 2024.12.1 with R version 4.5.0). For both regression models, condition was entered as a fixed effect (noun model: noun naming, noun plural, noun phrase; verb model: verb naming, verb past, verb future). Random intercepts and random slopes for condition were included for subjects. Post-hoc pairwise comparisons using Tukey-adjusted *emmeans* (Lenth & Piaskowski, 2025) were used if significant main effects were identified in the model.

#### 2.7.2. MEG Data

Vector source current estimates were analyzed using mass-univariate permutation tests based on the Hotelling-*T*^2^ statistic (Das et al., 2020). The *T*^2^ statistic is analogous to a *t*-value, but summarizes the 3-dimensional vector at a given source location with a value that incorporates consistency across participants in direction and magnitude. For each test, a *T^2^*map was computed across the whole brain. This map was processed with threshold-free cluster enhancement to emphasize signal peaks that extended in space and time (Brodbeck et al., 2018). To control for family-wise error, a null distribution of max *T*^2^ across space and time was estimated based on 10,000 permutations of the data.

#### 2.7.3. Source Localization

Source current analysis first focused on brain activity associated with lexical retrieval separately in each of the two lexical categories. This was done using separate one-sample vector field *T*^2^ tests for the noun naming and verb naming conditions. These tests reveal source currents in space and time with a non-random current direction across subjects. Responses associated with noun and verb naming were directly compared using a paired *T*^2^ vector field test. Brain activity associated with specific morphosyntactic operations was isolated by comparing with the respective lexical category response. Within the noun category, two comparisons were used to examine noun morphosyntax: 1) the noun plural versus noun naming contrast allowed identification of the spatiotemporal response associated with noun inflection. 2) the noun phrase versus noun naming contrast identified the spatiotemporal response to noun constituent assembly. For verbs 1) the future tense versus verb naming contrast identified the response to constituent assembly, and 2) the past tense versus verb naming contrast identified the response to inflection and constituent assembly. For all comparisons, both stimulus-locked (from picture onset until 2 seconds after speak icon) and speech onset-locked (1 second prior to and 1 second following speak onset time) analyses were conducted.

Both the one-sample and paired *T*^2^ vector field tests were applied across the whole cortical region using aparc.a2009s parcellation (Destrieux et al., 2010). Following this whole brain statistical test, differences between experimental conditions were examined in specific regions of interest (ROI), illustrated in Figure 2: primary motor cortex (M1, G_precentral-lh, 60 voxels), Inferior frontal gyrus (IFG, consisted of G_front_inf-Opercular-lh, G_front_inf-Triangul-lh, G_front_inf-Orbital-lh, 46 voxels), inferior parietal lobe (IPL; supramarginal: G_pariet_inf-Supramar-lh, angular gyrus: G_pariet_inf-Angular-lh, 111 voxels), anterior temporal lobe (ATL, defined by the anterior ⅖ th region of the temporal lobe, 71 voxels), and posterior temporal region, defined by the posterior ⅗ th of the temporal lobe (PTL, 123 voxels). The temporal lobe includes the G_temp_sup-Lateral-lh, S_temporal_sup-lh and G_temporal_middle-lh. Aside from the language ROIs, the primary visual cortex region (V1: S_calcarine-lh, Pole_occipital-lh, 82 voxels) was also selected to examine any potential differences in noun and verb image processing.

**Figure 2.**
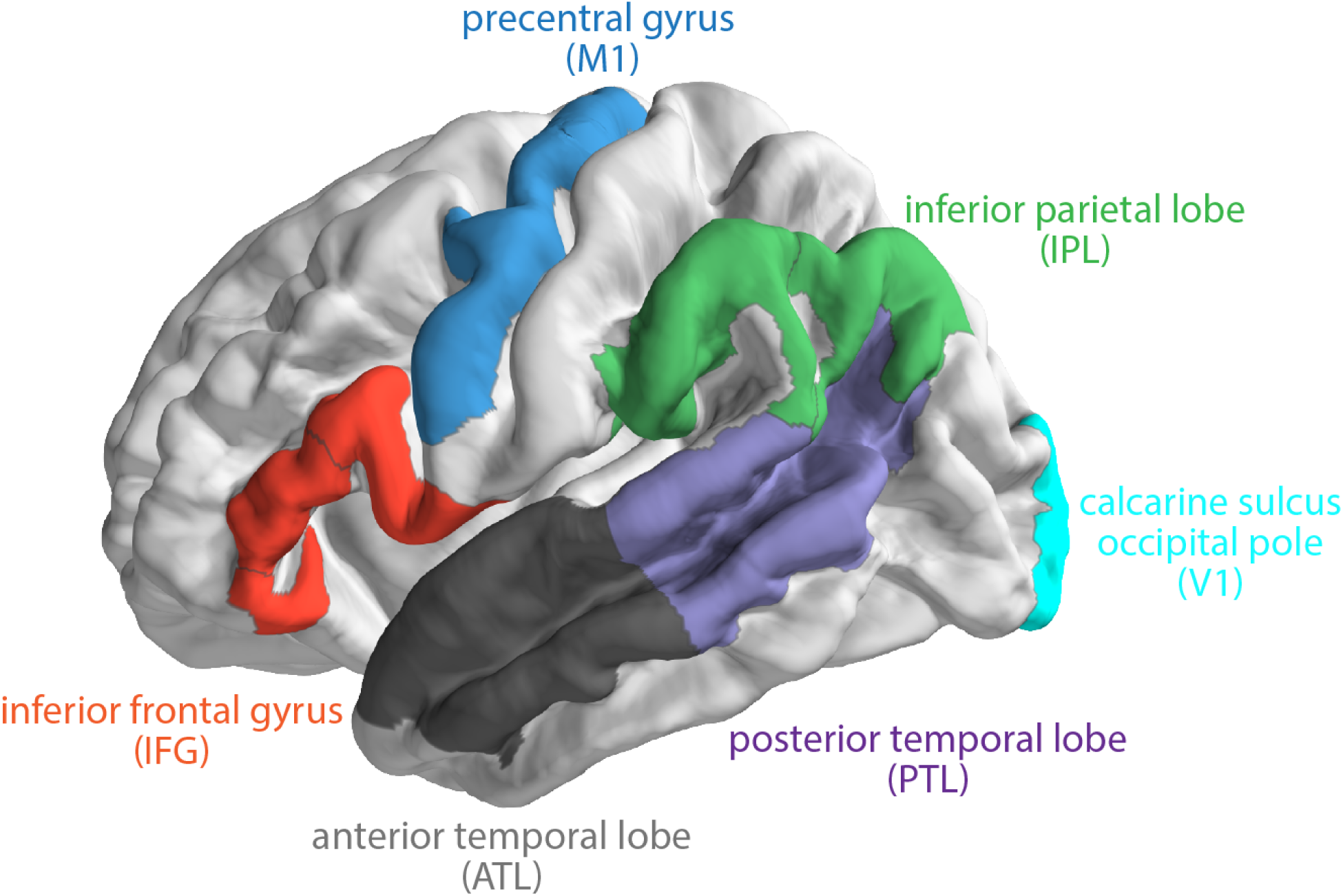
The regions of interest (ROI) used in this study, which are based on Destrieux et al. (2010) parcellation of brain regions. The anatomical labels included in each ROI are given in Section 2.7.3.

For each of the comparisons, we report the findings when 1) more than 30% of voxels within an ROI were significantly different between the conditions, 2) this difference lasted for at least 10 ms, and 3) the averaged *T*^2^ over these significant voxels was larger than 4 at every time point. All significant findings (with more than 30% of voxels within an ROI being significantly different between the conditions) are reported in Supplementary Tables S1-S3. Since the Hotelling *T^2^*-test indicates that vectors are significantly different between comparisons, but not the strength of the activation amplitude, we also calculated the root mean square (RMS) of each condition at any time point when the Hotelling *T^2^*-test showed a significant difference between the conditions.

## 3. Results

### 3.1. Speech onset times

Speech onset times in seconds (Mean (SD)) across the conditions were as follows: 0.6 (0.33) for noun naming, 0.63 (0.30) for noun inflection, 0.62 (0.26) for noun constituent assembly, 0.66 (0.38) for verb naming, 0.77 (0.42) for verb constituent assembly (future) and 0.79 (0.46) for verb inflection + constituent assembly (past) (illustrated in Supplementary Figure S3). As described in the statistical analysis section, separate regression models were built for noun and verb conditions. Results showed no significant main effect of the noun conditions but a significant effect of verb conditions, F(2, 17.99) = 7.26, p = 0.005. Post-hoc pairwise comparisons revealed that verb naming times were significantly faster than both the verb constituent assembly (future) condition (estimate = -0.01, SE - 0.03, p = 0.003), and the verb inflection + constituent assembly (past) condition (estimate = -0.13, SE - 0.04, p = 0.018). The two verb morphosyntax conditions did not differ from each other (estimate = -0.02, SE = 0.03, p = 0.67).

### 3.2. Stimulus locked analyses

The left hemisphere responses that met our reporting criteria (contains at least 30% ROI voxels for at least 10 ms and the average *T*^2^ of at least 4) are listed in Tables 1 through 7. All significant results (contains at least 30% ROI of voxels) are provided in Supplementary Tables S1 - S3. Figures 3 – 10 illustrate the time course and associated spatial distribution of each contrast.

**Figure 3.**
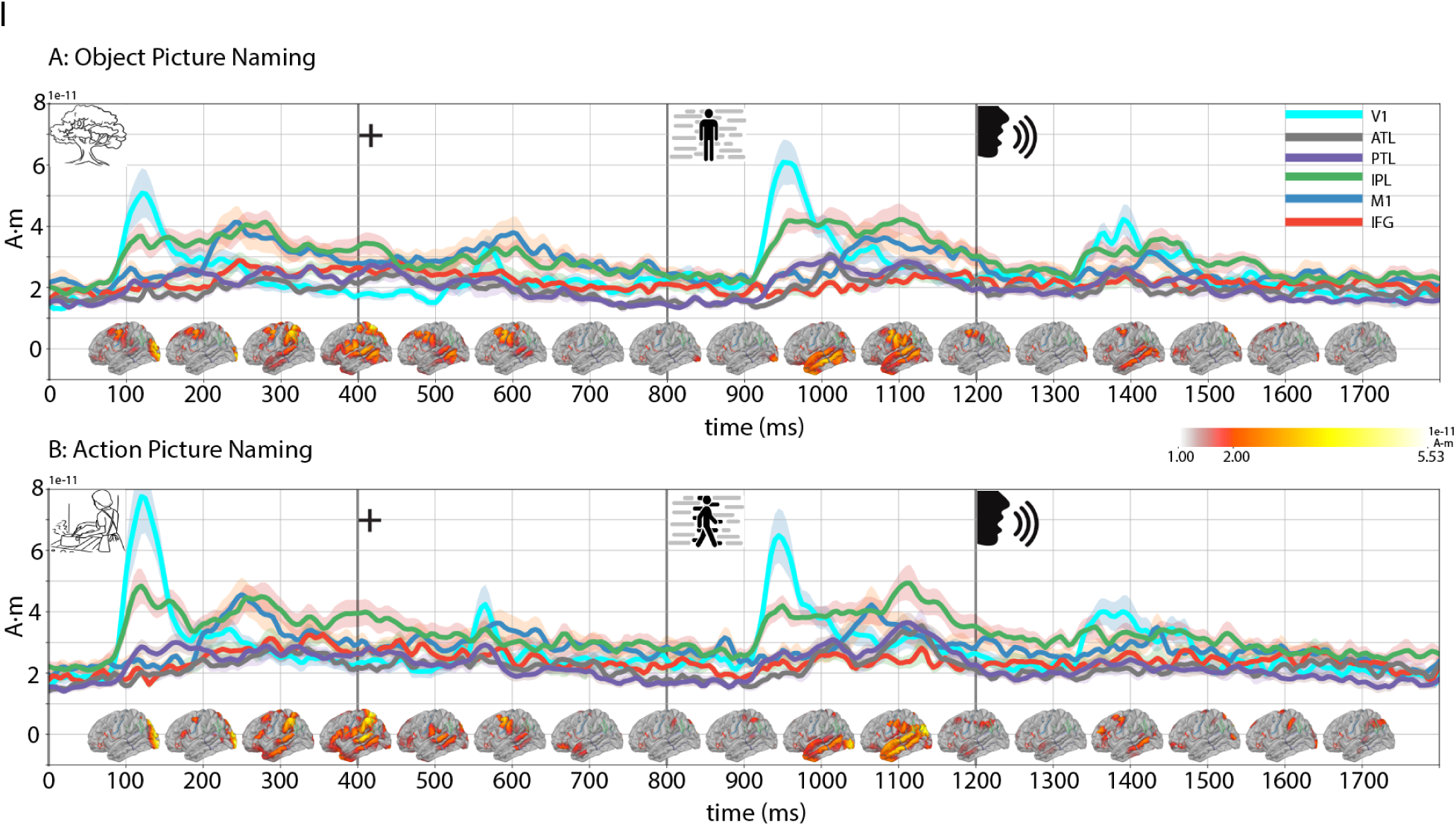
MEG responses in the noun (A) and verb naming conditions (B). Line traces indicate the group-averaged RMS of the source vector magnitude in regions of interest (cf. Figure 2). Shading indicates the between-subject standard error. Anatomical plots at the bottom of each panel show sources with significant responses in 100 ms time bins (centered on 100 ms, 200 ms, etc.; *p* ≤ .05, corrected for the whole brain and time course). Line traces representing ROIs are color coded; Visual cortex is light blue; ATL is gray, PTL is purple, IPL is green, M1 is dark blue, and IFG is red.

#### 3.2.1. Lexical retrieval

Figure 3 shows group average MEG responses and results from one-sample vector field *T*^2^ tests for noun naming and verb naming. A complete list of significant results are provided in Supplementary Table S1. Visual cortex responses occurred 120-200 ms after the presentation of any image or icon during the trial. Note that for V1, the higher vector magnitude in the verb naming condition is consistent with the higher visual complexity of the action images (section 2.2 Stimuli). Visual cortex responses will not be discussed further.

Within 100 ms of picture presentation both noun and verb naming showed responses in all five ROIs. The IPL had the earliest and most sustained response with the largest RMS source vector magnitude across most time zones (see Figure 3). Response strengths of other ROIs varied by time range, with PTL showing an increased RMS at 100-200 ms, and M1 showing a larger RMS source magnitude around 250-350 ms and 550-650 ms relative to other ROIs. A similar ROI response pattern re-appeared after the *condition* icon, with stronger RMS magnitude of the IPL, PTL, and M1 (900-1100 ms after picture onset).

Figure 4 and Table 1 show the results of a paired vector field *T*^2^ test directly comparing noun and verb naming (see Supplementary Table S2 for all significant contrasts). Compared to noun naming (i.e., verb naming > noun naming), verb naming was characterized by stronger activations in four out of five ROIs (except for M1) in the 130-800 ms time range. This stronger verb response first emerged in the IPL and PTL ∼110 ms after the picture onset. In the reverse contrast (i.e., noun naming > verb naming), noun naming showed a stronger response than verb naming at a few points in three of the five ROIs (ATL, PTL, and IFG).

**Figure 4.**
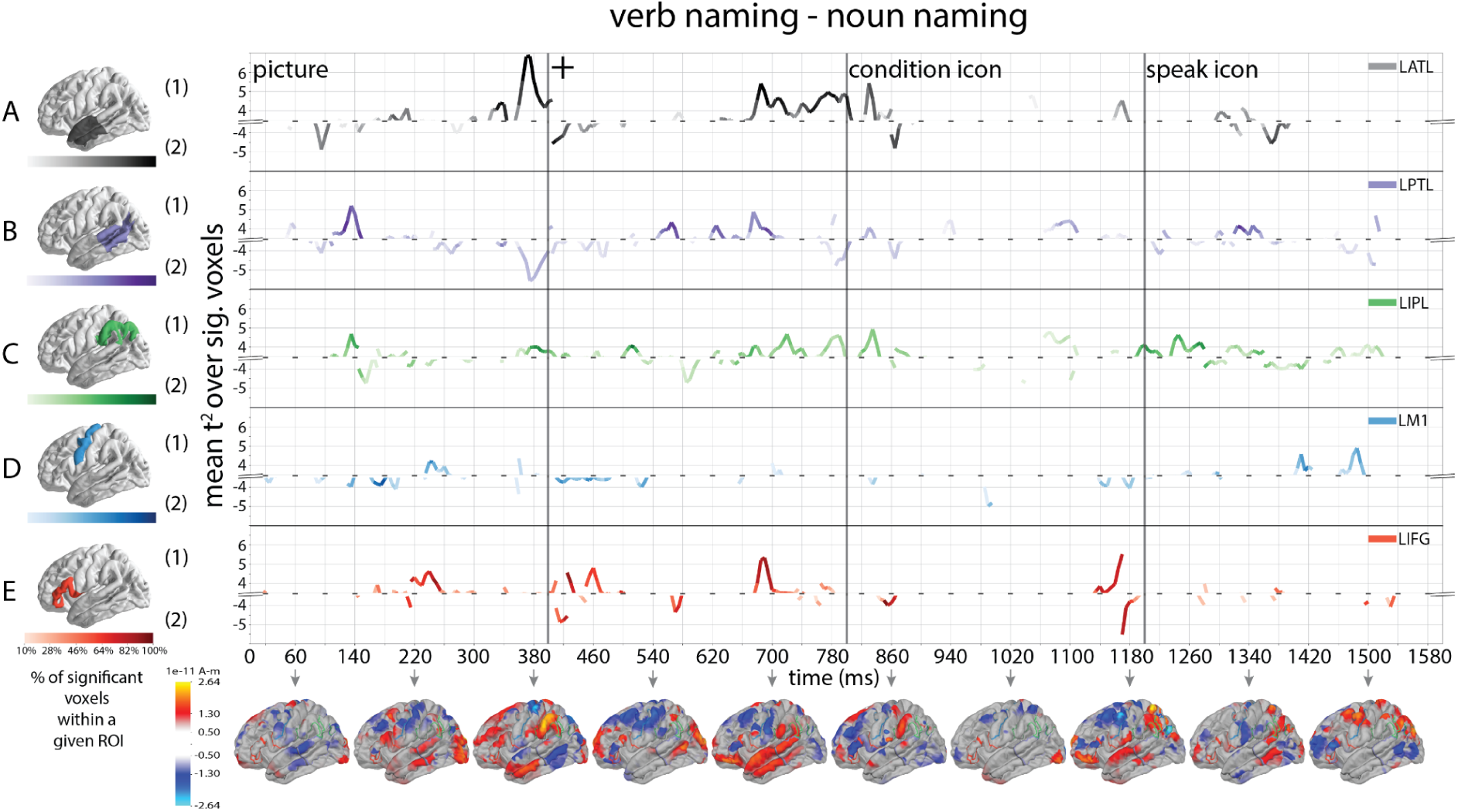
Direct comparison between verb and noun naming by region of interest (ROI). Line traces indicate when Hotelling *T^2^*-test showed a significant difference between the two conditions. Positive values show greater verb response (verb naming > noun naming) by ROI, indicated by greater RMS of source vector magnitude. Negative values show greater noun response (noun naming > verb naming) by ROI (e.g., A(1) shows greater verb response in ATL region, and A(2) shows greater noun response in ATL region). ROIs are color coded, ATL is gray, PTL is purple, IPL is green, M1 is dark blue, and IFG is red. The color saturation of the line traces reflects the percentage of significant voxels (i.e., the darker the color, the greater the percentage of voxels in an ROI that showed significant differences between the two conditions). The anatomical plots (bottom row) show significant response differences between the two conditions across the whole brain (only the left hemisphere is shown, *p* < .05 corrected). The difference values were computed by first taking the norm of the difference vector between the two conditions at each source, and then assigning a sign to each source based on which of the two conditions’ current vector had the larger norm. Lastly, each anatomical plot represents an 80 ms average bin centered at the time indicated by the gray arrow. Red hue reflects greater verb response and blue hue reflects greater noun response.

**Table 1.**
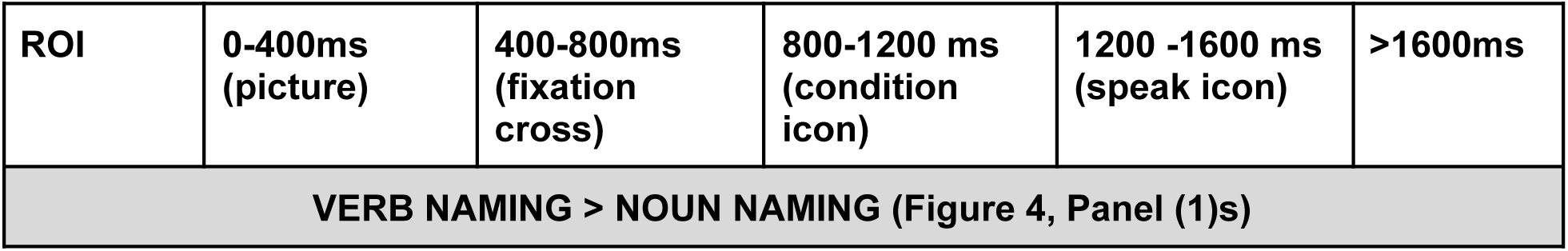
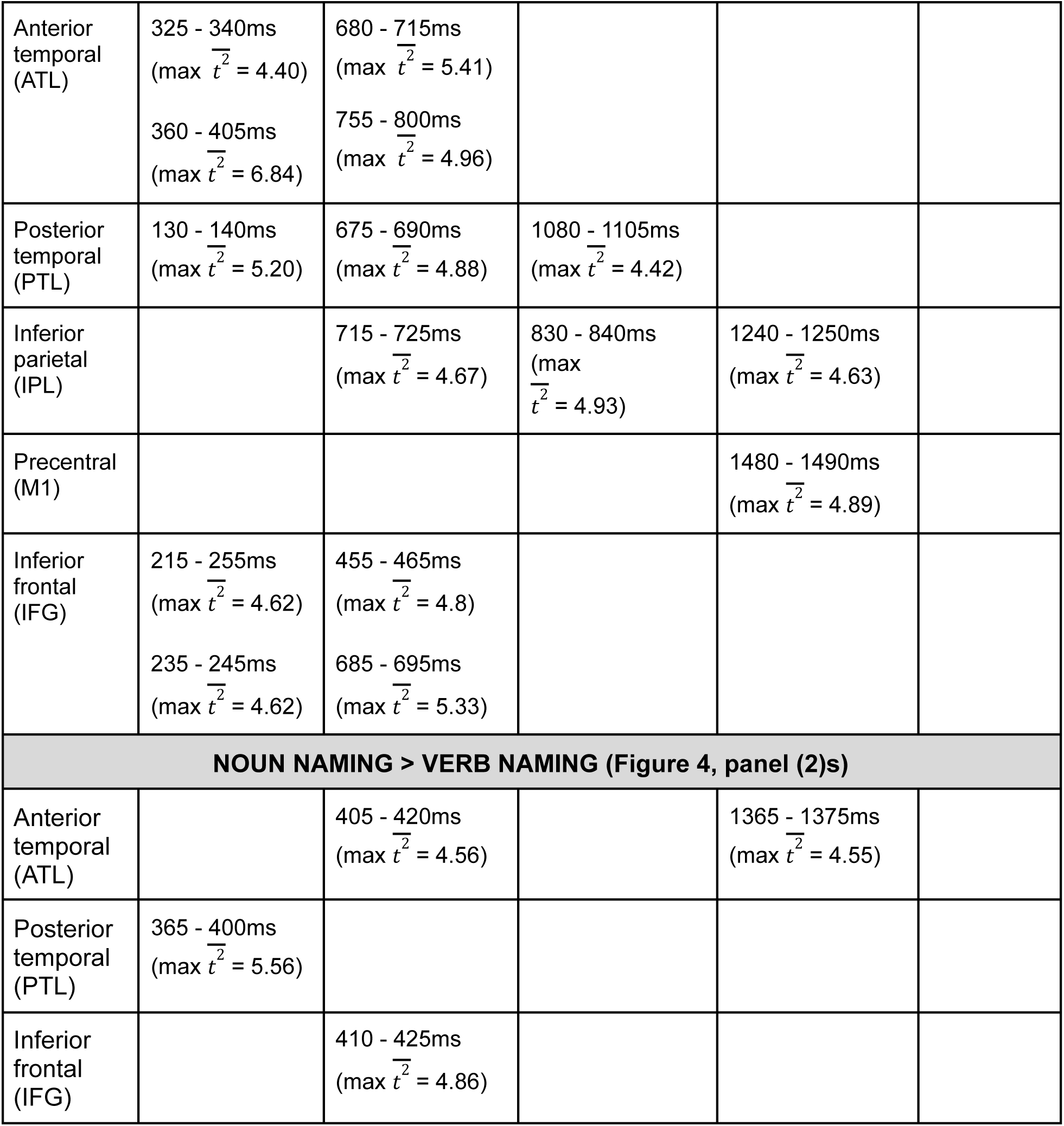
Significant differences between verb and noun naming contrasts in the five left hemisphere regions of interest (ROI) across the experimental trial.

| ROI | 0-400ms<br>(picture) | 400-800ms<br>(fixation<br>cross) | 800-1200 ms<br>(condition<br>icon) | 1200 -1600 ms<br>(speak icon) | >1600ms |
| --- | --- | --- | --- | --- | --- |
| **VERB NAMING > NOUN NAMING (Figure 4, Panel (1)s)** |  |  |  |  |  |
| Anterior temporal (ATL) | 325 - 340ms<br>(max $\overline{t^2} = 4.40$)<br>360 - 405ms<br>(max $\overline{t^2} = 6.84$) | 680 - 715ms<br>(max $\overline{t^2} = 5.41$)<br>755 - 800ms<br>(max $\overline{t^2} = 4.96$) | | | |
| Posterior temporal (PTL) | 130 - 140ms<br>(max $\overline{t^2} = 5.20$) | 675 - 690ms<br>(max $\overline{t^2} = 4.88$) | 1080 - 1105ms<br>(max $\overline{t^2} = 4.42$) | | |
| Inferior parietal (IPL) | | 715 - 725ms<br>(max $\overline{t^2} = 4.67$) | 830 - 840ms<br>(max $\overline{t^2} = 4.93$) | 1240 - 1250ms<br>(max $\overline{t^2} = 4.63$) | |
| Precentral (M1) | | | | 1480 - 1490ms<br>(max $\overline{t^2} = 4.89$) | |
| Inferior frontal (IFG) | 215 - 255ms<br>(max $\overline{t^2} = 4.62$)<br>235 - 245ms<br>(max $\overline{t^2} = 4.62$) | 455 - 465ms<br>(max $\overline{t^2} = 4.8$)<br>685 - 695ms<br>(max $\overline{t^2} = 5.33$) | | | |
| **NOUN NAMING > VERB NAMING (Figure 4, panel (2)s)** |  |  |  |  |  |
| Anterior temporal (ATL) | | 405 - 420ms<br>(max $\overline{t^2} = 4.56$) | | 1365 - 1375ms<br>(max $\overline{t^2} = 4.55$) | |
| Posterior temporal (PTL) | 365 - 400ms<br>(max $\overline{t^2} = 5.56$) | | | | |
| Inferior frontal (IFG) | | 410 - 425ms<br>(max $\overline{t^2} = 4.86$) | | | |

#### 3.2.2. Morphosyntactic planning

Noun and verb naming served as the baseline condition for noun and verb morphosyntax, respectively. Given that all three noun conditions were identical for the first 800 ms (400 ms of picture presentation and 400 ms of fixation cross), whole brain analyses unsurprisingly showed no significant differences in the five ROIs between any noun contrasts for the first 800 ms. The same was found for both verb contrasts. Therefore, all figures and tables are presented starting from 800 ms, and all the timing of reported results here and in the discussion section is with reference to the *condition icon* onset time.

##### 3.2.2.1. Noun morphosyntax

Figure 5 illustrates the results of the noun inflection contrast (plural) relative to noun naming, which were assessed using paired vector field *T*^2^ test (see also Table 2 and Supplemental Table S3). Relative to the presentation of the *condition* icon, noun inflection (noun inflection > noun naming) was associated with two phases of neural activity, one starting around 400 ms and engaging M1, IFG, IPL, and ATL, and another around 600 ms re-engaging the IPL, IFG and engaging the PTL (Figure 5, panel (1)s)^4^. Noun naming relative to noun inflection engaged the M1 at 775 ms following the condition icon (Figure 5, panel (2)s).

**Figure 5.**
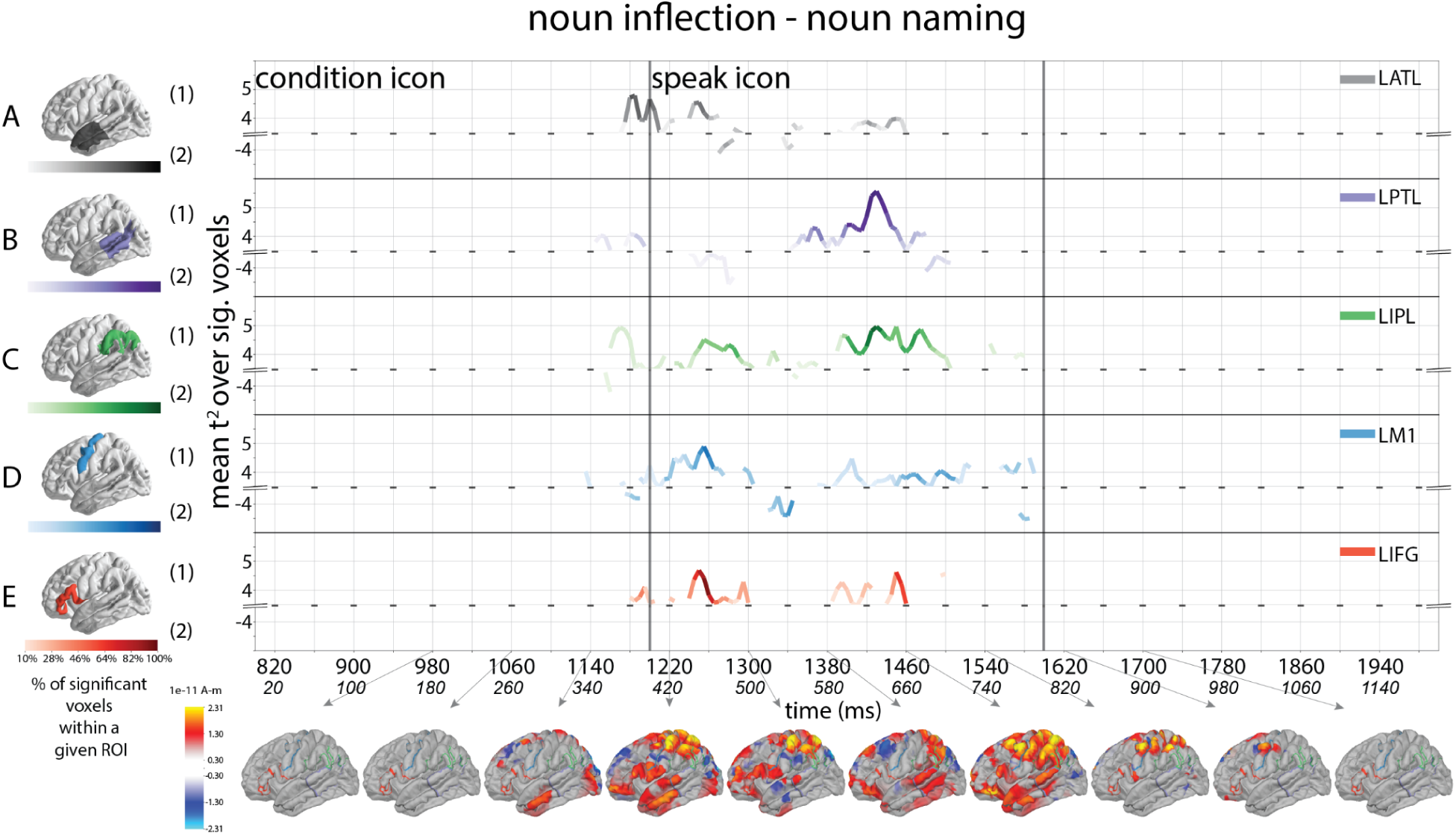
Results of direct comparisons between noun inflection (plural) and noun naming by ROI. The time axis lists both trial time since picture onset time (top row numbers), as well as time relative to condition icon onset (second row italic numbers). Following the same plotting schemes from figure 4, line traces indicate when Hotelling *T^2^*-test showed a significant difference between the two conditions. Positive values show greater noun inflection response (noun plural > noun naming) by ROI, while negative values show greater noun naming response by ROI. The color saturation of the line traces reflects the percentage of significant voxels. The anatomical plots (bottom row) show significant response differences between the two conditions across the whole brain. Each anatomical plot represents an 80 ms average bin centered at the time indicated by the gray arrow.

**Table 2.**
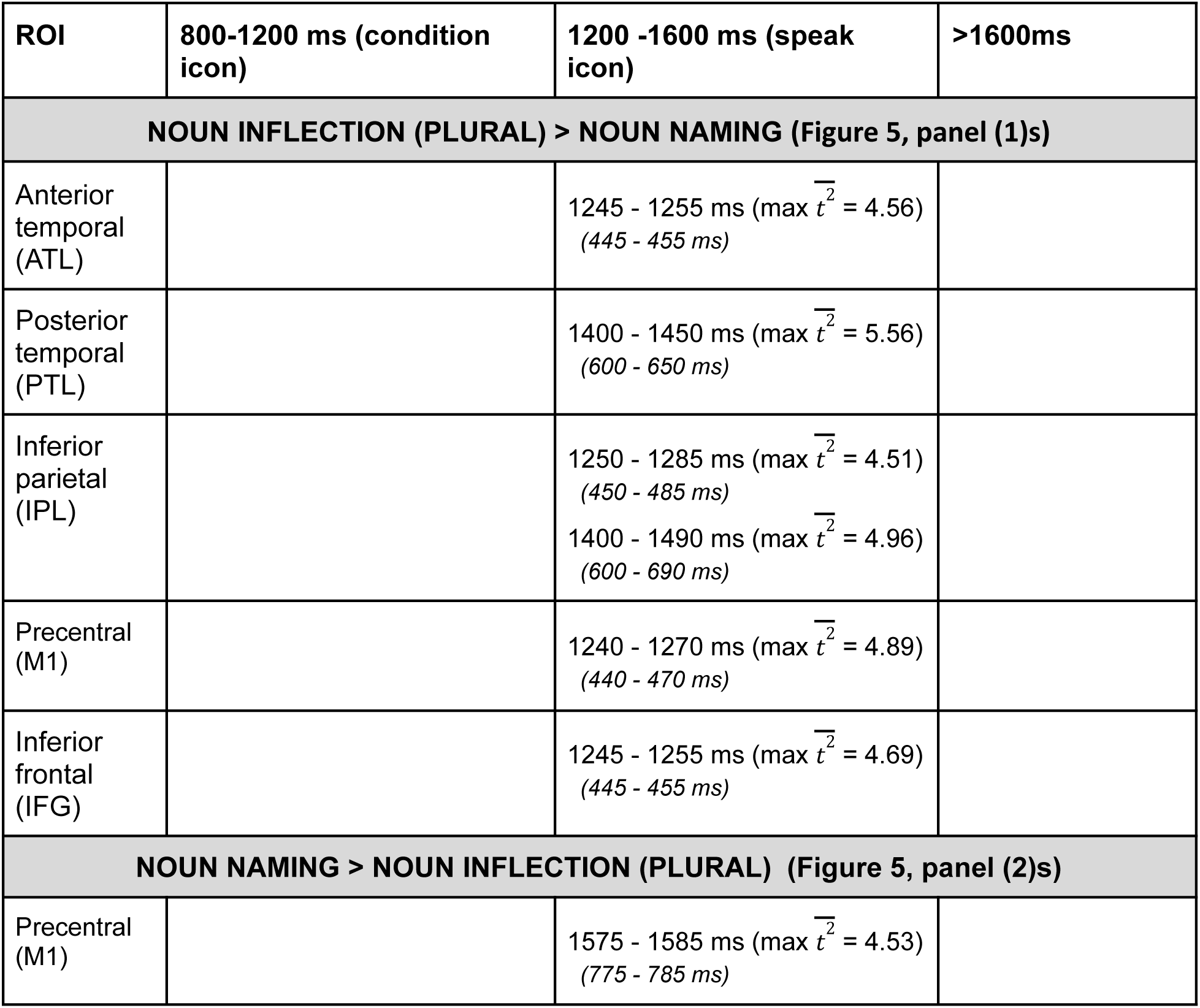
Significant differences between noun inflection (plural) and noun naming contrast in the five left hemisphere regions of interest (ROI) across the experimental trial.

| ROI | 800-1200 ms (condition icon) | 1200 -1600 ms (speak icon) | >1600ms |
| --- | --- | --- | --- |
| **NOUN INFLECTION (PLURAL) > NOUN NAMING (Figure 5, panel (1)s)** |  |  |  |
| Anterior temporal (ATL) | | 1245 - 1255 ms ($\max \bar{t}^2 = 4.56$)<br>(445 - 455 ms) | |
| Posterior temporal (PTL) | | 1400 - 1450 ms ($\max \bar{t}^2 = 5.56$)<br>(600 - 650 ms) | |
| Inferior parietal (IPL) | | 1250 - 1285 ms ($\max \bar{t}^2 = 4.51$)<br>(450 - 485 ms)<br>1400 - 1490 ms ($\max \bar{t}^2 = 4.96$)<br>(600 - 690 ms) | |
| Precentral (M1) | | 1240 - 1270 ms ($\max \bar{t}^2 = 4.89$)<br>(440 - 470 ms) | |
| Inferior frontal (IFG) | | 1245 - 1255 ms ($\max \bar{t}^2 = 4.69$)<br>(445 - 455 ms) | |
| **NOUN NAMING > NOUN INFLECTION (PLURAL) (Figure 5, panel (2)s)** |  |  |  |
| Precentral (M1) | | 1575 - 1585 ms ($\max \bar{t}^2 = 4.53$)<br>(775 - 785 ms) | |

Figure 6 and Table 3 show the noun constituent assembly vs. noun naming comparison results. Following the *condition* icon, noun constituent assembly (i.e., noun phrase > noun naming) showed an early activation of PTL at 255–275 ms, followed by IFG and ATL activation at 275-295 ms and the IPL shortly thereafter (Figure 6, panel (1)s). This is followed by biphasic activity of IFG, IPL, and PTL around 400-460 and 535-600 ms. M1 activity emerged later and showed three activation peaks between 485 and 585 ms following the *condition* icon. The comparison of noun naming > noun phrase showed an activation of PTL between 395-410 ms, followed by ATL between 435-455 ms, and finally M1 between 805-815 ms (Figure 6, panel (2)s).

**Figure 6.**
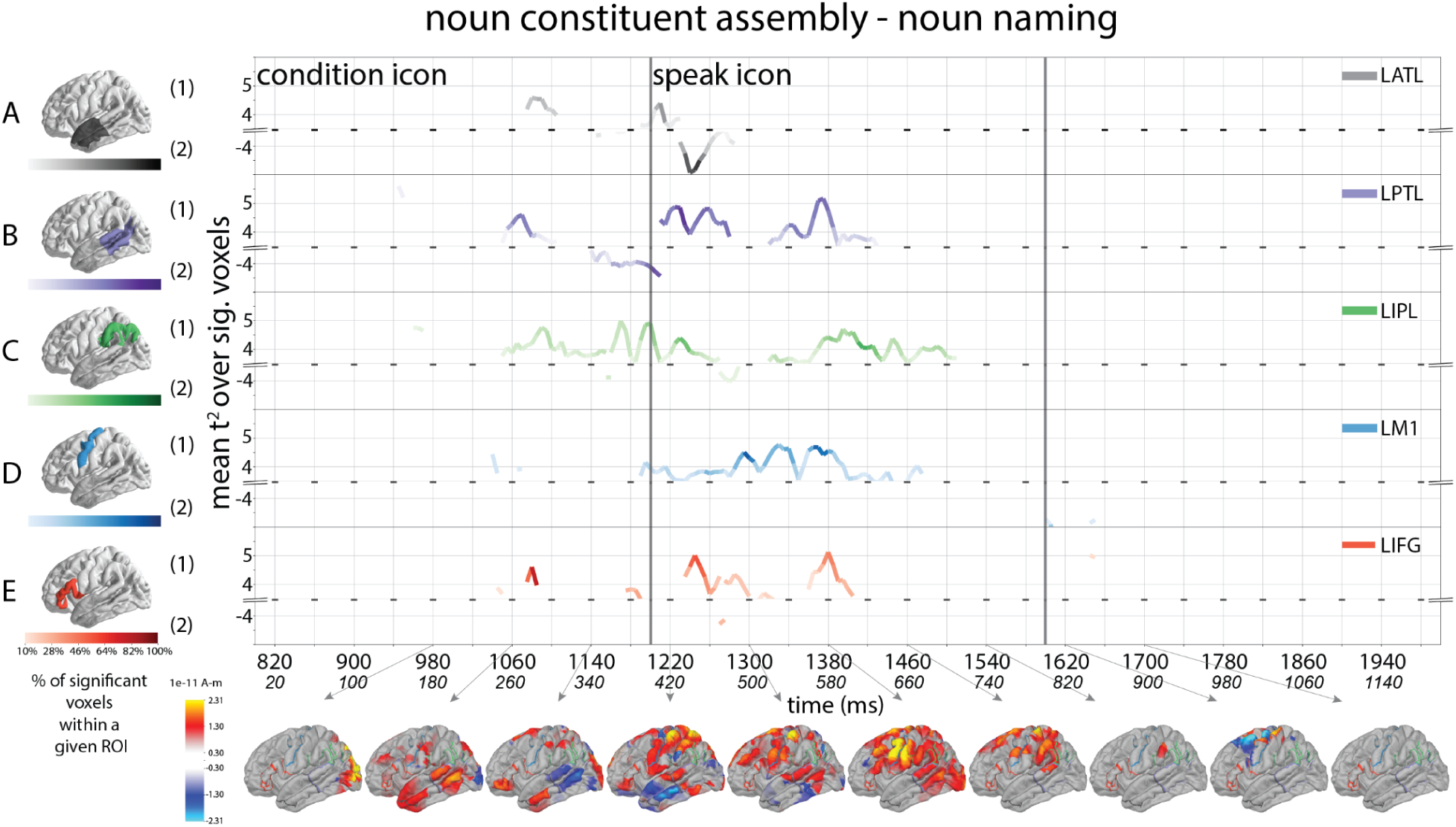
Results of direct comparisons between noun constituent assembly (phrase) and noun naming by ROI. Following the same plotting schemes from figure 4, line traces indicate when Hotelling *T^2^*-test showed a significant difference between the two conditions. Positive values show greater noun constituent assembly response (noun phrase > noun naming) by ROI, while negative values show greater noun naming response by ROI. The color saturation of the line traces reflects the percentage of significant voxels. The anatomical plots (bottom row) show significant response differences between the two conditions across the whole brain. Each anatomical plot represents an 80 ms average bin centered at the time indicated by the gray arrow.

**Table 3.**
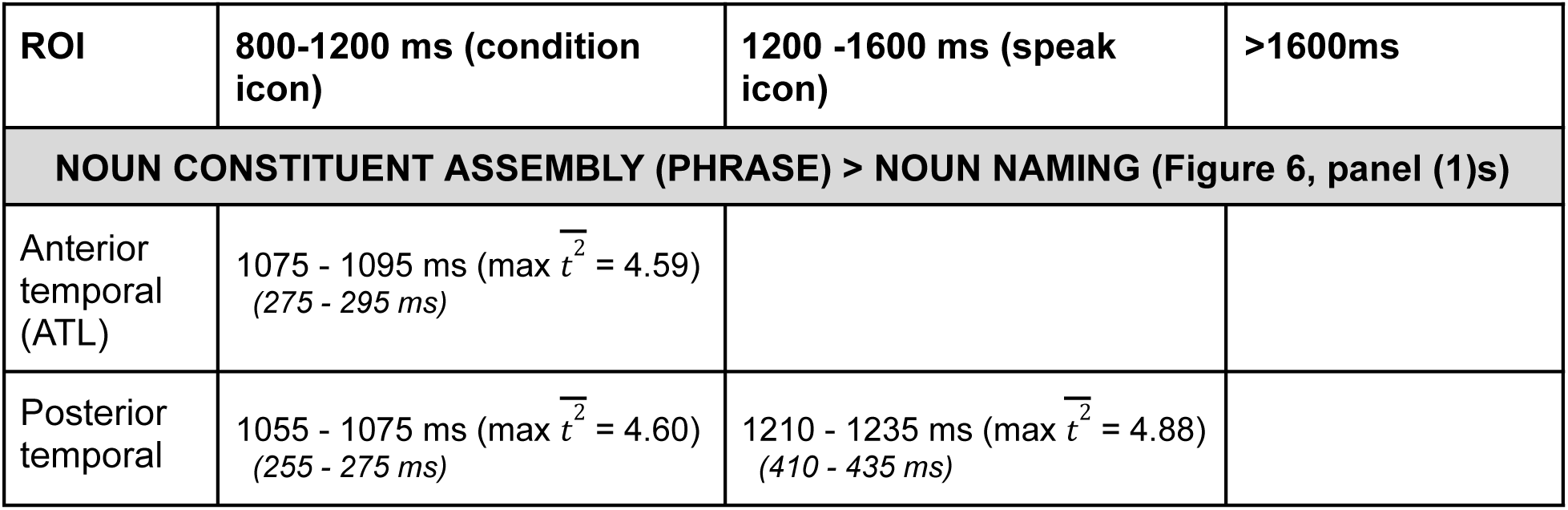
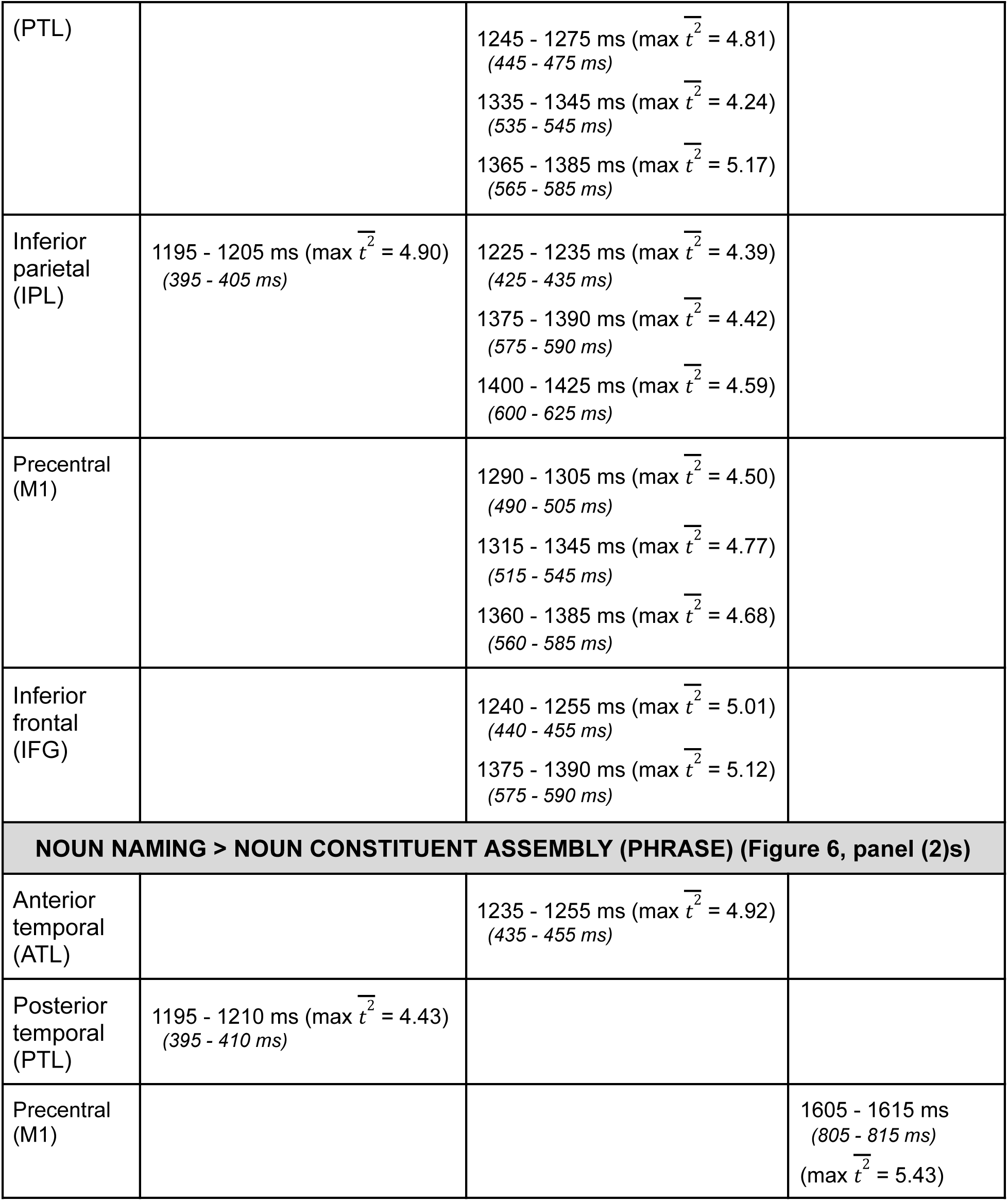
Significant differences between noun constituent assembly (phrase) and noun naming contrasts in the five left hemisphere ROIs across the experimental trial that met our reporting criteria (at least 30% ROI voxels for at least 10 ms and an averaged *T^2^* of at least 4). Times relative to the condition icon are italicized.

Figure 7 and Table 4 show the results between the two noun morphosyntax conditions (noun constituent assembly and noun inflection). Following the *condition* icon, noun constituent assembly (i.e., noun phrase > noun plural) showed an early activation of PTL and IPL between 140-180 ms, accompanied by M1 activation starting at 180 ms, and lasted until 620 ms. ATL was activated between 260-320 ms, and IFG showed intermittent activation between 200-700 ms (Figure 7, panel (1)s). The comparison of noun plural > noun phrase showed an early activation of IPL between 200-260 ms, followed by ATL and PTL between 340-440 ms, and a recurring activation of IPL between 460-500 ms, finally IFG, ATL, and PTL between 580-660 ms (Figure 7, panel (2)s).

**Figure 7.**
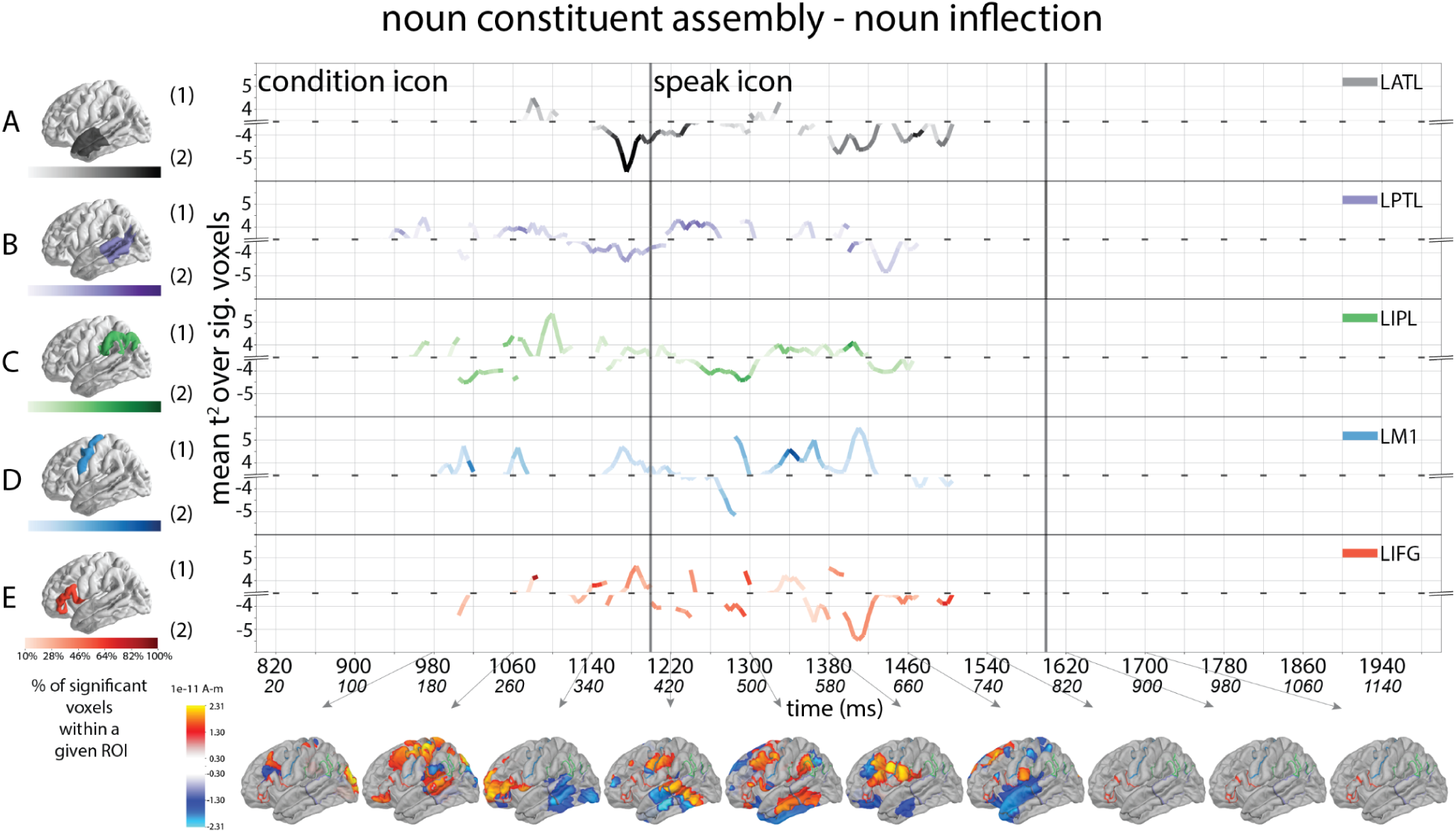
Results of direct comparisons between noun constituent assembly (noun phrase) and noun inflection (noun plural) by ROI. Following the same plotting schemes of Figure 4, line traces indicate when Hotelling *T^2^*-test showed a significant difference between the two conditions. Positive values show greater noun constituent assembly response (noun phrase > noun plural) by ROI, while negative values show greater noun inflection response by ROI. The color saturation of the line traces reflects the percentage of significant voxels. The anatomical plots (bottom row) show significant response differences between the two conditions across the whole brain. Each anatomical plot represents an 80 ms average bin centered at the time indicated by the gray arrow.

**Table 4.**
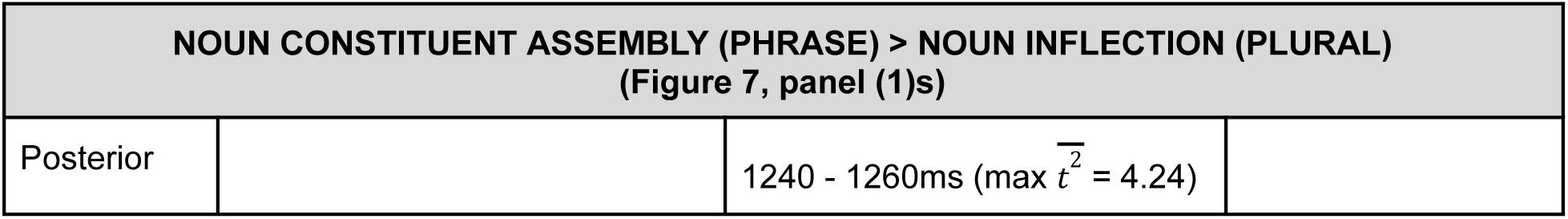
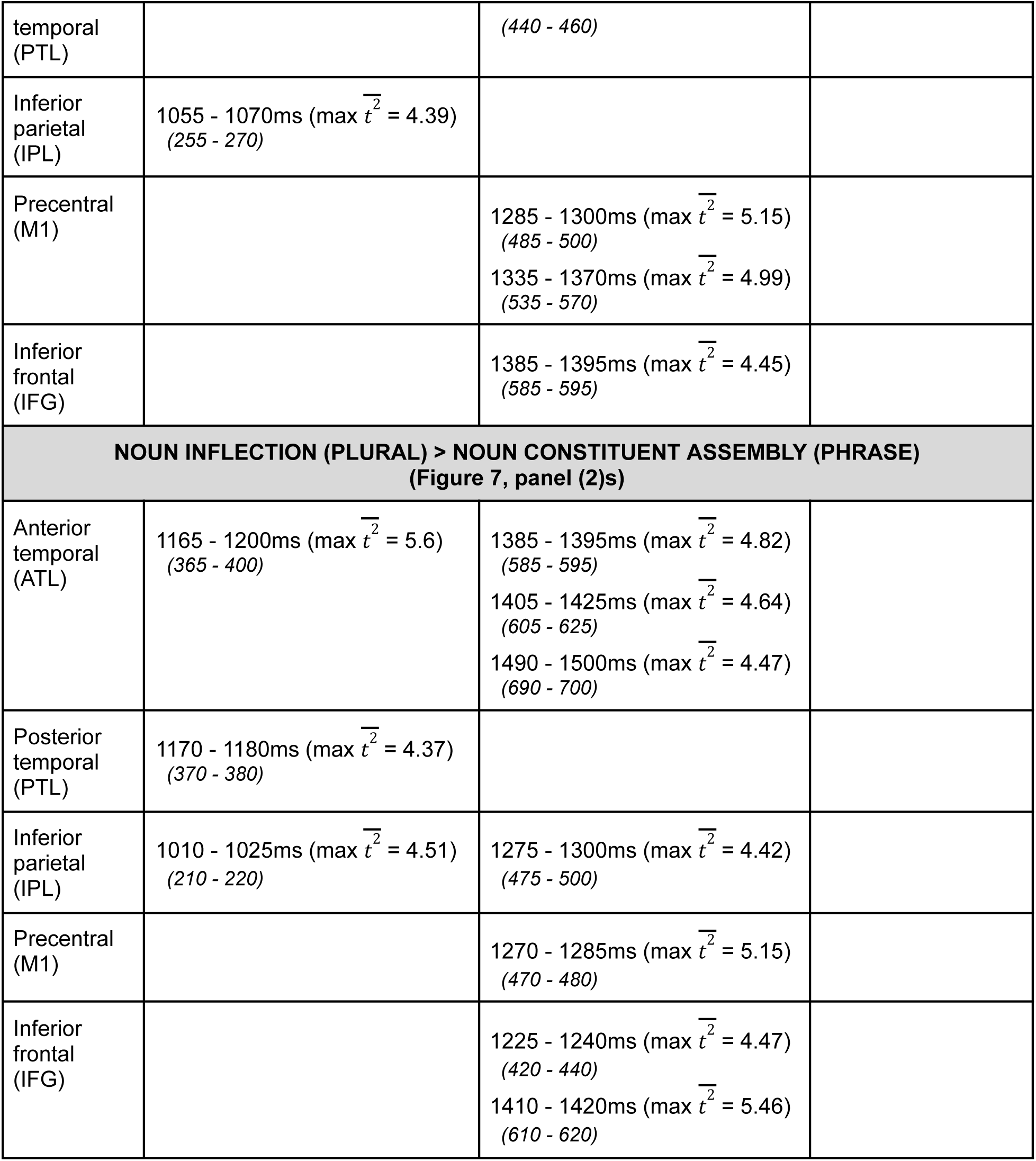
Significant differences between the two noun morphosyntax conditions in the five left hemisphere ROIs across the experimental trial that met our reporting criteria (at least 30% ROI voxels for at least 10 ms and an averaged *T^2^* of at least 4). Times relative to the condition icon are italicized.

##### 3.2.2.2. Verb morphosyntax

Figure 8 and Table 5 show the verb constituent assembly (future tense) vs. verb naming comparison results. Following the *condition* icon, verb constituent assembly (i.e., verb future > verb naming) was associated with a phase of activation of PTL, IPL, and M1 between 280 – 380 ms, followed by ATL and IFG activation between 535-565 ms, then again 700 - 740 ms. ATL and PTL re-engagement occurred between 975-1000 ms (Figure 8, panel (1)s). On the other hand, the comparison between verb naming > verb inflection future was associated with IFG activation between 325-335 ms, followed by PTL activation between 595-615 ms, M1 between 680-690 ms, and finally IPL between 700-910 ms (Figure 8, panel (2)s).

**Figure 8.**
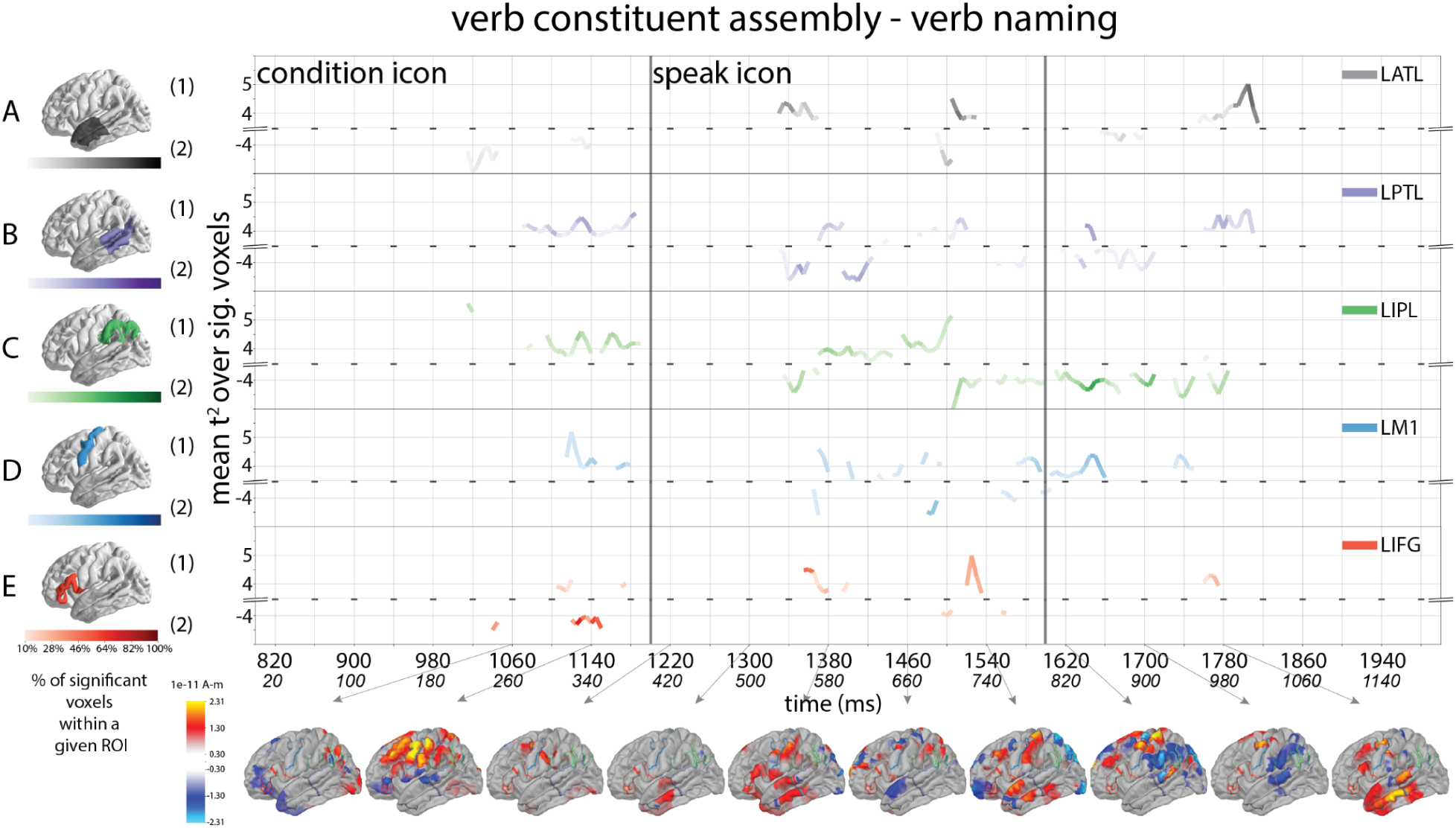
Results of direct comparisons between verb constituent assembly (future tense) and verb naming by ROI. Following the same plotting schemes of Figure 4, line traces indicate when Hotelling *T^2^*-test showed a significant difference between the two conditions. Positive values show greater verb constituent assembly response (verb future tense > verb naming) by ROI, while negative values show greater verb naming response by ROI. The color saturation of the line traces reflects the percentage of significant voxels. The anatomical plots (bottom row) show significant response differences between the two conditions across the whole brain. Each anatomical plot represents an 80 ms average bin centered at the time indicated by the gray arrow.

**Table 5.** Significant differences between verb morphosyntax and lexical access contrasts in the five left hemisphere ROIs across the experimental trial that met our reporting criteria (at least 30% ROI voxels for at least 10 ms and an averaged *T^2^*of at least 4). Times relative to the condition icon are italicized.

| ROI | 800-1200 ms (condition icon) | 1200 -1600 ms (speak icon) | >1600ms |
| --- | --- | --- | --- |
| **VERB CONSTITUENT ASSEMBLY (FUTURE) > VERB NAMING (Figure 8, panel (1)s)** |  |  |  |
| Anterior temporal (ATL) | | 1335 - 1345 ms ($\max \overline{t^2} = 4.36$)<br>(535 - 545 ms) | 1790 - 1810 ms<br>(990 - 1010 ms)<br>($\max \overline{t^2} = 5.01$) |
| Posterior temporal (PTL) | 1125 - 1135 ms ($\max \overline{t^2} = 4.46$)<br>(325 - 335 ms) | | 1775 - 1785 ms<br>(975 - 985 ms)<br>($\max \overline{t^2} = 4.56$) |
| Inferior frontal (IFG) | | 1355 - 1365 ms ($\max \overline{t^2} = 4.51$)<br>(555 - 565 ms) | |
| **VERB NAMING > VERB CONSTITUENT ASSEMBLY (FUTURE) (Figure 8, panel (2)s)** |  |  |  |
| Posterior temporal (PTL) | | 1395 - 1415 ms ($\max \overline{t^2} = 4.64$)<br>(595 - 615 ms) | |
| Inferior parietal (IPL) | | | 1635 - 1655 ms<br>(835 - 855 ms)<br>($\max \overline{t^2} = 4.32$) |
| Precentral (M1) | | 1480 - 1490 ms ($\max \overline{t^2} = 4.56$)<br>(680 - 690 ms) | |
| Inferior frontal (IFG) | 1125 - 1150 ms ($\max \overline{t^2} = 4.44$)<br>(325 - 350 ms) | | |

Figure 9 and Table 6 show the verb inflection and constituent assembly (past tense) vs. verb naming comparison results. Following the *condition* icon, verb inflection plus constituent assembly > verb naming (i.e., verb past > verb naming) was associated with activation of ATL and PTL regions between 160-200 ms, followed by activation in the IPL and PTL regions between 490-535 ms, then ATL between 655-745 ms. M1 showed two phases of activity, one between 180-400 ms, and another one between 580 - 900 ms (Figure 9, panel(1)s). The comparison of verb naming > verb inflection plus constituent assembly was associated with ATL, PTL, and IPL activation between 260-400 ms, followed by re-engagement of ATL and PTL between 620-660 ms (Figure 9, panel (2)s).

**Figure 9.**
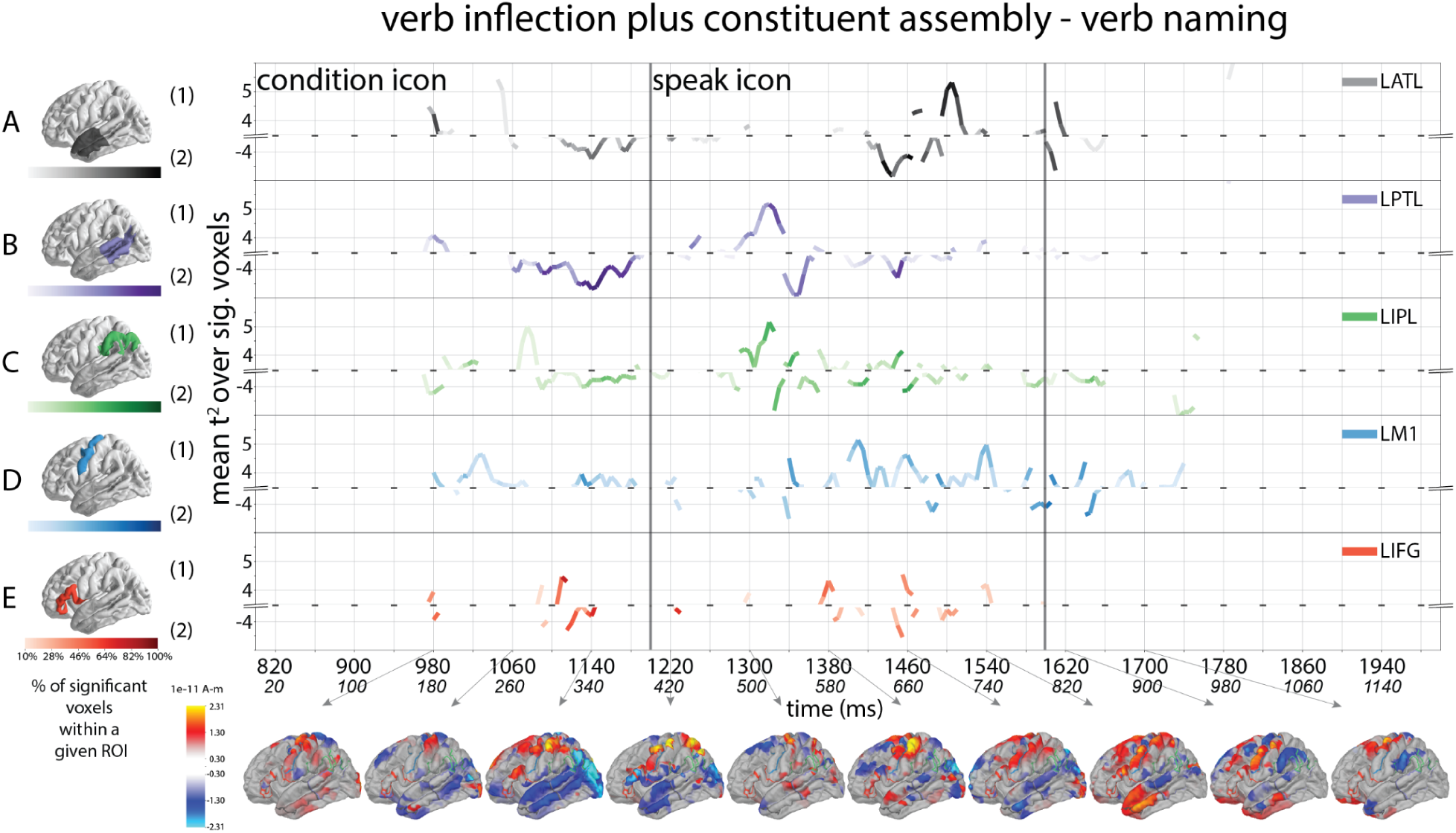
Results of direct comparisons between verb inflection plus constituent assembly (past tense) and verb naming by ROI. Following the same plotting schemes of figure 4, line traces indicate when Hotelling *T^2^*-test showed a significant difference between the two conditions. Positive values show greater verb inflection plus constituent assembly response (verb past tense > verb naming) by ROI, while negative values show greater verb naming response by ROI. The color saturation of the line traces reflects the percentage of significant voxels. The anatomical plots (bottom row) show significant response differences between the two conditions across the whole brain. Each anatomical plot represents an 80 ms average bin centered at the time indicated by the gray arrow.

**Table 6.**
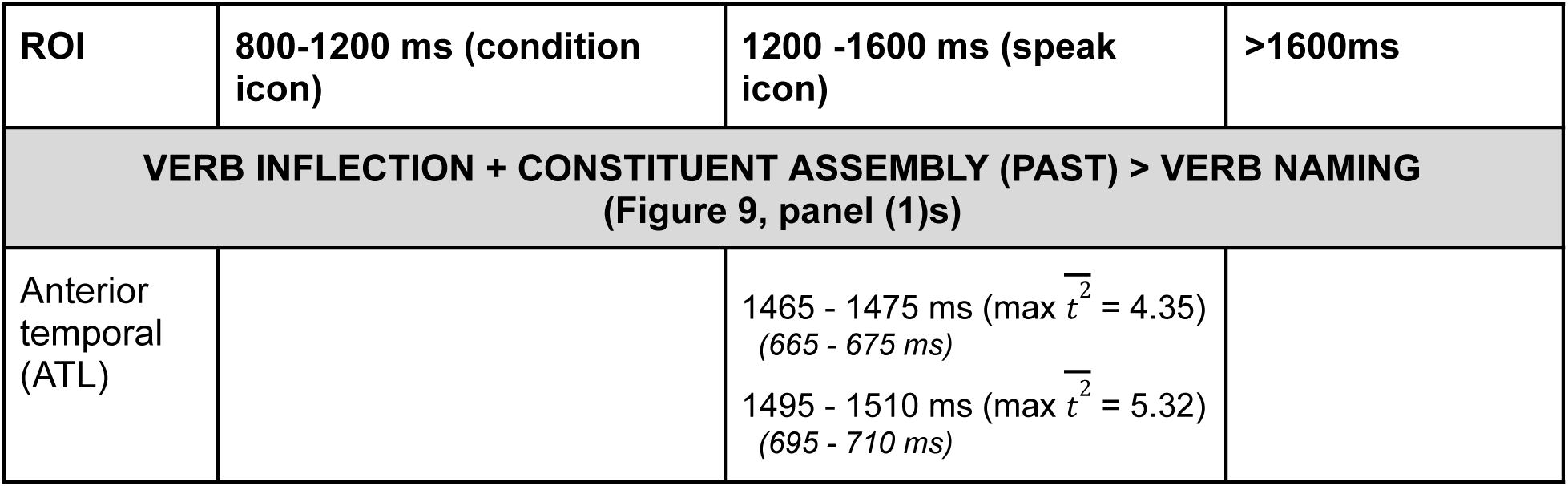
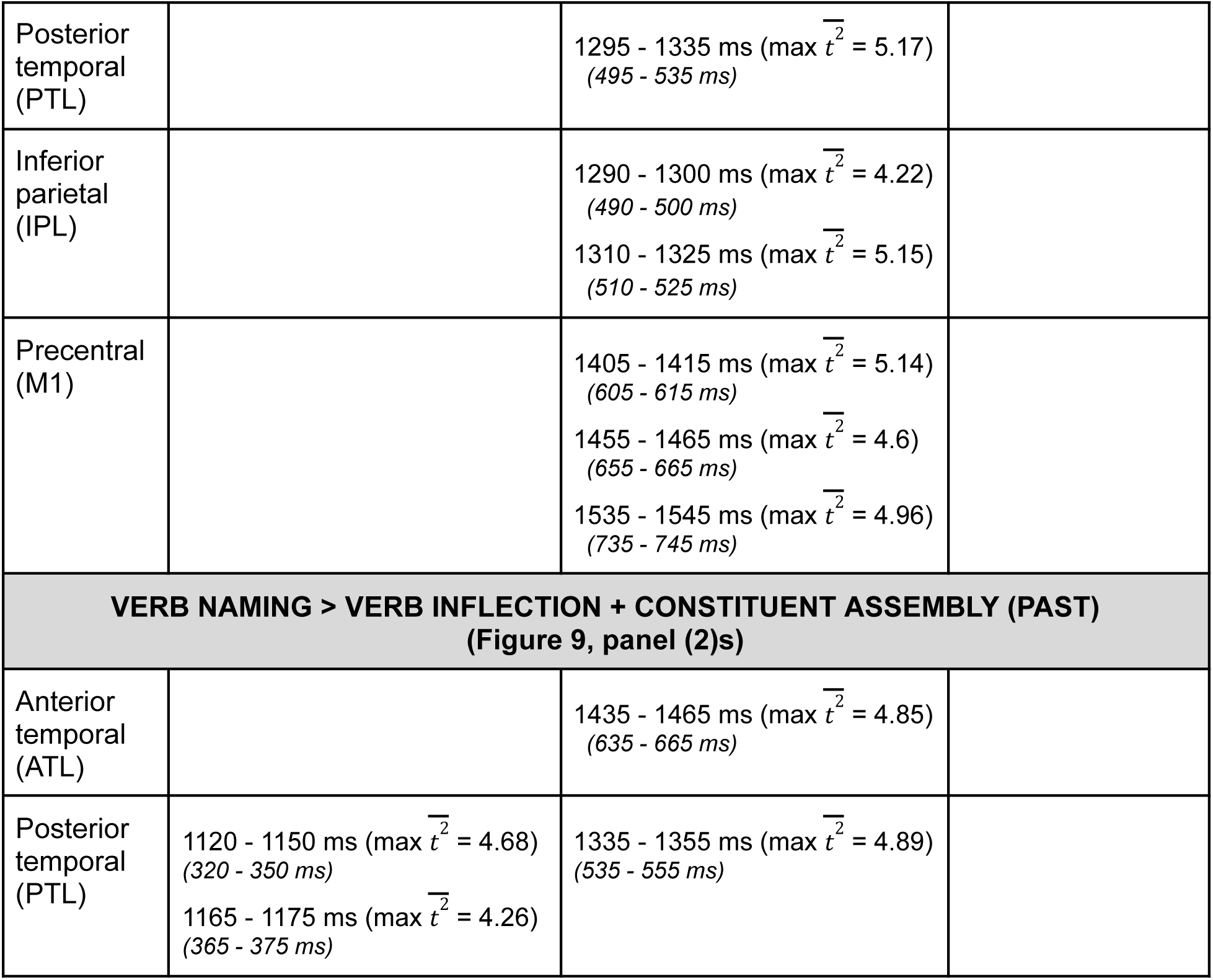
Significant differences between verb morphosyntax and lexical access contrasts in the five left hemisphere ROIs across the experimental trial that met our reporting criteria (at least 30% ROI voxels for at least 10 ms and an averaged *T^2^*of at least 4). Times relative to the condition icon are italicized.

| ROI | 800-1200 ms (condition icon) | 1200 -1600 ms (speak icon) | >1600ms |
| --- | --- | --- | --- |
| **VERB INFLECTION + CONSTITUENT ASSEMBLY (PAST) > VERB NAMING**<br>(Figure 9, panel (1)s) |  |  |  |
| Anterior temporal (ATL) | | 1465 - 1475 ms ($\max \bar{t^2} = 4.35$)<br>(665 - 675 ms)<br>1495 - 1510 ms ($\max \bar{t^2} = 5.32$)<br>(695 - 710 ms) | |
| Posterior temporal (PTL) | | 1295 - 1335 ms ($\max \overline{t^2} = 5.17$)<br>(495 - 535 ms) | |
| Inferior parietal (IPL) | | 1290 - 1300 ms ($\max \overline{t^2} = 4.22$)<br>(490 - 500 ms)<br>1310 - 1325 ms ($\max \overline{t^2} = 5.15$)<br>(510 - 525 ms) | |
| Precentral (M1) | | 1405 - 1415 ms ($\max \overline{t^2} = 5.14$)<br>(605 - 615 ms)<br>1455 - 1465 ms ($\max \overline{t^2} = 4.6$)<br>(655 - 665 ms)<br>1535 - 1545 ms ($\max \overline{t^2} = 4.96$)<br>(735 - 745 ms) | |
| **VERB NAMING > VERB INFLECTION + CONSTITUENT ASSEMBLY (PAST)**<br>**(Figure 9, panel (2)s)** |  |  |  |
| Anterior temporal (ATL) | | 1435 - 1465 ms ($\max \overline{t^2} = 4.85$)<br>(635 - 665 ms) | |
| Posterior temporal (PTL) | 1120 - 1150 ms ($\max \overline{t^2} = 4.68$)<br>(320 - 350 ms)<br>1165 - 1175 ms ($\max \overline{t^2} = 4.26$)<br>(365 - 375 ms) | 1335 - 1355 ms ($\max \overline{t^2} = 4.89$)<br>(535 - 555 ms) | |

Figure 10 and Table 7 show the results between the two verb morphosyntax conditions (verb inflection plus constituent assembly and verb constituent assembly). Following the *condition* icon, verb inflection plus constituent assembly (i.e., verb past tense > verb future tense) showed an early activation of PTL, IPL, and M1 between 160-200 ms (Figure 10, panel (1)s). The comparison of verb future tense > verb past tense showed sustained activation of ATL, PTL, and IPL between 260-460 ms, one peak of activation of M1 between 320-340 ms, then again another episode of activation between 620-720 ms for PTL and IPL (Figure 10, panel (2)s).

**Figure 10.**
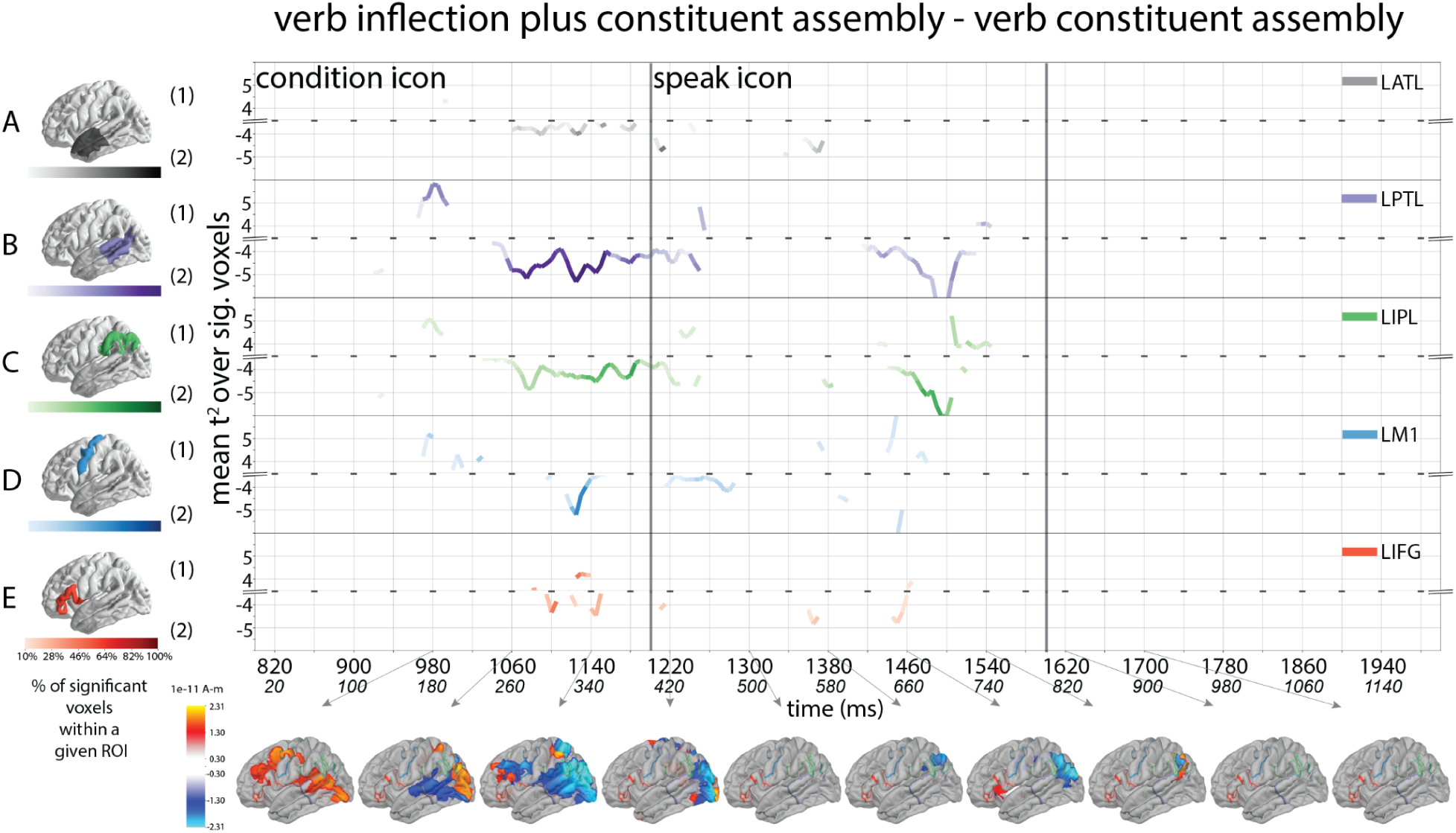
Results of direct comparisons between verb inflection plus constituent assembly (past tense) and verb constituent assembly (future tense) by ROI. Following the same plotting schemes of figure 4, line traces indicate when Hotelling *T^2^*-test showed a significant difference between the two conditions. Positive values show greater verb inflection plus constituent assembly response (verb past tense > verb future tense) by ROI, while negative values show greater verb constituent assembly response by ROI. The color saturation of the line traces reflects the percentage of significant voxels. The anatomical plots (bottom row) show significant response differences between the two conditions across the whole brain. Each anatomical plot represents an 80 ms average bin centered at the time indicated by the gray arrow.

**Table 7.**
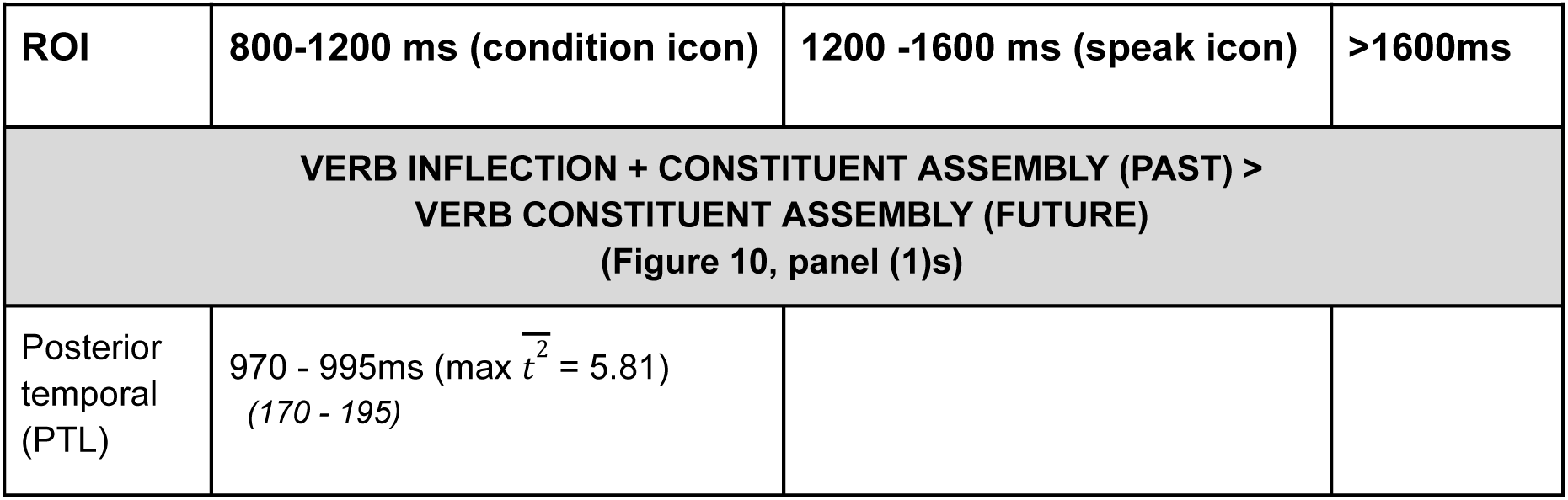
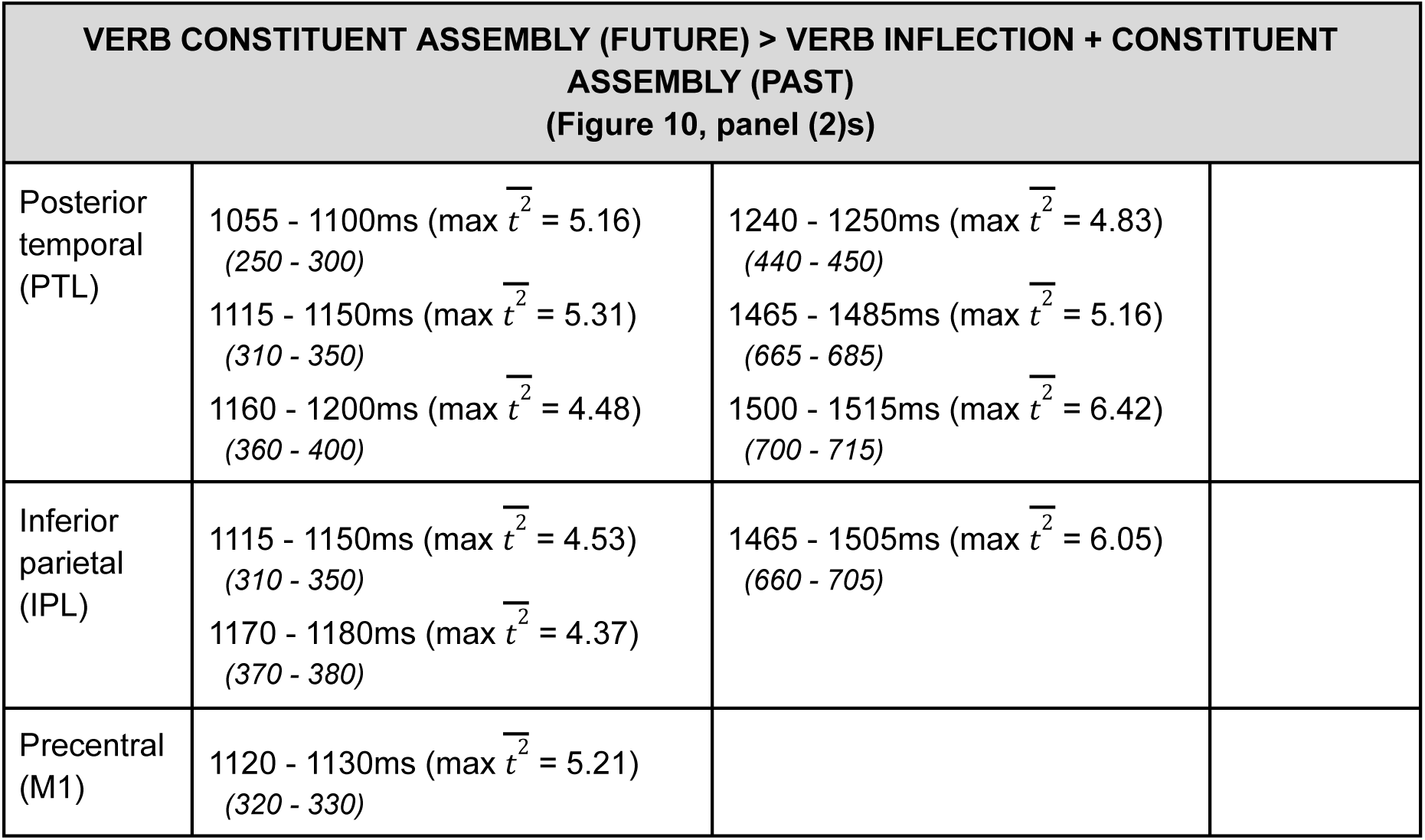
Significant differences between two verb morphosyntax conditions in the five left hemisphere ROIs across the experimental trial that met our reporting criteria (at least 30% ROI voxels for at least 10 ms and an averaged *T^2^* of at least 4). Times relative to the condition icon are italicized.

### 3.3. Speech onset locked analyses

The left hemisphere responses that met our reporting criteria (at least 30% ROI voxels active for at least 10 ms and an average *T^2^* of at least 4) are listed in Tables 8-9. All significant voxels are provided in Supplementary Table S4. Figures 11-12 illustrate the time course and associated spatial distribution of each contrast.

**Figure 11.**
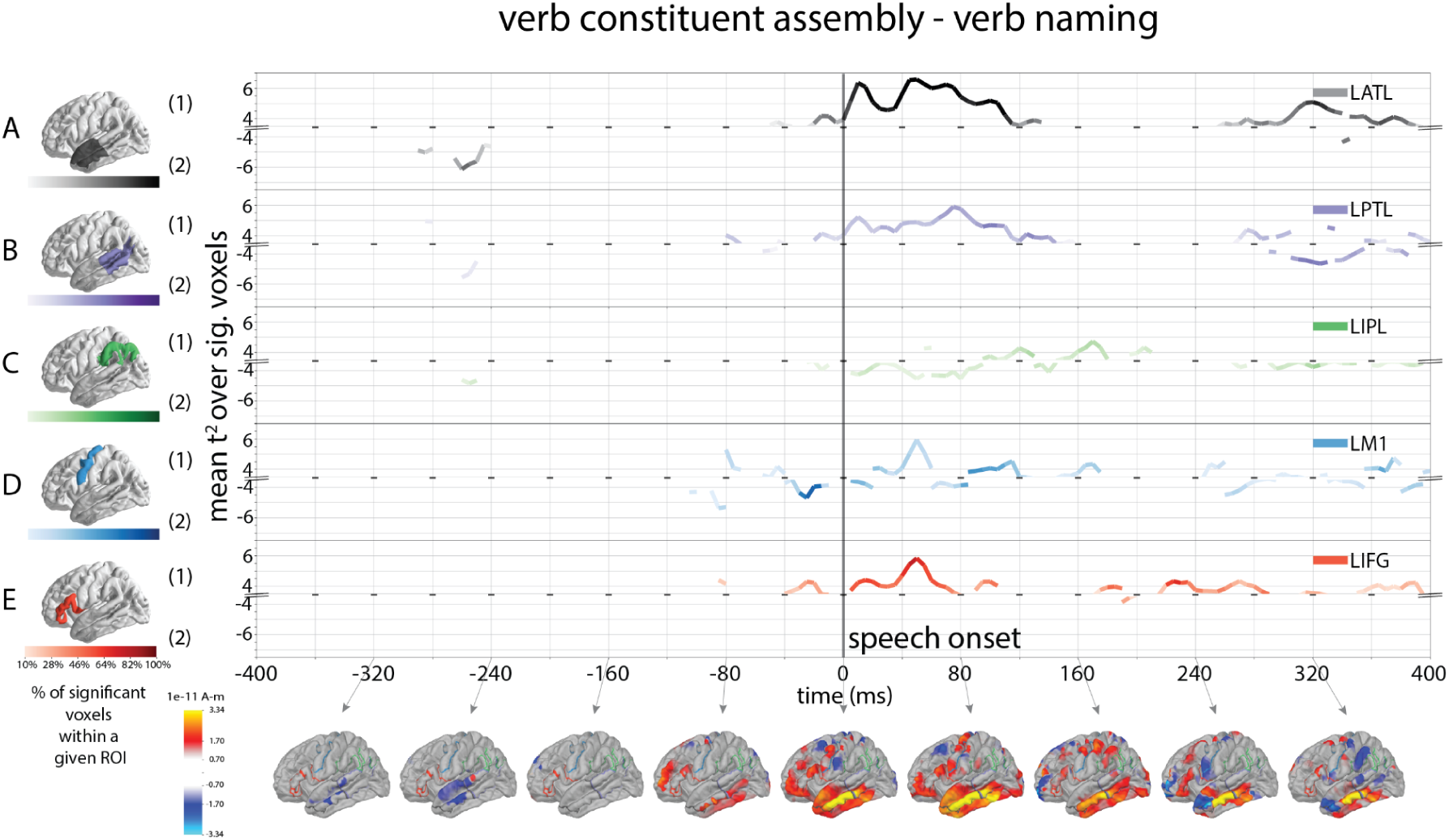
Results of direct comparisons between verb constituent assembly (future tense) and verb naming by ROI, time locked to speech onset. Following the same plotting schemes of Figure 4, line traces indicate when Hotelling *T^2^*-test showed a significant difference between the two conditions. Panel (1)s show greater verb constituent assembly response (verb constituent assembly (future tense) > verb naming) by ROI, while panel (2)s show greater verb naming response by ROI. The color saturation of the line traces reflects the percentage of significant voxels. The anatomical plots (bottom row) show significant response differences between the two conditions across the whole brain, averaged over an 80 ms bin centered by the timepoints with an arrow pointing to the brain.

**Figure 12.**
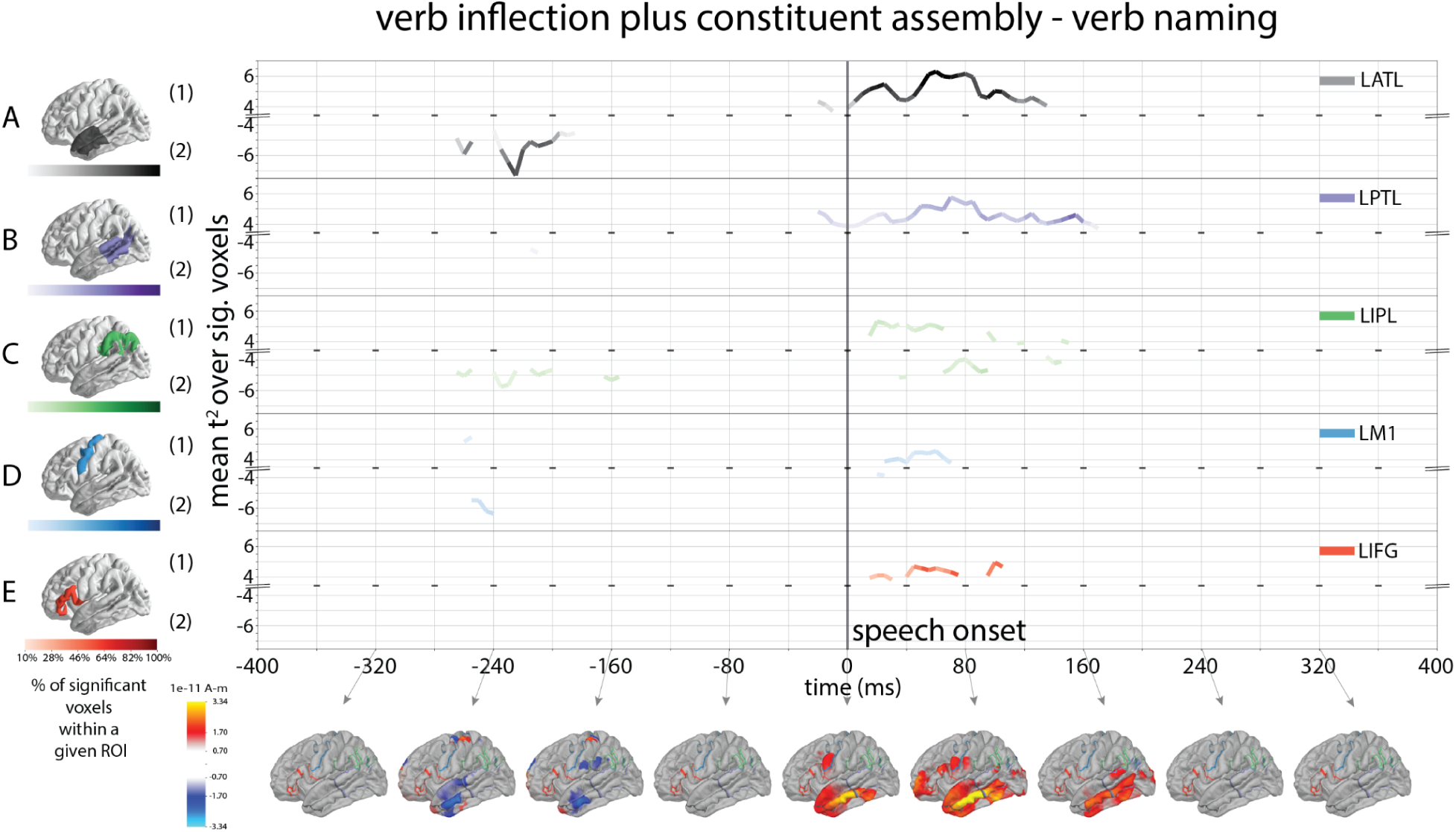
Results of direct comparisons between verb inflection plus constituent assembly (past tense) and verb naming by ROI, time locked to speech onset. Following the same plotting schemes of Figure 4, line traces indicate when Hotelling *T^2^*-test showed a significant difference between the two conditions. Panel (1)s show greater verb inflection plus constituent assembly response (verb past tense > verb naming) by ROI, while panel (2)s show greater verb naming response by ROI. The color saturation of the line traces reflects the percentage of significant voxels. The anatomical plots (bottom row) show significant response differences between the two conditions across the whole brain, averaged over an 80 ms bin centered by the timepoints with an arrow pointing to the brain.

**Table 8.**
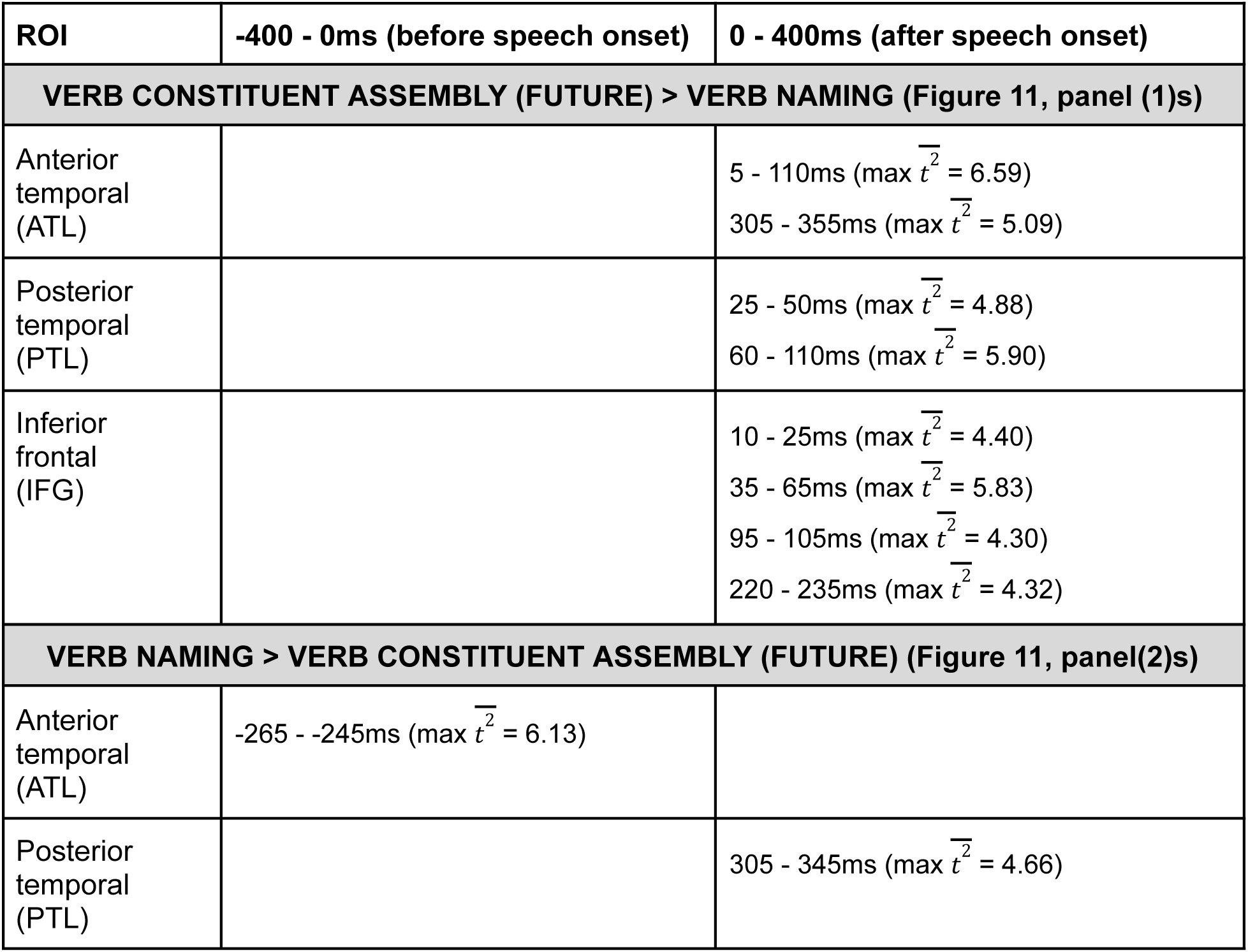
Significant differences between verb morphosyntax and verb naming contrasts in the hemisphere regions of interest (ROI) for the speech onset-locked analysis that met our reporting criteria (at least 30% ROI voxels for at least 10 ms and t^2^max of at least 4).

**Table 9.**
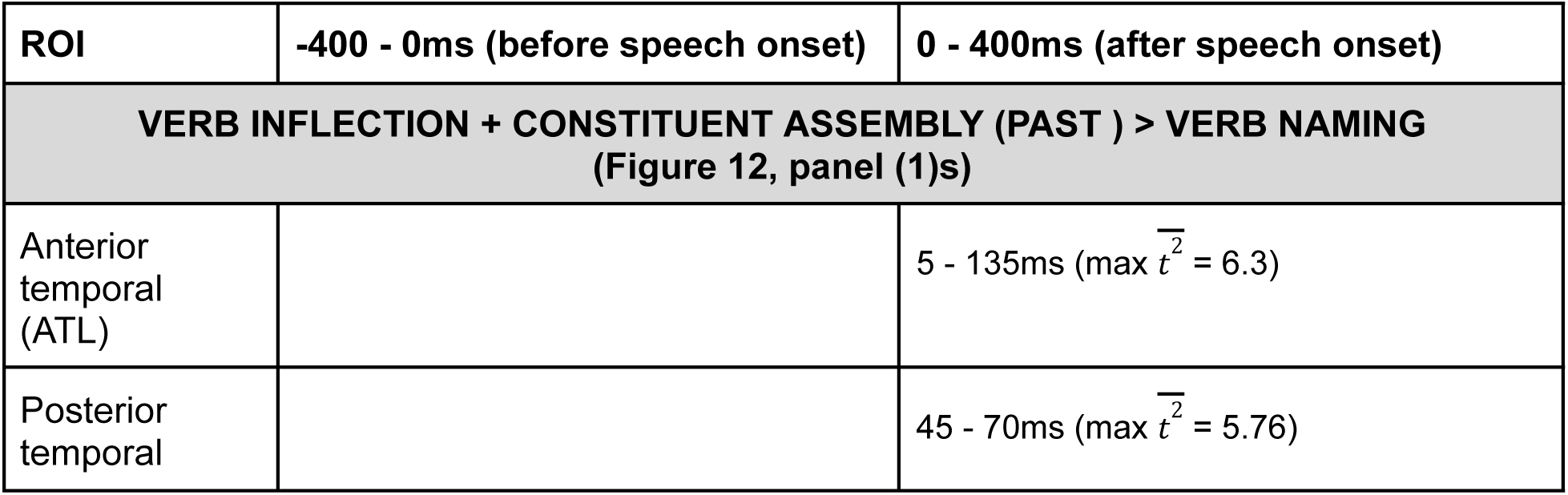
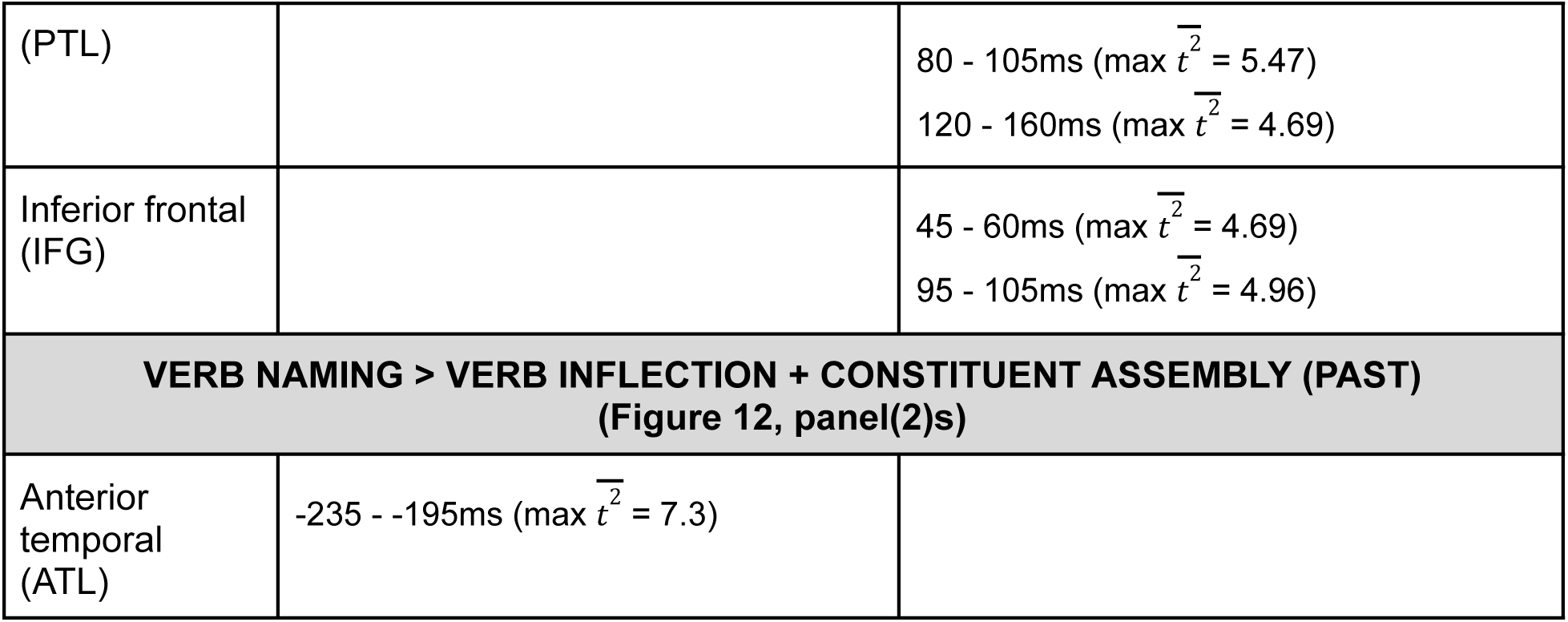
Significant differences between verb morphosyntax and verb naming contrasts in the hemisphere regions of interest (ROI) for the speech onset-locked analysis that met our reporting criteria (at least 30% ROI voxels for at least 10 ms and t^2^max of at least 4).

#### 3.3.1. Lexical retrieval

There were no activation differences in any of the language ROIs between noun naming and verb naming conditions within the 800 ms window centered at speech onset.

#### 3.3.2. Morphosyntactic planning

For both the comparisons between noun naming and each of the two noun morphosyntax conditions, no activation differences in any of the language ROIs were observed within the 800 ms window centered at the speech onset time.

For the verb morphosyntax contrasts, verb constituent assembly (future) > verb naming was associated with a sustained strong activation of ATL (5-110 ms after the response onset time), accompanied by IFG activation between 10-105 ms, and PTL between 25-110 ms (Figure 11, panel (1)s). Verb naming > verb constituent assembly was associated with a short activation of ATL 265 - 245 ms before speech onset, and again 305-345 ms after speech onset (Figure 11, panel (2)s).

Similar to the above pattern, verb inflection plus constituent assembly (past) > verb naming was associated with a sustained strong activation of ATL (5-135 ms after speech onset), accompanied by IFG activation 45-105 ms and PTL activation 45-160 ms after speech onset. Meanwhile, verb naming > verb inflection plus constituent assembly was associated with 235 - 195 ms of ATL activation before speech onset (Figure 12, panel (1)s). This latter pattern is similar to the verb naming > verb constituent assembly contrast (Figure 12, panel (2)s).

## 4. Discussion

This study sought to delineate the spatiotemporal response to subcomponents of overt sentence production, particularly lexical and morphosyntactic processes. The use of MEG allowed us to track the time-locked flow of neural activity from the beginning of stimulus presentation until the completion of speech production over a trial duration that exceeded two seconds. The MEG results showed unique spatiotemporal patterns depending on the lexical category (noun versus verb) and the specific morphosyntactic processes.

Before discussing the implications of the findings within the context of prior literature, it is important to underscore that contrasting results across studies could arise from the use of different brain imaging methods, analyses procedures, and experimental paradigms. MEG and functional magnetic resonance imaging (fMRI) have been shown to yield different results for language (and motor) tasks, with MEG showing greater sensitivity to temporal and occipital activity and fMRI being more sensitive to inferior frontal and subcortical activations (Ellis et al., 2020; Liljeström et al., 2009; McDonald et al., 2010; Vartiainen et al., 2011). The ROI sizes used in the present study (43 - 126 voxels) are compatible with the spatial resolution of MEG, but larger than those reported in many fMRI studies. Further, some of these ROIs have functionally specialized subregions for language which warrants caution in attributing specific linguistic functions to these larger ROIs (e.g., Amunts et al., 2010; Flinker et al., 2010; Ishkhanyan et al., 2020; Vasileiadi et al., 2023; Zaccarella & Friederici, 2020).

Different MEG source localization approaches, such as using individual structural MRI versus a template brain, or applying free-vector versus fixed-orientation constraints in the inverse solution can yield different results. A free-vector orientation model was used in our current study to accommodate inter-individual neuroanatomical variability. While cortical macro-currents are typically assumed to be oriented approximately normal to the cortical surface, cortical folding varies between individuals, and is imperfectly modeled by a template brain. A free-vector orientation constraint can account for this by accommodating more individual variability in current orientations across larger ROIs, and has been suggested to be beneficial for minimum norm estimate models under certain conditions (e.g., Henson et al., 2009).

Noteworthy aspects of the present study’s experimental paradigm include overt production, within-trial separation of lexical access (0-800 ms) and linguistic structure (800+ ms), randomized sequence of experimental conditions, and use of the same picture stimuli for lexical and morphosyntactic conditions within a lexical category.

### 4.1 Lexical retrieval

This study not only revealed the general pattern of neural activity associated with lexical retrieval (Figure 3), but also lexical category differences between object and action picture naming (Figure 4). The first notable finding was the relatively early engagement of all five ROIs during lexical retrieval. IPL and PTL were the earliest to respond at 120 ms post-picture, and followed by ATL, IFG, and M1 at 200-400 ms post-picture. All ROIs sustained this activity until the *condition* icon was presented. This pattern of early and nearly simultaneous activation of ROIs is consistent with findings of early neural activation of temporal and frontal regions between 130 and 240 ms (Miozzo et al., 2014; Mundig et al., 2016; Strijkers et al., 2017) and supports the simultaneous ignition hypothesis (Miozzo et al., 2015). The sustained activity of all ROIs until the presentation of the *condition* icon is suggestive of neural reverberation proposed by Miozzo et al. (2014). A prior MEG study with the most similar experimental paradigm to the current study (randomized sequence, overt naming of verbs and nouns) used different data analysis procedures (Sörös et al., 2003) and found a temporo-parietal response that peaked at 200-400 ms and a fronto-parietal response that peaked at 400-600 ms. The dipoles identified by Sörös et al. (see Figure 5 of their paper) have considerable overlap with the ROIs in the present study, even including precentral regions and multiple parietal dipoles. However, only visual sources were activated within the first 200 ms after picture onset time, and no significant latency differences were found between action and object naming for their neurotypical population group within their ROIs.

The second notable finding of the present study is that the IPL consistently showed a strong and sustained response (reverberation). This is a noteworthy difference from Indefrey’s (2011) spatio-temporal model of word production which did not identify the IPL. The IPL (which includes the angular gyrus and supramarginal gyrus) has been associated with a variety of linguistic functions including semantics, phonological processing, stored motor programs, and articulatory sequencing (Brownsett & Wise, 2009; Oberhuber et al., 2016). Given its direct structural connections with sensory cortices, the IPL has also been identified as a multimodal convergence zone, especially for word retrieval (Binder & Desai, 2011; Kuhnke et al., 2023; Seghier, 2013). In addition to its lexical role, the sustained IPL activity observed in the present study is consistent with its role in the intention to speak and inner speech (Carota et al., 2010; Kuhnke et al., 2023; Wandelt et al., 2024). In an MEG study, Carota et al. (2010) found increased IPL activity whenever participants intended to speak, and this increased IPL activity was associated with concomitant activity in the IFG. This coactivation of IPL and IFG was assumed to be a cortical circuit for speech intention and preparation (Carota et al., 2010).

The third notable finding concerns the PTL. While the PTL was consistently activated by morphosyntactic planning, it also showed periods of stronger activity for picture naming, when compared with the four morphosyntax conditions (except noun inflection) (Tables 3-5). This suggests a unique role of the PTL in lexical retrieval in addition to morphosyntactic planning, and is consistent with the large body of literature identifying this region for both semantic and phonological aspects of words (Indefrey, 2011, Maess et al., 2002; Miozzo et al., 2014).

Fourth, we found an early and sustained M1 response, which started around 120 ms after picture presentation, peaked at 200-400 ms, and again at 600 ms. Given that these early M1 responses occurred much before the *condition* and *speak* icons were presented, the most likely explanation is feedforward activation of motor programs (Hickok, 2012; Miller & Guenther, 2021). The simultaneous ignition hypothesis of Miozzo et al. (2014) could be expanded to include the motor cortex (Mundig et al., 2015). M1 activity also increased 150-400 ms after the *condition* icon, and stayed elevated after the *speak* icon. These timelines are consistent with actual motor planning and articulation because the speaker has knowledge of what to say at this time. The motor planning timeline is consistent with prior research (Carota et al., 2023; Munding et al., 2016), but is considerably faster than the timeline described by Indefrey (2011).

Finally, there were several differences between noun and verb picture naming (Figure 4). Consistent with prior research, verb naming elicited a stronger and more widespread fronto-temporal response that sequentially included the PTL (130 - 140 ms), IFG (215 - 255 ms), and ATL (325 - 405 ms) (Bedny et al., 2012; Berlingeri et al., 2008; Faroqi-Shah et al., 2018 contra Liljeström et al., 2009). Stronger responses to verbs have been attributed to both more complex event knowledge (semantics) associated with verbs as well as the often more complex morphosyntactic representations of verbs, which include argument structure information and a larger variety of inflectional configurations (Berlingeri et al., 2008; Hauptman et al., 2022; Vigliocco et al., 2011). The earliest PTL response is consistent with the identified role of this area for lexical access and action concepts (Bedny et al., 2012). The subsequent IFG response can be associated with morphological differences (Sahin et al., 2009; Shapiro et al., 2006).

Interestingly, these same three ROIs showed a stronger response to noun naming compared to verb naming at a later timeline (Table 2; PTL: 365-400 ms, IFG: 410-425 ms, ATL: 405-420 ms). The larger IPL response for verbs occurred at a much later time point (> 715 ms), and could be attributed to verb-argument structure (Thompson et al., 2007). There was no increased engagement of the motor cortex for verbs (until 1400 ms). This finding contrasts with theories of embodied cognition, which suggest that verb concepts include cell assemblies in the motor cortex (Pulvermuller, 1999). Finally, speech onset locked analysis showed a stronger ATL response for verb naming relative to verb morphosyntax in the 200 ms immediately preceding articulation (Table 8 & 9). This is an intriguing finding considering the ATL’s role in conceptual combination (Pylkkanen, 2019). Perhaps this pre-articulatory ATL response is associated with re-activation of the verb concept. It is also possible that ATL’s involvement can happen at a later time when additional morphosyntactic planning is required. Future research can test these explanations of ATL activation.

The present study adds to prior understanding of word production in several ways. It not only demonstrates the overall simultaneous nature of the cortical response to word production, but also identifies the time course of ebb and flow of individual ROIs, including a prominent role of IPL. Next, this study delineates spatial and temporal differences in cortical responses across lexical categories - it shows that verb naming is associated with both a stronger and an earlier cortical response than noun naming in language ROIs. Finally, this study integrates word production and motor planning models by showing that M1 responds early and concurrently with language planning.

### 4.2 Morphosyntactic planning

The present study’s trial structure separated the brain response to lexical access (the first 800 ms after picture presentation) from utterance planning, which started after the *condition* icon was presented. Consistent with this, there were no activation differences between any of the morphosyntax conditions within a lexical category for the first 800 ms after picture presentation.

One potential confound across the lexical and morphosyntactic conditions is that the length of the verbal response (number of syllables) was different across experimental conditions. In particular, the conditions with constituent assembly resulted in multiword responses compared to picture naming and noun inflection. To test if syllable length differences across conditions could have contributed to the observed effect, a two-stage regression analysis of MEG source-level responses was performed using the number of syllables as a continuous regressor. The results are given in Supplementary Table S5 and Figure S4. Although the number of syllables significantly contributes to response in M1 and IFG, the temporal profile of this syllable length effect was distinct from the morphosyntactic responses of IFG and M1 reported in Tables 3-5. Thus it is unlikely that the morphosyntactic effects from IFG and M1 were driven primarily by the verbal response length. The distinct temporal profile further suggests that syllable length and morphosyntactic planning may contribute to neural responses at different stages of sentence production.

#### 4.2.1. Nouns

The response for noun inflection (“trees” > “tree”) emerged 440 ms after the *condition* icon and simultaneously engaged the IFG, IPL, ATL, and M1. The PTL was engaged at about 600 ms, and the IFG, IPL and M1 were re-engaged at this same time. The cortical regions activated for noun plurals are consistent with overt and covert noun inflection generation in prior studies (e.g., Beretta et al., 2003; Marangolo et al., 2006). The typical functions associated with these regions include, concepts and semantics (ATL, IPL), lexical syntax (PTL), inflection (IFG), and motor planning (M1). This response was more widespread and more prolonged than the ATL-ventral prefrontal activity reported in a prior MEG study of noun plurals (Hauptman et al., 2022). The present study’s response also started about 100 ms later than the response reported in Hauptman et al. A possible explanation is the additional time required for processing the *condition* icons in the present study.

In contrast to noun inflections, the neural response for constituent assembly (“a blue tree” > “tree”; “a blue tree” > “trees”) started earlier and engaged the PTL, IPL and ATL in quick succession (Tables 3, 4, Supplementary S3, Figures 5 - 7). Given that the constituent assembly trials required participants to produce a determiner + color + noun sequence, the initial PTL, ATL and IPL response can be interpreted as involving conceptual combination (color + noun) and planning of the noun phrase structure. This is consistent with the roles of these regions in conceptual combination, integrating syntactic and semantic information, and generating phrase structure during language production (Del Prato & Pylkkanen, 2014; Giglio et al., 2024b; Goldman et al., 2023; Habets et al., 2008; Murphy et al., 2022).

IFG engagement occurred earlier for noun inflections than for constituent assembly, suggesting its critical role for inflections (Sahin et al., 2006; Shapiro et al., 2006). A similarity across both noun morphosyntax conditions was relatively prolonged IFG response (Figure 5-7), which suggests that this region is likely engaged in a process that is common across both conditions, such as linearization of morphemes (Hagoort, 2005; Matchin & Hickok, 2020) and phonological planning/syllabification (Carota et al., 2023; Indefrey, 2011; Sahin et al., 2009).

The M1 response for constituent assembly was also earlier and relatively stronger than for noun inflection (Figure 7), likely due to the higher syllable length of constituent assembly compared to noun inflection response (Roelofs, 1997).

The present study adds to existing knowledge of noun morphosyntax in several ways. It identifies distinct neural patterns to noun inflection and noun constituent assembly. One salient difference is the timeline of IFG and PTL engagement, consistent with these regions’ roles in inflection and constituent assembly respectively. The second difference is that the neural response to constituent assembly occurred earlier. Both morphosyntax conditions showed simultaneous and prolonged engagement of multiple ROIs, which was particularly noticeable for the IFG. A noteworthy finding is the IPL response for both conditions, given that this region has not been frequently associated with morphosyntax (but see Gronering & Corina, 2023 for its role in syntactic processing). There was early engagement of M1 in both conditions, although the stronger M1 engagement for constituent assembly may be a confound of utterance length.

#### 4.2.2. Verbs & Sentences

The two verb-based morphosyntactic conditions elicited sentences (pronoun + tense marked verb, with and without an inflectional suffix). These two conditions were the most neurocomputationally challenging, as indicated by their significantly slower speech onset latencies than other conditions (Supplementary Figure S3), and were the only conditions that showed robust neural responses in the speech onset locked analyses (relative to verb naming).

There were several remarkable similarities in the neural response to the two sentence conditions. The earliest region to engage for sentences (compared to verb naming) was the PTL (320-325 ms following the *condition icon*). As mentioned previously, the PTL was also the first neural region to engage for noun constituent assembly (at 255 ms post *condition* icon). The early PTL response across both verb and noun constituent assembly conditions confirms that the PTL plays a critical role in syntactic planning (Giglio et al., 2024a; Lee et al., 2018; Matchin & Hickok, 2020).

The IFG response timeline was also similar across the two conditions (starting at 300 and 325 ms after the *condition* icon, Supplementary Table S3). Verb naming (> verb past and > verb future) also showed an IFG response at this time (Table 5, Supplementary Table S3). The similarity in IFG response across the three verb conditions suggests that verbs inherently activate syntactic representations irrespective of task demands.

Across both sentence conditions, the ATL response emerged relatively late (535 ms and 665 ms after the *condition* icon) compared to the timeline of ATL engagement for noun morphosyntax (275 ms and 445 ms, Table 2). The late ATL response is unsurprising given the lack of evidence for its role in morphosyntactic planning (Giglio et al., 2024; Wilson et al., 2014). This late ATL response possibly reflects a post-syntactic re-confirmation of the concept being expressed. The noun morphosyntax conditions, in contrast, were more reliant on conceptual combination (color, number) resulting in earlier ATL engagement (Pylkkanen, 2020).

Differences between the two verb morphosyntax conditions primarily showed a stronger spatiotemporal response for future tense compared to past tense across all five ROIs (Supplementary Table S3, see also Table 7 & Figure 10). One might hypothesize that the neural response for language planning would increase in magnitude with each morphosyntactic computation (constituent assembly + inflection), however the present findings do not support this linearly additive view^5^. Instead, the stronger neural response for future tense is likely a consequence of its significantly lower frequency of usage in English compared to simple past tense (Askarian, 2026; Wang & Zhu, 2023) given that lower frequency linguistic units require more cognitive effort and involve more widespread neural activity to produce (Berglund-Barraza et al., 2019; Schuster et al., 2025).

In summary, the present study showed that verb-related morphosyntactic planning engages a widespread neural response that is similar across the two morphosyntactic conditions in terms of the timeline of PTL, IFG and ATL activity. PTL and IFG were engaged earlier and almost simultaneously, while the ATL responded much later. Future tense showed a stronger neural response than past tense, presumably due to the higher cognitive effort needed to plan the less frequent future tense. There was no evidence of an incremental increase in neural response when both inflection and constituent assembly were planned compared to constituent assembly alone.

### 4.3. A spatiotemporal map of language production

Figure 13 summarizes the spatiotemporal patterns found across the six experimental conditions of the present study, based on the statistical threshold reporting criteria used in the Results section. This spatiotemporal map of noun and verb lexical and morphosyntactic production goes beyond the existing spatiotemporal model of noun production which was based on a meta analysis of chronometric data from reaction time and electrophysiologic studies and neuroimaging findings (Indefrey, 2011; Indefrey & Levelt, 2004). In proposing their noun production model, Indefrey and Levelt (2004) cautioned against a strict interpretation of their proposed timelines because these were estimated from inherently different experimental paradigms (p.7). In terms of morphosyntax, this map extends the anatomical model of Matchin and Hickok (2020) and provides timelines.

**Figure 13.**
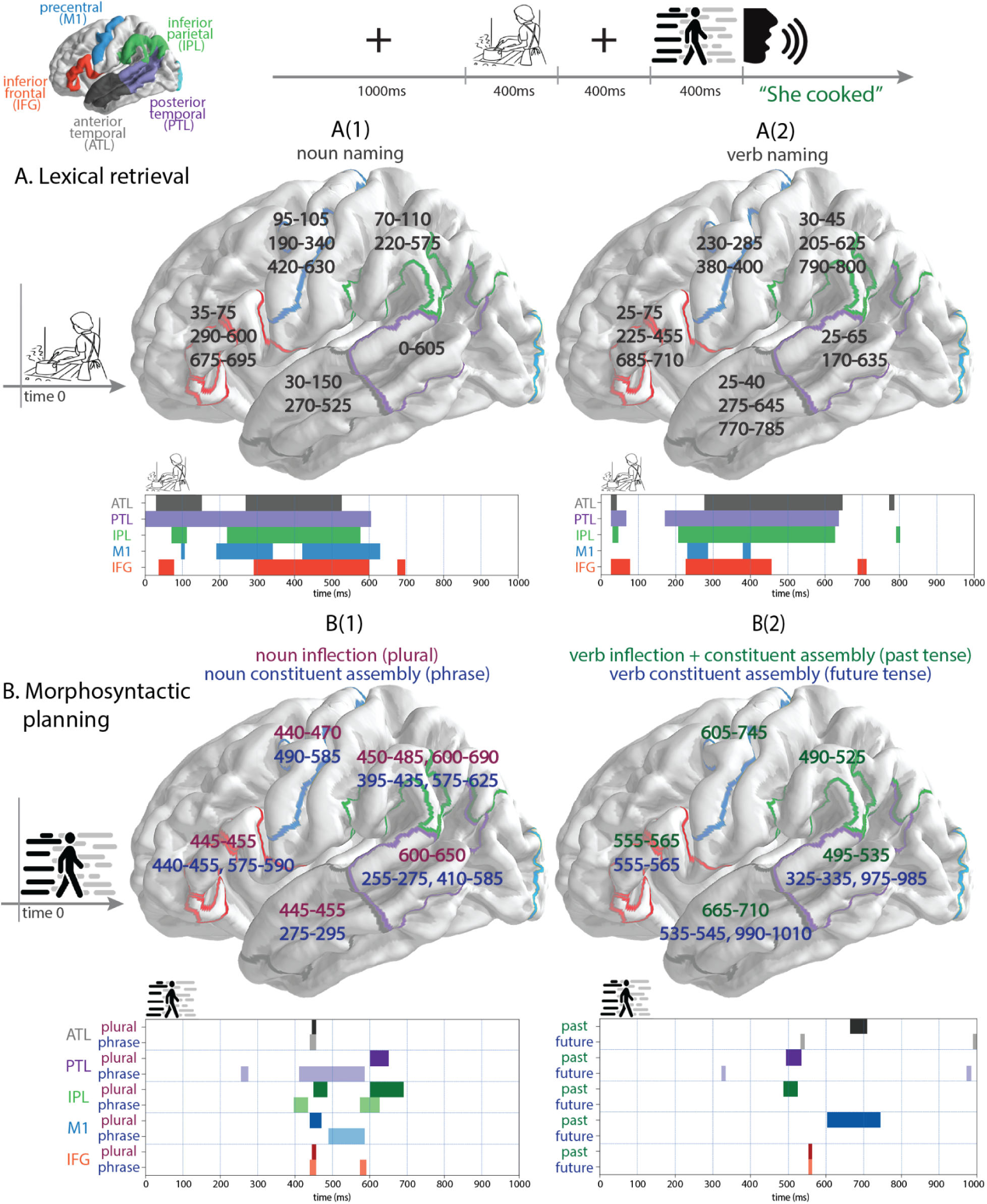
Summary of the spatiotemporal response to lexical retrieval (A) and morphosyntactic planning (B) showing the response timelines for the five language ROIs. The starting time for lexical retrieval (A) is picture presentation, while the starting time for morphosyntactic planning (B) is the presentation of the condition icon (see Figure 1). Response timelines are presented both as milliseconds overlaying with ROIs, and in the timeline chart below each anatomical plot. The results for lexical retrieval are based on one-sample vector field T^2^ tests and for morphosyntactic planning are based on paired comparisons with lexical retrieval. The time ranges are thresholded at 30% significant voxels within each ROI, a minimum significant duration > 10 ms, and a minimum average *T*^2^ > 4. Any two significant activation time ranges that are less than 75 ms apart were combined. ROIs are color coded, ATL is gray, PTL is purple, IPL is green, M1 is dark blue, and IFG is red.

This spatiotemporal map does not aim to assign specific linguistic computations to specific brain regions (e.g., syntax = IFG); rather it captures the temporal choreography across the cortical ROIs. It is evident that all ROIs are engaged in lexical and morphosyntactic planning for production, and either re-engage multiple times or continue in an active state for a prolonged period. This is consistent with multiple sources of evidence that point to a network model of language in which the syntactic, semantic, and phonological properties of a word are bound to a single anatomically distributed representation. It has been proposed that during language production, this representation is initially activated as a whole, which is followed by task relevant operations that flexibly engage different ROIs (Fairs et al., 2021; Miozzo et al., 2015; Pickering & Strijkers, 2024). Moving away from localized centers for specific operations (e.g., syntax), a network view proposes distributed but relatively specialized subnetworks. Dynamic interactions between these subnetworks give rise to a succession of new equilibrium states (see for example Duffau, 2018; Herbert & Duffau, 2020). Neural oscillations and their coupling across brain regions have been proposed as the mechanism underlying the language network (Benitez-Burraco & Murphy, 2019; Piai & Zheng, 2019). More work will be needed to integrate the findings of this study with neural oscillations across experimental conditions and also the role of subcortical regions in language production (Murphy et al., 2021; Turker et al., 2023) to develop a neurophysiological model of language production.

### 4.4. Limitations

There were some inherent confounds across the experimental conditions. While the study sought to compare morphosyntactic planning across lexical categories, this added a layer of conceptual and motoric differences across the conditions. The noun morphosyntax conditions likely involved numerosity processes (noun plural) for noun inflection and implausibility (few objects are blue in the real world) for noun constituent assembly. Similarly, there were differences in verb tense across the two verb morphosyntax/sentence conditions, and it has been suggested that the depth of cognitive simulation of action events could differ across verb tenses (Bergen & Wheeler, 2010; Gennari, 2004). In regards to motor planning, the length of the verbal response (number of syllables) was different across experimental conditions. In particular, the conditions with constituent assembly resulted in multiword responses compared to the picture naming and noun inflection. As discussed in section 4.2., an effect of syllable length was evident on the M1 and IFG responses (Supplementary Table S4 and Figure S3). However, the distinct timelines of the syllable length effect and the morphosyntactic responses of IFG and M1 reported in Tables 3-5 gives assurance that the morphosyntactic effects are not reducible to differences in the number of syllables.

While the elicitation of verbal responses increased the ecological validity of the experimental design for studying overt sentence production, the study did not completely delineate speech motor planning from linguistic planning. The study initially included a nonlinguistic verbal production condition, intended to elicit speech planning in the absence of linguistic processing. This involved a *condition* icon that indicated participants should say “blah blah” in response to the same action and object picture stimuli (for comparable visual processing). However, this condition resulted in stronger brain responses in language ROIs compared to the verb/noun naming conditions. This may be because producing an atypical verbal response demands additional inhibitory control to suppress the automatic lexical access of the picture names (cf. Navarrete & Costa, 2005). Nevertheless, the inclusion of M1 as an ROI, together with the supplementary analysis of syllable length effects, supports the interpretation that the linguistic planning effects we observed in this study are not primarily driven by speech motor planning.

### 4.4. Conclusions

The current study systematically investigated the spatiotemporal dynamics of lexical and morphosyntactic planning for language production, addressing a significant gap in current understanding of how the human brain formulates sentences. Using MEG and a novel overt picture naming paradigm combined with EMG to track speech-related movement artifacts, the present study examined the brain activity in key left hemisphere ROIs (ATL, IFG, IPL, M1, and PTL) across a two-second time course. The findings support a network view of language production with simultaneous ignition and flexible reverberation of ROIs (Fairs et al., 2021; Pickering & Strijkoff, 2024; Miozzo et al., 2015). This study also found early engagement of M1, suggesting that feed-forward action of motor plans occurs concurrently with linguistic planning (Miller & Guenther, 2021). In spite of recruitment of all ROIs, this study also found unique neural patterns across the two lexical categories, and also for each morphosyntactic operation (Faroqi-Shah et al., 2018; Liljestrom et al., 2009). In conclusion, the present study provides a much more nuanced spatiotemporal map of lexical and morphosyntactic production than previously known (Indefrey, 2011; Matchin & Hickok, 2020). .

## Acknowledgements

This research was funded by University of Maryland, College Park, and by National Institutes of Health NIDCD R01DC020483A. We thank Ciaran Stone for assisting with programming the experiment and data collection.

## CRediT author statement

Yi Wei: Software, Writing - original draft, Formal Analysis, Data Curation, Visualization

L. Robert Slevc: Methodology, Writing - review and editing, Funding acquisition

Christian Brodbeck: Conceptualization, Methodology, Software, Writing - review and editing, Resources or Supervision, Funding acquisition

Yasmeen Faroqi-Shah: Conceptualization, Methodology, Writing -original draft, Supervision, Project administration, Funding acquisition

## Footnotes

1 The presentation sequence of the two trial sequences, picture-preceding-icon and icon-preceding-picture, was counterbalanced across participants.

2 Presentation scripts introduce an approximately 10 ms delay for each image presentation due to the screen refresh rate, which was not accounted for in our stimuli locked analysis, but was accounted for in our speech onset locked analysis.

3 In addition to the 300 critical trials, there were 100 trials of a non-speech condition (“blah blah”), which is not reported in this study.

4 The timelines in the text refer to the time elapsed since the *condition* icon whereas the timescale in the Figures refers to the start of the experimental trial

5 This interpretation is being made in the context of present study’s experimental conditions. Other syntactically more complex sentences such as those with embedded clauses or objective relatives could very well show an additive effect of computations on the neural response.

