## Supplemental materials for "Spatiotemporal dynamics of lexical and morphosyntactic planning during language production revealed by magnetoencephalography"

### Supplementary Materials

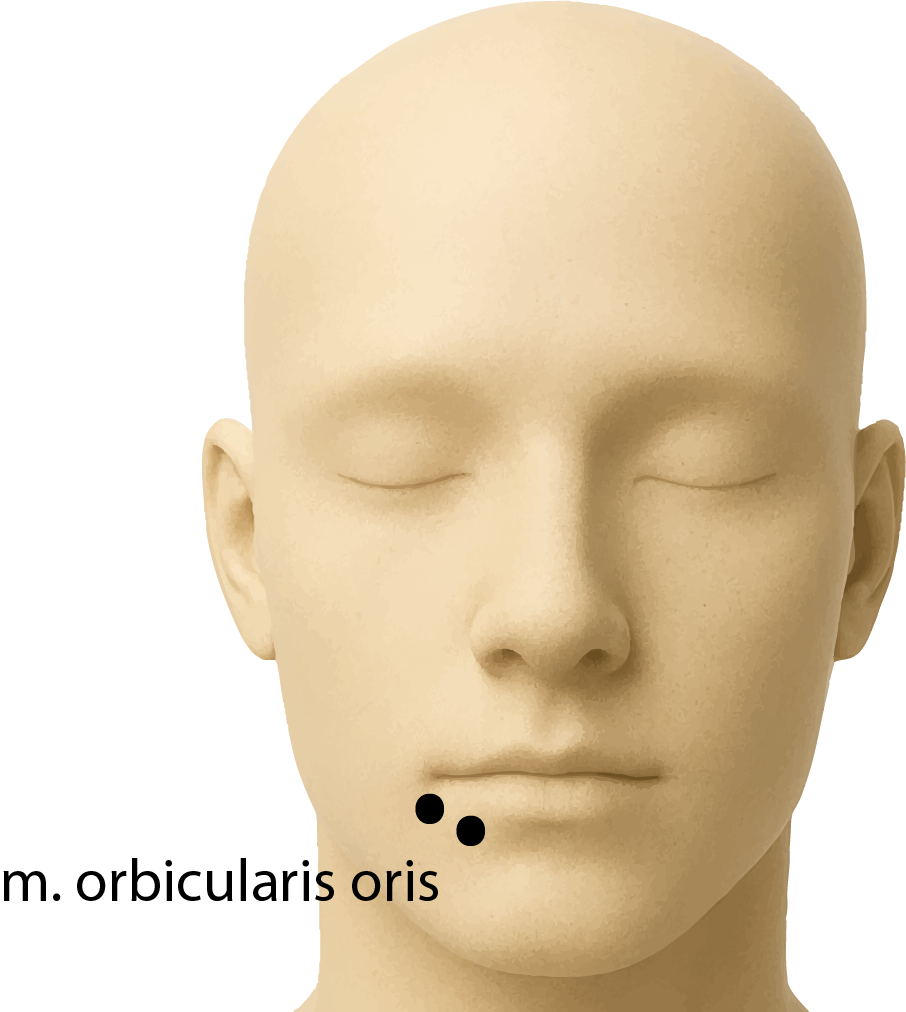

**Supplementary Figure S1.** EMG sensor placement. Two active electrodes were placed below the right lip approximately 1cm apart from each other to capture muscle activities from the *m. orbicularis oris*. A third (ground) sensor was placed on the right clavicle near the shoulder.

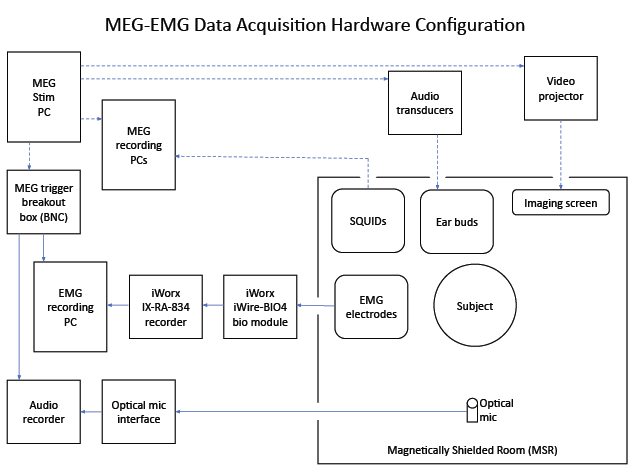

**Supplementary Figure S2.** Hardware setup for experiment presentation, and recording of MEG, EMG and audio responses.

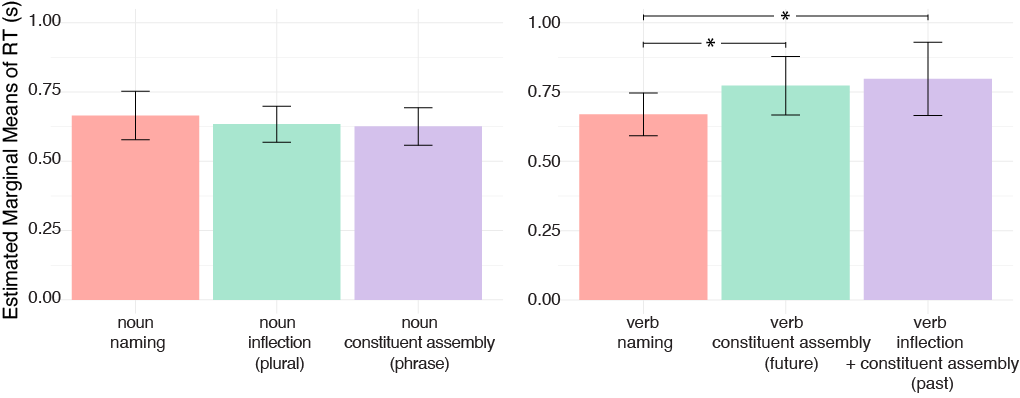

**Supplementary Figure S3.** Plot of speech onset times (measured from the onset of the speak icon to verbal response) across the six experimental conditions.

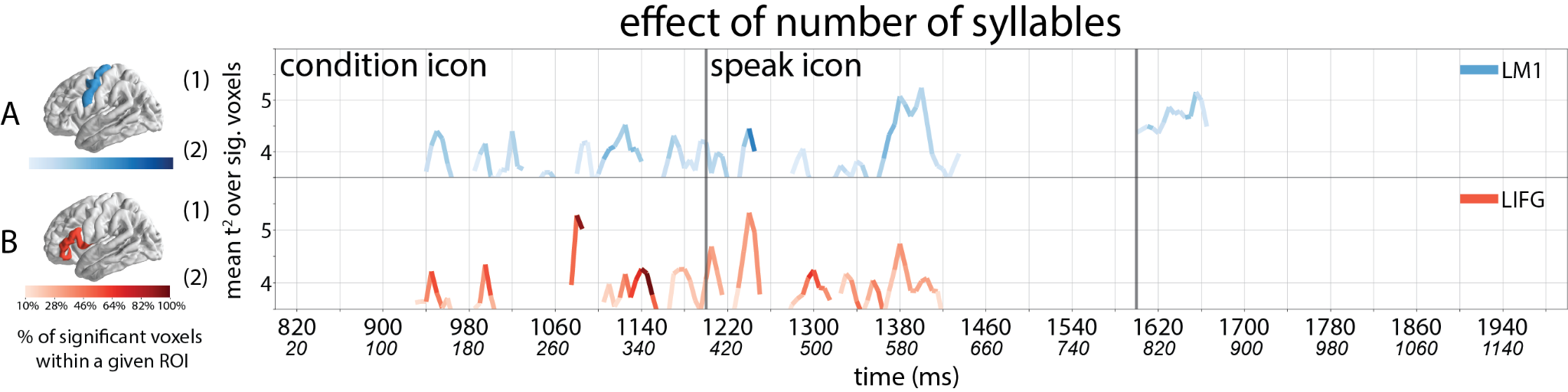

**Supplementary Figure S4.** Results of a two-stage regression analysis of MEG source-level responses, assessing the effect of syllable number from selected ROIs. Line traces indicate time points at which the group-level Hotelling*T^2^*-test revealed a significant effect of syllable number from M1 (A) and IFG (B). The color saturation of the line traces reflects the percentage of significant voxels. The timeline of significant M1 and IFG responses is listed in Supplementary Table 4.

**Supplementary Table S1a.** Results for the one-sample vector field T^2^ tests for noun and verb naming. These results are thresholded at 30% significant voxels within each ROI, a minimum significant duration > 10ms, minimum T^2^ > 4. The starting time is the picture presentation (Figure 1). ATL- anterior temporal lobe, PTL - posterior temporal lobe, IPL - inferior parietal lobe, M1 - primary motor cortex, and IFG - inferior frontal gyrus.

| **LH Regions** | **NOUN NAMING** | | **VERB NAMING** |
| --- | --- | --- | --- |
| ATL | 30 - 45ms (max $\underline{t^{2}}$ = 4.68)  55 - 65ms (max $\underline{t^{2}}$ = 4.23)  80 - 110ms (max $\underline{t^{2}}$ = 5.99)  140 - 150ms (max $\underline{t^{2}}$ = 4.61)  270 - 475ms (max $\underline{t^{2}}$ = 6.06)  515 - 525ms (max $\underline{t^{2}}$ = 4.29) | | 25 - 40ms (max $\underline{t^{2}}$ = 7.46)  275 - 510ms (max $\underline{t^{2}}$ = 6.97)  565 - 585ms (max $\underline{t^{2}}$ = 4.29)  610 - 645ms (max $\underline{t^{2}}$ = 4.58)  770 - 785ms (max $\underline{t^{2}}$ = 4.73) |
| PTL | 0 - 105ms (max $\underline{t^{2}}$ = 5.46)  140 - 150ms (max $\underline{t^{2}}$ = 4.2)  170 - 180ms (max $\underline{t^{2}}$ = 5.19)  235 - 245ms (max $\underline{t^{2}}$ = 4.12)  260 - 605ms (max $\underline{t^{2}}$ = 8.24) | | 25 - 65ms (max $\underline{t^{2}}$ = 5.88)  170 - 180ms (max $\underline{t^{2}}$ = 4.92)  205 - 250ms (max $\underline{t^{2}}$ = 4.65)  310 - 635ms (max $\underline{t^{2}}$ = 7.63) |
| IPL | 70 - 110ms (max $\underline{t^{2}}$ = 5.16)  220 - 270ms (max $\underline{t^{2}}$ = 4.67)  330 - 350ms (max $\underline{t^{2}}$ = 4.46)  390 - 435ms (max $\underline{t^{2}}$ = 4.87)  450 - 490ms (max $\underline{t^{2}}$ = 5.17)  500 - 510ms (max $\underline{t^{2}}$ = 4.27)  535 - 575ms (max $\underline{t^{2}}$ = 4.48) | | 30 - 45ms (max $\underline{t^{2}}$ = 4.8)  205 - 230ms (max $\underline{t^{2}}$ = 5.52)  265 - 275ms (max $\underline{t^{2}}$ = 4.49)  295 - 325ms (max $\underline{t^{2}}$ = 4.3)  335 - 455ms (max $\underline{t^{2}}$ = 5.59)  470 - 490ms (max $\underline{t^{2}}$ = 4.7)  500 - 515ms (max $\underline{t^{2}}$ = 4.99)  535 - 580ms (max $\underline{t^{2}}$ = 5.2)  595 - 625ms (max $\underline{t^{2}}$ = 4.5)  790 - 800ms (max $\underline{t^{2}}$ = 4.22) |
| M1 | 95 - 105ms (max $\underline{t^{2}}$ = 5.1)  190 - 290ms (max $\underline{t^{2}}$ = 4.75)  325 - 340ms (max $\underline{t^{2}}$ = 4.3)  420 - 490ms (max $\underline{t^{2}}$ = 5.72)  500 - 510ms (max $\underline{t^{2}}$ = 4.5)  520 - 595ms (max $\underline{t^{2}}$ = 4.55)  610 - 630ms (max $\underline{t^{2}}$ = 4.51) | | 230 - 250ms (max $\underline{t^{2}}$ = 4.62)  265 - 285ms (max $\underline{t^{2}}$ = 4.88)  380 - 400ms (max $\underline{t^{2}}$ = 4.55) |
| IFG | 35 - 55ms (max $\underline{t^{2}}$ = 5.02)  65 - 75ms (max $\underline{t^{2}}$ = 4.52)  290 - 300ms (max $\underline{t^{2}}$ = 4.5)  325 - 485ms (max $\underline{t^{2}}$ = 6.17)  505 - 525ms (max $\underline{t^{2}}$ = 4.38)  545 - 600ms (max $\underline{t^{2}}$ = 5.46)  675 - 695ms (max $\underline{t^{2}}$ = 4.61) | | 25 - 35ms (max $\underline{t^{2}}$ = 4.88)  50 - 75ms (max $\underline{t^{2}}$ = 5.14)  225 - 290ms (max $\underline{t^{2}}$ = 4.62)  300 - 310ms (max $\underline{t^{2}}$ = 4.23)  330 - 360ms (max $\underline{t^{2}}$ = 4.56)  385 - 430ms (max $\underline{t^{2}}$ = 5.68)  440 - 455ms (max $\underline{t^{2}}$ = 4.74)  685 - 710ms (max $\underline{t^{2}}$ = 5.06) |

**Supplementary Table S1b.** All significant results for the one-sample vector field T^2^ tests for noun and verb naming. The starting time is the picture presentation (Figure 1). ATL- anterior temporal lobe, PTL - posterior temporal lobe, IPL - inferior parietal lobe, M1 - primary motor cortex, and IFG - inferior frontal gyrus.

| **LH Regions** | **NOUN NAMING** | | **VERB NAMING** |
| --- | --- | --- | --- |
| ATL | 0 - 10ms (max $\underline{t^{2}}$ = 3.72)  30 - 155ms (max $\underline{t^{2}}$ = 5.99)  230 - 240ms (max $\underline{t^{2}}$ = 3.7)  255 - 600ms (max $\underline{t^{2}}$ = 6.06)  610 - 630ms (max $\underline{t^{2}}$ = 3.86)  645 - 665ms (max $\underline{t^{2}}$ = 3.56)  685 - 695ms (max $\underline{t^{2}}$ = 4.17)  795 - 800ms (max $\underline{t^{2}}$ = 3.19) | | 10 - 50ms (max $\underline{t^{2}}$ = 7.46)  60 - 85ms (max $\underline{t^{2}}$ = 4.67)  115 - 120ms (max $\underline{t^{2}}$ = 3.82)  165 - 195ms (max $\underline{t^{2}}$ = 4.33)  210 - 515ms (max $\underline{t^{2}}$ = 6.97)  530 - 725ms (max $\underline{t^{2}}$ = 4.58)  740 - 805ms (max $\underline{t^{2}}$ = 4.73) |
| PTL | 0 - 115ms (max $\underline{t^{2}}$ = 5.46)  135 - 190ms (max $\underline{t^{2}}$ = 5.19)  215 - 660ms (max $\underline{t^{2}}$ = 8.24)  845 - 950ms (max $\underline{t^{2}}$ = 4.85) | | 0 - 120ms (max $\underline{t^{2}}$ = 5.88)  145 - 155ms (max $\underline{t^{2}}$ = 3.71)  165 - 195ms (max $\underline{t^{2}}$ = 4.92)  205 - 250ms (max $\underline{t^{2}}$ = 4.65)  285 - 700ms (max $\underline{t^{2}}$ = 7.63)  735 - 745ms (max $\underline{t^{2}}$ = 4.06) |
| IPL | 0 - 5ms (max $\underline{t^{2}}$ = 4.29)  15 - 165ms (max $\underline{t^{2}}$ = 5.16)  190 - 490ms (max $\underline{t^{2}}$ = 5.17)  500 - 625ms (max $\underline{t^{2}}$ = 4.48)  640 - 655ms (max $\underline{t^{2}}$ = 3.87)  710 - 800ms (max $\underline{t^{2}}$ = 3.95) | | 25 - 70ms (max $\underline{t^{2}}$ = 4.8)  95 - 150ms (max $\underline{t^{2}}$ = 4.32)  205 - 250ms (max $\underline{t^{2}}$ = 5.52)  260 - 635ms (max $\underline{t^{2}}$ = 5.59)  650 - 700ms (max $\underline{t^{2}}$ = 3.7)  720 - 730ms (max $\underline{t^{2}}$ = 3.58)  740 - 845ms (max $\underline{t^{2}}$ = 4.22) |
| M1 | 65 - 110ms (max $\underline{t^{2}}$ = 5.1)  130 - 310ms (max $\underline{t^{2}}$ = 4.75)  325 - 510ms (max $\underline{t^{2}}$ = 5.72)  520 - 695ms (max $\underline{t^{2}}$ = 4.55)  725 - 800ms (max $\underline{t^{2}}$ = 3.89) | | 110 - 120ms (max $\underline{t^{2}}$ = 3.97)  150 - 180ms (max $\underline{t^{2}}$ = 4.47)  195 - 200ms (max $\underline{t^{2}}$ = 3.19)  220 - 320ms (max $\underline{t^{2}}$ = 4.88)  345 - 370ms (max $\underline{t^{2}}$ = 3.49)  380 - 405ms (max $\underline{t^{2}}$ = 4.55)  575 - 610ms (max $\underline{t^{2}}$ = 3.73)  630 - 640ms (max $\underline{t^{2}}$ = 3.81)  670 - 680ms (max $\underline{t^{2}}$ = 3.36)  720 - 725ms (max $\underline{t^{2}}$ = 3.49)  775 - 780ms (max $\underline{t^{2}}$ = 3.68) |
| IFG | 0 - 105ms (max $\underline{t^{2}}$ = 5.02)  170 - 175ms (max $\underline{t^{2}}$ = 3.4)  205 - 225ms (max $\underline{t^{2}}$ = 3.46)  240 - 310ms (max $\underline{t^{2}}$ = 4.5)  320 - 610ms (max $\underline{t^{2}}$ = 6.17)  620 - 705ms (max $\underline{t^{2}}$ = 4.61)  730 - 750ms (max $\underline{t^{2}}$ = 3.61)  775 - 780ms (max $\underline{t^{2}}$ = 3.18) | | 0 - 5ms (max $\underline{t^{2}}$ = 3.38)  15 - 80ms (max $\underline{t^{2}}$ = 5.14)  155 - 175ms (max $\underline{t^{2}}$ = 3.57)  195 - 200ms (max $\underline{t^{2}}$ = 3.56)  215 - 520ms (max $\underline{t^{2}}$ = 5.68)  530 - 565ms (max $\underline{t^{2}}$ = 3.95)  580 - 610ms (max $\underline{t^{2}}$ = 3.91)  680 - 735ms (max $\underline{t^{2}}$ = 5.06)  770 - 785ms (max $\underline{t^{2}}$ = 3.83) |

**Supplementary Table S2.** All significant results comparing noun and verb naming contrasts across the five left hemisphere regions of interest (time locked to stimuli onset). ATL- anterior temporal lobe, PTL - posterior temporal lobe, IPL - inferior parietal lobe, M1 - primary motor cortex, and IFG - inferior frontal gyrus.

| **LH Regions** | **0-400ms (picture)** | **400-800ms**  **(fixation cross)** | **800-1200 ms (condition icon)** | **1200 -1600 ms (speak icon)** | **>1600ms** |
| --- | --- | --- | --- | --- | --- |
| **VERB NAMING > NOUN NAMING (Figure 5a)** | | | | | |
| ATL | 170 - 215ms (max $\underline{t^{2}}$ = 4.12)  295 - 305ms (max $\underline{t^{2}}$ = 3.69)  320 - 405ms (max $\underline{t^{2}}$ = 6.84) | 630 - 650ms (max $\underline{t^{2}}$ = 3.58)  660 - 810ms (max $\underline{t^{2}}$ = 5.41) | 825 - 845ms (max $\underline{t^{2}}$ = 5.43)  855 - 860ms (max $\underline{t^{2}}$ = 4.08)  1160 - 1180ms (max $\underline{t^{2}}$ = 4.53) | 1295 - 1315ms (max $\underline{t^{2}}$ = 3.9)  1330 - 1355ms (max $\underline{t^{2}}$ = 4.14) |  |
| PTL | 110 - 155ms (max $\underline{t^{2}}$ = 5.2)  190 - 195ms (max $\underline{t^{2}}$ = 3.53)  210 - 215ms (max $\underline{t^{2}}$ = 3.62) | 400 - 405ms (max $\underline{t^{2}}$ = 4.13)  545 - 575ms (max $\underline{t^{2}}$ = 4.33)  620 - 660ms (max $\underline{t^{2}}$ = 4.17)  670 - 705ms (max $\underline{t^{2}}$ = 4.88)  730 - 740ms (max $\underline{t^{2}}$ = 3.28) | 820 - 840ms (max $\underline{t^{2}}$ = 4.06)  1080 - 1110ms (max $\underline{t^{2}}$ = 4.42) | 1225 - 1235ms (max $\underline{t^{2}}$ = 3.93)  1300 - 1305ms (max $\underline{t^{2}}$ = 3.89)  1320 - 1360ms (max $\underline{t^{2}}$ = 4.17)  1510 - 1515ms (max $\underline{t^{2}}$ = 4.7) |  |
| IPL | 110 - 145ms (max $\underline{t^{2}}$ = 4.68)  370 - 430ms (max $\underline{t^{2}}$ = 4.01) | 505 - 520ms (max $\underline{t^{2}}$ = 4.07)  625 - 630ms (max $\underline{t^{2}}$ = 3.76)  655 - 685ms (max $\underline{t^{2}}$ = 4.02)  695 - 750ms (max $\underline{t^{2}}$ = 4.67)  770 - 775ms (max $\underline{t^{2}}$ = 4.06)  790 - 795ms (max $\underline{t^{2}}$ = 4.59) | 820 - 845ms (max $\underline{t^{2}}$ = 4.93)  865 - 885ms (max $\underline{t^{2}}$ = 3.93)  1185 - 1225ms (max $\underline{t^{2}}$ = 4.1) | 1235 - 1280ms (max $\underline{t^{2}}$ = 4.63)  1320 - 1325ms (max $\underline{t^{2}}$ = 4.05)  1345 - 1360ms (max $\underline{t^{2}}$ = 3.81) |  |
| M1 | 235 - 250ms (max $\underline{t^{2}}$ = 4.18)  260 - 265ms (max $\underline{t^{2}}$ = 3.93) |  |  | 1210 - 1215ms (max $\underline{t^{2}}$ = 3.46)  1295 - 1300ms (max $\underline{t^{2}}$ = 3.61)  1405 - 1425ms (max $\underline{t^{2}}$ = 4.58)  1475 - 1495ms (max $\underline{t^{2}}$ = 4.89) |  |
| IFG | 15 - 20ms  (max $\underline{t^{2}}$ = 3.61)  170 - 175ms  (max $\underline{t^{2}}$ = 3.92)  195 - 200ms  (max $\underline{t^{2}}$ = 3.55)  215 - 255ms  (max $\underline{t^{2}}$ = 4.62)  340 - 345ms (max $\underline{t^{2}}$ = 3.79) | 400 - 410ms (max $\underline{t^{2}}$ = 4.14)  425 - 435ms (max $\underline{t^{2}}$ = 4.56)  450 - 485ms (max $\underline{t^{2}}$ = 4.8)  495 - 510ms (max $\underline{t^{2}}$ = 3.54)  675 - 730ms (max $\underline{t^{2}}$ = 5.33)  740 - 750ms (max $\underline{t^{2}}$ = 3.51)  770 - 775ms (max $\underline{t^{2}}$ = 3.79) | 1130 - 1170ms (max $\underline{t^{2}}$ = 5.52)  1375 - 1385ms (max $\underline{t^{2}}$ = 3.75) |  |  |
| **NOUN NAMING > VERB NAMING (Figure 4, panel (2)s)** | | | | | |
| ATL | 85 - 110ms (max $\underline{t^{2}}$ = 4.87)  145 - 155ms (max $\underline{t^{2}}$ = 3.96) | 405 - 470ms (max $\underline{t^{2}}$ = 4.56)  500 - 505ms (max $\underline{t^{2}}$ = 3.69) | 860 - 875ms (max $\underline{t^{2}}$ = 4.81) | 1315 - 1330ms (max $\underline{t^{2}}$ = 4.14)  1360 - 1400ms (max $\underline{t^{2}}$ = 4.55) |  |
| PTL | 105 - 110ms (max $\underline{t^{2}}$ = 3.73)  275 - 280ms (max $\underline{t^{2}}$ = 3.78)  335 - 345ms (max $\underline{t^{2}}$ = 3.7)  365 - 400ms (max $\underline{t^{2}}$ = 5.56) | 795 - 805ms (max $\underline{t^{2}}$ = 4.47) |  | 1215 - 1225ms (max $\underline{t^{2}}$ = 4.16)  1375 - 1380ms (max $\underline{t^{2}}$ = 3.61)  1505 - 1510ms (max $\underline{t^{2}}$ = 4.7) |  |
| IPL | 145 - 150ms (max $\underline{t^{2}}$ = 4.43)  210 - 220ms (max $\underline{t^{2}}$ = 3.84)  245 - 250ms (max $\underline{t^{2}}$ = 3.49) | 650 - 655ms (max $\underline{t^{2}}$ = 3.75) |  | 1280 - 1290ms (max $\underline{t^{2}}$ = 3.9)  1360 - 1370ms (max $\underline{t^{2}}$ = 3.85)  1405 - 1415ms (max $\underline{t^{2}}$ = 3.98) |  |
| M1 | 95 - 105ms (max $\underline{t^{2}}$ = 3.58)  130 - 140ms (max $\underline{t^{2}}$ = 3.96)  160 - 185ms (max $\underline{t^{2}}$ = 3.88) | 410 - 485ms (max $\underline{t^{2}}$ = 3.75)  520 - 535ms (max $\underline{t^{2}}$ = 3.99) | 990 - 995ms  (max $\underline{t^{2}}$ = 4.98)  1140 - 1150ms  (max $\underline{t^{2}}$ = 3.91)  1175 - 1185ms  (max $\underline{t^{2}}$ = 4.04)  1300 - 1305ms  (max $\underline{t^{2}}$ = 3.55)  1340 - 1345ms  (max $\underline{t^{2}}$ = 3.33) |  |  |
| IFG | 140 - 145ms (max $\underline{t^{2}}$ = 3.57)  210 - 215ms (max $\underline{t^{2}}$ = 4.1)  375 - 380ms (max $\underline{t^{2}}$ = 3.35) | 410 - 425ms (max $\underline{t^{2}}$ = 4.86)  565 - 580ms (max $\underline{t^{2}}$ = 4.35) | 850 - 870ms (max $\underline{t^{2}}$ = 3.99)  1170 - 1185ms (max $\underline{t^{2}}$ = 5.52) | 1495 - 1500ms (max $\underline{t^{2}}$ = 3.94)  1530 - 1535ms (max $\underline{t^{2}}$ = 4.07) |  |

**Supplementary Table S3.** All significant results for morphosyntactic processes across the five left hemisphere regions of interest (time locked to stimuli onset). ATL- anterior temporal lobe, PTL - posterior temporal lobe, IPL - inferior parietal lobe, M1 - primary motor cortex, and IFG - inferior frontal gyrus.

| **LH Regions** | **800-1200 ms (condition icon)** | **1200 -1600 ms (speak icon)** | **>1600ms** |
| --- | --- | --- | --- |
| **NOUN INFLECTION (PLURAL) > NOUN NAMING (Figure 5, panel (1)s)** | | | |
| Anterior temporal  (ATL) |  | 1175 - 1210ms (max $\underline{t^{2}}$ = 4.79)  1240 - 1265ms (max $\underline{t^{2}}$ = 4.56)  1285 - 1295ms (max $\underline{t^{2}}$ = 3.64)  1420 - 1430ms (max $\underline{t^{2}}$ = 3.88)  1440 - 1455ms (max $\underline{t^{2}}$ = 3.98) |  |
| Posterior temporal  (PTL) |  | 1360 - 1375ms (max $\underline{t^{2}}$ = 4.28)  1395 - 1455ms (max $\underline{t^{2}}$ = 5.56)  1465 - 1475ms (max $\underline{t^{2}}$ = 4.14) |  |
| Inferior parietal  (IPL) |  | 1250 - 1290ms (max $\underline{t^{2}}$ = 4.51)  1400 - 1490ms (max $\underline{t^{2}}$ = 4.96) |  |
| Precentral  (M1) |  | 1215 - 1225ms (max $\underline{t^{2}}$ = 4.48)  1240 - 1270ms (max $\underline{t^{2}}$ = 4.89)  1295 - 1305ms (max $\underline{t^{2}}$ = 4.14)  1455 - 1480ms (max $\underline{t^{2}}$ = 3.93)  1490 - 1510ms (max $\underline{t^{2}}$ = 4.04)  1560 - 1575ms (max $\underline{t^{2}}$ = 4.33) |  |
| Inferior frontal  (IFG) | 1190 - 1195ms (max $\underline{t^{2}}$ = 4.11) | 1240 - 1270ms (max $\underline{t^{2}}$ = 4.69)  1450 - 1460ms (max $\underline{t^{2}}$ = 4.63) |  |
| **NOUN NAMING > NOUN INFLECTION (PLURAL) (Figure 5, panel (2)s)** | | | |
| Anterior temporal  (ATL) |  | 1270 - 1285ms (max $\underline{t^{2}}$ = 4.11) |  |
| Precentral  (M1) | 1180 - 1190ms (max $\underline{t^{2}}$ = 3.81) | 1325 - 1345ms (max $\underline{t^{2}}$ = 4.41)  1575 - 1585ms (max $\underline{t^{2}}$ = 4.53) |  |
| **NOUN CONSTITUENT ASSEMBLY (PHRASE) > NOUN NAMING (Figure 6, panel (1)s)** | | | |
| Anterior temporal  (ATL) | 1075 - 1095ms (max $\underline{t^{2}}$ = 4.59) | 1205 - 1215ms (max $\underline{t^{2}}$ = 4.39) |  |
| Posterior temporal  (PTL) | 1055 - 1080ms (max $\underline{t^{2}}$ = 4.6) | 1210 - 1280ms (max $\underline{t^{2}}$ = 4.88)  1335 - 1385ms (max $\underline{t^{2}}$ = 5.17) |  |
| Inferior parietal  (IPL) | 1195 - 1205ms (max $\underline{t^{2}}$ = 4.9) | 1225 - 1240ms (max $\underline{t^{2}}$ = 4.39)  1375 - 1390ms (max $\underline{t^{2}}$ = 4.42)  1400 - 1435ms (max $\underline{t^{2}}$ = 4.59) |  |
| Precentral  (M1) |  | 1255 - 1270ms (max $\underline{t^{2}}$ = 3.85)  1285 - 1305ms (max $\underline{t^{2}}$ = 4.5)  1315 - 1345ms (max $\underline{t^{2}}$ = 4.77)  1360 - 1385ms (max $\underline{t^{2}}$ = 4.68) |  |
| Inferior frontal  (IFG) | 1075 - 1085ms (max $\underline{t^{2}}$ = 4.61) | 1240 - 1260ms (max $\underline{t^{2}}$ = 5.01)  1375 - 1390ms (max $\underline{t^{2}}$ = 5.12) |  |
| **NOUN NAMING > NOUN CONSTITUENT ASSEMBLY (PHRASE) (Figure 6, panel (2)s)** | | | |
| Anterior temporal  (ATL) |  | 1230 - 1260ms (max $\underline{t^{2}}$ = 4.92) |  |
| Parietal temporal  (PTL) | 1160 - 1170ms (max $\underline{t^{2}}$ = 4.1)  1185 - 1210ms (max $\underline{t^{2}}$ = 4.43) |  |  |
| Precentral  (M1) |  |  | 1605 - 1615ms (max $\underline{t^{2}}$ = 5.43) |
| **VERB CONSTITUENT ASSEMBLY (FUTURE) > VERB NAMING (Figure 7, panel (1)s)** | | | |
| Anterior temporal  (ATL) |  | 1330 - 1345ms (max $\underline{t^{2}}$ = 4.36)  1355 - 1360ms (max $\underline{t^{2}}$ = 4.38)  1505 - 1530ms (max $\underline{t^{2}}$ = 4.52) | 1790 - 1815ms (max $\underline{t^{2}}$ = 5.01) |
| Posterior temporal  (PTL) | 1100 - 1105ms (max $\underline{t^{2}}$ = 4.0)  1125 - 1140ms (max $\underline{t^{2}}$ = 4.46)  1180 - 1185ms (max $\underline{t^{2}}$ = 4.62) |  | 1640 - 1650ms (max $\underline{t^{2}}$ = 4.19)  1775 - 1785ms (max $\underline{t^{2}}$ = 4.56) |
| Inferior parietal  (IPL) | 1125 - 1130ms (max $\underline{t^{2}}$ = 4.59) | 1375 - 1385ms (max $\underline{t^{2}}$ = 3.98) |  |
| Precentral  (M1) |  |  | 1640 - 1660ms (max $\underline{t^{2}}$ = 4.37) |
| Inferior frontal  (IFG) |  | 1355 - 1365ms (max $\underline{t^{2}}$ = 4.51) |  |
| **VERB NAMING > VERB CONSTITUENT ASSEMBLY (FUTURE) (Figure 7, panel (2)s)** | | | |
| Anterior temporal  (ATL) |  | 1500 - 1505ms (max $\underline{t^{2}}$ = 4.71) |  |
| Anterior parietal  (PTL) |  | 1350 - 1355ms (max $\underline{t^{2}}$ = 4.39)  1395 - 1420ms (max $\underline{t^{2}}$ = 4.64) |  |
| Inferior parietal  (IPL) |  |  | 1635 - 1660ms (max $\underline{t^{2}}$ = 4.32)  1695 - 1710ms (max $\underline{t^{2}}$ = 4.18) |
| Precentral  (M1) |  | 1480 - 1490ms (max $\underline{t^{2}}$ = 4.56) |  |
| Inferior frontal  (IFG) | 1125 - 1150ms (max $\underline{t^{2}}$ = 4.44) |  |  |
| **VERB INFLECTION + CONSTITUENT ASSEMBLY (PAST) > VERB NAMING**  **(Figure 8, panel (1)s)** | | | |
| Anterior temporal  (ATL) | 975 - 985ms (max $\underline{t^{2}}$ = 4.47) | 1295 - 1300ms (max $\underline{t^{2}}$ = 3.82)  1415 - 1420ms (max $\underline{t^{2}}$ = 3.62)  1465 - 1475ms (max $\underline{t^{2}}$ = 4.35)  1495 - 1540ms (max $\underline{t^{2}}$ = 5.32)  1590 - 1600ms (max $\underline{t^{2}}$ = 3.65) | 1610 - 1620ms (max $\underline{t^{2}}$ = 4.65) |
| Posterior temporal  (PTL) | 980 - 990ms (max $\underline{t^{2}}$ = 4.07) | 1240 - 1250ms (max $\underline{t^{2}}$ = 4.07)  1295 - 1335ms (max $\underline{t^{2}}$ = 5.17)  1505 - 1510ms (max $\underline{t^{2}}$ = 3.64)  1535 - 1540ms (max $\underline{t^{2}}$ = 3.88) |  |
| Inferior parietal  (IPL) | 1015 - 1020ms (max $\underline{t^{2}}$ = 3.79) | 1290 - 1325ms (max $\underline{t^{2}}$ = 5.15)  1335 - 1350ms (max $\underline{t^{2}}$ = 4.08)  1445 - 1455ms (max $\underline{t^{2}}$ = 4.14) |  |
| Precentral  (M1) | 980 - 990ms (max $\underline{t^{2}}$ = 4.01)  1010 - 1015ms (max $\underline{t^{2}}$ = 3.89)  1130 - 1150ms (max $\underline{t^{2}}$ = 3.94) | 1340 - 1350ms (max $\underline{t^{2}}$ = 4.51)  1405 - 1435ms (max $\underline{t^{2}}$ = 5.14)  1455 - 1465ms (max $\underline{t^{2}}$ = 4.6)  1490 - 1500ms (max $\underline{t^{2}}$ = 3.96)  1530 - 1545ms (max $\underline{t^{2}}$ = 4.96)  1580 - 1585ms (max $\underline{t^{2}}$ = 4.17) | 1605 - 1610ms (max $\underline{t^{2}}$ = 3.93)  1635 - 1640ms (max $\underline{t^{2}}$ = 4.37) |
| Inferior frontal  (IFG) | 975 - 980ms (max $\underline{t^{2}}$ = 3.94)  1105 - 1115ms (max $\underline{t^{2}}$ = 4.46) | 1375 - 1385ms (max $\underline{t^{2}}$ = 4.31)  1455 - 1465ms (max $\underline{t^{2}}$ = 4.54) |  |
| **VERB NAMING > VERB INFLECTION + CONSTITUENT ASSEMBLY (PAST)**  **(Figure 8, panel (2)s)** | | | |
| Anterior temporal  (ATL) | 1110 - 1185ms (max $\underline{t^{2}}$ = 4.24)  1195 - 1205ms (max $\underline{t^{2}}$ = 3.45) | 1215 - 1225ms (max $\underline{t^{2}}$ = 3.6)  1420 - 1465ms (max $\underline{t^{2}}$ = 4.85)  1475 - 1495ms (max $\underline{t^{2}}$ = 4.35) | 1600 - 1610ms (max $\underline{t^{2}}$ = 4.65) |
| Parietal temporal  (PTL) | 1060 - 1070ms (max $\underline{t^{2}}$ = 4.08)  1085 - 1190ms (max $\underline{t^{2}}$ = 4.68) | 1335 - 1365ms (max $\underline{t^{2}}$ = 4.89)  1445 - 1460ms (max $\underline{t^{2}}$ = 4.29) |  |
| Inferior parietal  (IPL) | 1135 - 1185ms (max $\underline{t^{2}}$ = 3.98) | 1325 - 1335ms (max $\underline{t^{2}}$ = 4.83)  1360 - 1370ms (max $\underline{t^{2}}$ = 4.2)  1405 - 1420ms (max $\underline{t^{2}}$ = 3.92)  1455 - 1470ms (max $\underline{t^{2}}$ = 4.14)  1590 - 1595ms (max $\underline{t^{2}}$ = 3.98) | 1640 - 1650ms (max $\underline{t^{2}}$ = 3.81) |
| Precentral  (M1) |  | 1335 - 1340ms (max $\underline{t^{2}}$ = 4.51)  1480 - 1490ms (max $\underline{t^{2}}$ = 4.26)  1585 - 1605ms (max $\underline{t^{2}}$ = 4.17) | 1640 - 1650ms (max $\underline{t^{2}}$ = 4.37) |
| Inferior frontal  (IFG) | 980 - 985ms (max $\underline{t^{2}}$ = 3.94)  1115 - 1125ms (max $\underline{t^{2}}$ = 4.3)  1135 - 1145ms (max $\underline{t^{2}}$ = 3.8) | 1225 - 1230ms (max $\underline{t^{2}}$ = 3.7)  1450 - 1455ms (max $\underline{t^{2}}$ = 4.54)  1465 - 1470ms (max $\underline{t^{2}}$ = 3.83) |  |
| **NOUN CONSTITUENT ASSEMBLY (PHRASE) > NOUN INFLECTION (PLURAL)**  **(Figure 9, panel (1)s)** | | | |
| ATL | 1075 - 1090ms (max $\underline{t^{2}}$ = 4.48) | 1300 - 1305ms (max $\underline{t^{2}}$ = 3.8)  1325 - 1330ms (max $\underline{t^{2}}$ = 4.3) |  |
| Posterior temporal (PTL) | 940 - 950ms (max $\underline{t^{2}}$ = 3.86)  1055 - 1075ms (max $\underline{t^{2}}$ = 3.98) | 1220 - 1265ms (max $\underline{t^{2}}$ = 4.26)  1395 - 1400ms (max $\underline{t^{2}}$ = 4.11) |  |
| IPL | 1050 - 1070ms (max $\underline{t^{2}}$ = 4.39) | 1330 - 1345ms (max $\underline{t^{2}}$ = 4.01)  1395 - 1415ms (max $\underline{t^{2}}$ = 4.11) |  |
| M1 | 1015 - 1020ms (max $\underline{t^{2}}$ = 4.09)  1060 - 1075ms (max $\underline{t^{2}}$ = 4.67) | 1285 - 1300ms (max $\underline{t^{2}}$ = 5.15)  1330 - 1370ms (max $\underline{t^{2}}$ = 4.99) |  |
| IFG | 1080 - 1085ms (max $\underline{t^{2}}$ = 4.18)  1140 - 1150ms (max $\underline{t^{2}}$ = 3.86)  1175 - 1185ms (max $\underline{t^{2}}$ = 4.63) | 1240 - 1245ms (max $\underline{t^{2}}$ = 4.47)  1295 - 1300ms (max $\underline{t^{2}}$ = 4.42)  1385 - 1395ms (max $\underline{t^{2}}$ = 4.45) |  |
| **NOUN INFLECTION (PLURAL) > NOUN CONSTITUENT ASSEMBLY (PHRASE)**  **(Figure 9, panel (2)s)** | | | |
| Anterior temporal (ATL) | 1150 - 1245ms (max $\underline{t^{2}}$ = 5.6) | 1295 - 1300ms (max $\underline{t^{2}}$ = 3.91)  1385 - 1430ms (max $\underline{t^{2}}$ = 4.82)  1445 - 1475ms (max $\underline{t^{2}}$ = 4.29)  1490 - 1505ms (max $\underline{t^{2}}$ = 4.47) |  |
| Posterior temporal (PTL) | 1135 - 1185ms (max $\underline{t^{2}}$ = 4.37)  1195 - 1205ms (max $\underline{t^{2}}$ = 3.91) | 1400 - 1410ms (max $\underline{t^{2}}$ = 3.96) |  |
| Inferior parietal (IPL) | 1010 - 1025ms (max $\underline{t^{2}}$ = 4.51)  1045 - 1050ms (max $\underline{t^{2}}$ = 4.0)  1060 - 1065ms (max $\underline{t^{2}}$ = 4.39) | 1250 - 1300ms (max $\underline{t^{2}}$ = 4.42) |  |
| M1 |  | 1270 - 1285ms (max $\underline{t^{2}}$ = 5.15) |  |
| IFG |  | 1200 - 1205ms (max $\underline{t^{2}}$ = 4.06)  1225 - 1240ms (max $\underline{t^{2}}$ = 4.47)  1275 - 1295ms (max $\underline{t^{2}}$ = 4.42)  1395 - 1400ms (max $\underline{t^{2}}$ = 4.61)  1410 - 1425ms (max $\underline{t^{2}}$ = 5.46)  1450 - 1460ms (max $\underline{t^{2}}$ = 3.88)  1490 - 1505ms (max $\underline{t^{2}}$ = 3.87) |  |
| **VERB INFLECTION + CONSTITUENT ASSEMBLY (PAST) >**  **VERB CONSTITUENT ASSEMBLY (FUTURE)**  **(Figure 10, panel (1)s)** | | | |
| Posterior temporal (PTL) | 970 - 995ms (max $\underline{t^{2}}$ = 5.81) | 1250 - 1255ms (max $\underline{t^{2}}$ = 4.83)  1535 - 1540ms (max $\underline{t^{2}}$= 4.11) |  |
| Inferior frontal  (IFG) | 1125 - 1130ms (max $\underline{t^{2}}$ = 4.24) |  |  |
| **VERB CONSTITUENT ASSEMBLY (FUTURE) > VERB INFLECTION + CONSTITUENT ASSEMBLY (PAST)**  **(Figure 10, panel (2)s)** | | | |
| Anterior temporal (ATL) | 1085 - 1090ms (max $\underline{t^{2}}$ = 4.04)  1120 - 1130ms (max $\underline{t^{2}}$ = 4.02)  1150 - 1155ms (max $\underline{t^{2}}$ = 3.67) | 1210 - 1215ms (max $\underline{t^{2}}$ = 4.73)  1370 - 1375ms (max $\underline{t^{2}}$ = 4.8) |  |
| Posterior temporal (PTL) | 1055 - 1210ms (max $\underline{t^{2}}$ = 5.31) | 1240 - 1250ms (max $\underline{t^{2}}$ = 4.83)  1430 - 1445ms (max $\underline{t^{2}}$ = 4.23)  1465 - 1485ms (max $\underline{t^{2}}$ = 5.16)  1500 - 1515ms (max $\underline{t^{2}}$ = 6.42) |  |
| Inferior parietal  (IPL) | 1115 - 1190ms (max $\underline{t^{2}}$ = 4.53) | 1465 - 1505ms (max $\underline{t^{2}}$ = 6.05) |  |
| Precentral  (M1) | 1120 - 1140ms (max $\underline{t^{2}}$ = 5.21) |  |  |
| Inferior frontal  (IFG) | 1100 - 1105ms (max $\underline{t^{2}}$ = 4.34) |  |  |

**Supplementary Table S4.** All significant results for morphosyntactic planning time-locked to speech onset, across the five left hemisphere regions of interest (time locked to stimuli onset)(ATL- anterior temporal lobe, PTL - posterior temporal lobe, IPL - inferior parietal lobe, M1 - primary motor cortex, and IFG - inferior frontal gyrus).

| **LH Regions** | **-400 - 0ms (before speech onset)** | **0 - 400ms (after speech onset)** |
| --- | --- | --- |
| **VERB NAMING > NOUN NAMING** | | |
| ATL |  | 10 - 15ms (max $\underline{t^{2}}$ = 4.67) |
| **VERB CONSTITUENT ASSEMBLY (FUTURE) > VERB NAMING (Figure 9, panel (1)s)** | | |
| ATL | -20 - 135ms (max $\underline{t^{2}}$ = 6.59) | 260 - 390ms (max $\underline{t^{2}}$ = 5.09)  425 - 435ms (max $\underline{t^{2}}$ = 4.06) |
| PTL |  | 25 - 50ms (max $\underline{t^{2}}$ = 4.88)  60 - 145ms (max $\underline{t^{2}}$ = 5.9)  280 - 305ms (max $\underline{t^{2}}$ = 4.26)  330 - 335ms (max $\underline{t^{2}}$ = 4.55)  355 - 360ms (max $\underline{t^{2}}$ = 3.67)  385 - 390ms (max $\underline{t^{2}}$ = 4.11) |
| M1 |  | 20 - 25ms (max $\underline{t^{2}}$ = 4.09)  85 - 120ms (max $\underline{t^{2}}$ = 4.52)  360 - 375ms (max $\underline{t^{2}}$ = 4.74) |
| IFG | -25 - -20ms (max $\underline{t^{2}}$ = 4.29) | 5 - 75ms (max $\underline{t^{2}}$ = 5.83)  95 - 105ms (max $\underline{t^{2}}$ = 4.3)  180 - 190ms (max $\underline{t^{2}}$ = 3.97)  220 - 245ms (max $\underline{t^{2}}$ = 4.32)  255 - 260ms (max $\underline{t^{2}}$ = 3.83)  280 - 290ms (max $\underline{t^{2}}$ = 3.99) |
| **VERB NAMING > VERB CONSTITUENT ASSEMBLY (FUTURE) (Figure 9, panel(2)s)** | | |
| ATL | -290 - -285ms (max $\underline{t^{2}}$ = 5.02)  -265 - -245ms (max $\underline{t^{2}}$ = 6.13) | 340 - 345ms (max $\underline{t^{2}}$ = 4.36) |
| PTL |  | 290 - 295ms (max $\underline{t^{2}}$ = 3.91)  305 - 355ms (max $\underline{t^{2}}$ = 4.66)  380 - 385ms (max $\underline{t^{2}}$ = 4.11) |
| IPL |  | 320 - 325ms (max $\underline{t^{2}}$ = 3.76) |
| M1 | -30 - -15ms (max $\underline{t^{2}}$ = 4.69) | 5 - 20ms (max $\underline{t^{2}}$ = 4.04)  80 - 85ms (max $\underline{t^{2}}$ = 3.86) |
| **VERB INFLECTION + CONSTITUENT ASSEMBLY (PAST ) > VERB NAMING**  **(Figure 10, panel (1)s)** | | |
| ATL |  | 5 - 135ms (max $\underline{t^{2}}$ = 6.3) |
| PTL |  | 45 - 70ms (max $\underline{t^{2}}$ = 5.76)  80 - 105ms (max $\underline{t^{2}}$ = 5.47)  120 - 160ms (max $\underline{t^{2}}$ = 4.69) |
| IFG |  | 45 - 60ms (max $\underline{t^{2}}$ = 4.69)  70 - 75ms (max $\underline{t^{2}}$ = 4.32)  95 - 105ms (max $\underline{t^{2}}$ = 4.96) |
| **VERB NAMING > VERB INFLECTION + CONSTITUENT ASSEMBLY (PAST)**  **(Figure 10, panel(2)s)** | | |
| ATL | -260 - -255ms (max $\underline{t^{2}}$ = 5.91)  -235 - -195ms (max $\underline{t^{2}}$ = 7.3) |  |

**Supplementary Table S5.** Results of a two-stage regression analysis of MEG source-level responses, assessing the effect of number of syllables from selected ROIs (M1 - primary motor cortex, and IFG - inferior frontal gyrus).

| **LH Regions** | **800-1200 ms (condition icon)** | **1200 -1600 ms (speak icon)** | **>1600ms** |
| --- | --- | --- | --- |
| **Syllable effect** | | | |
| M1 | 945 - 960ms (max $\underline{t^{2}}$ = 4.4)  995 - 1000ms (max $\underline{t^{2}}$ = 4.16)  1105 - 1115ms (max $\underline{t^{2}}$ = 4.1)  1125 - 1140ms (max $\underline{t^{2}}$ = 4.52) | 1235 - 1245ms (max $\underline{t^{2}}$ = 4.45)  1365 - 1390ms (max $\underline{t^{2}}$ = 5.08) | 1610 - 1615ms (max $\underline{t^{2}}$ = 4.5)  1645 - 1655ms (max $\underline{t^{2}}$= 5.16) |
| IFG | 940 - 955ms (max $\underline{t^{2}}$ = 4.22)  990 - 1000ms (max $\underline{t^{2}}$ = 4.35)  1075 - 1085ms (max $\underline{t^{2}}$ = 5.28)  1120 - 1155ms (max $\underline{t^{2}}$ = 4.26) | 1200 - 1205ms (max $\underline{t^{2}}$ = 4.69)  1240 - 1250ms (max $\underline{t^{2}}$ = 5.33)  1290 - 1315ms (max $\underline{t^{2}}$ = 4.24)  1335 - 1345ms (max $\underline{t^{2}}$ = 4.05)  1355 - 1365ms (max $\underline{t^{2}}$ = 4.04)  1380 - 1385ms (max $\underline{t^{2}}$ = 4.74) |  |
